# Estimating Bayesian Phylogenetic Information Content Using Geodesic Distances

**DOI:** 10.64898/2026.03.31.715656

**Authors:** Analisa Milkey, Paul O. Lewis

## Abstract

A new Bayesian measure of phylogenetic information content is introduced based on geodesic distances in treespace. The measure is based on the relative variance of phylogenetic trees sampled from the posterior distribution compared to the prior distribution. This ratio is expected to equal 1 if there is no information in the data about phylogeny and 0 if there is complete information. Trees can be scaled to have the same mean tree length to avoid dominance by edge length information and focus on topological information. The method scales well, requiring only that a valid sample can be obtained from both prior and posterior distributions. We show how dissonance (information conflict) among data sets can also be estimated. Both simulated and empirical examples are provided to illustrate that the new approach produces sensible and intuitive results.

## Introduction

The canonical paraphrase of Claude Shannon’s concept of *information* in his 1948 foundational paper on information theory (Shannon, 1948) is that information is the resolution of uncertainty. The amount of information in data is often equated with the amount of data itself, but it is clearly true that a large quantity of data may contain little information (e.g. an entire book filled with randomly-generated text) and a small quantity of data may eliminate all uncertainty (e.g. the six words “I will have the Matar Paneer” uttered at a restaurant with hundreds of menu items). Systematists have long been concerned about how much information relevant to the problem of interest is available in the data they collect. Early work was concerned with determining the degree to which noise from homoplasy was affecting results (consistency index; Kluge and Farris, 1969) and how confident we should feel about different clades in a phylogenetic tree (bootstrapping; Felsenstein, 1985). Archie (1989) and Faith (1991) developed permutation methods designed to test whether there was a significant amount of historical information in data. Hillis and Huelsenbeck (1992) advocated using parsimony tree length skewness to assess the informativeness of data. This early work was followed by many papers addressing various aspects of information content in the data used by systematists to infer phylogenies (Steel et al., 1993, 1995; Lyons-Weiler et al., 1996; Goldman, 1998; Massingham and Goldman, 2000; Shpak and Churchill, 2000; Xia et al., 2003; Geuten et al., 2007; Townsend, 2007; Shi et al., 2008; Fischer and Steel, 2009; Lemey et al., 2009; San Mauro et al., 2009; Xia, 2009; Tippery et al, 2012; Townsend et al., 2012; Brown, 2014; Lewis et al., 2016; Duchêne et al., 2022).

The Bayesian statistical framework provides a reference distribution (the prior) for comparison with the posterior distribution. Information in data transforms the prior distribution into the posterior distribution, increasing the plausibility of some parameter combinations at the expense of other combinations. Comparing the posterior distribution to the prior distribution provides an explicit way to measure the information contained in the data (Lindley, 1956).

Lewis et al. (2016) proposed measures of phylogenetic information content and phylogenetic dissonance based on the relative entropy of a posterior distribution of tree topologies compared to the prior topology distribution. Common practice disperses prior probability mass evenly over every possible tree topology (i.e. maximum entropy). Zero topological **information** occurs when the posterior distribution exactly matches this maximum-entropy prior distribution. Complete information obtains when the entire posterior is concentrated over a single tree topology (i.e. minimum entropy).

Because information can be misinformation, it is important to also have ways of measuring conflict information among data sets. Phylogenetic **dissonance** was defined (Lewis et al., 2016) as the difference between the entropy of the merged posterior tree samples from two or more data subsets (e.g. loci) and the average posterior entropy within each subset. If all data subsets concentrate posterior mass in similar ways, then merging the sampled trees is expected to be simply a larger sample from the same distribution. If, however, some data sets disagree with others about which tree topologies are best, then the average entropy within datasets is expected to be much smaller than the entropy of the merged samples.

One of the drawbacks of the method described by Lewis et al. (2016) is scalability. The number of possible tree topologies quickly becomes too vast to sample adequately as the number of taxa increases. Consider a problem involving trees of only 12 taxa. Suppose 1 million tree topologies were sampled from the posterior distribution and every sampled tree topology was distinct from all other sampled topologies. It would seem that there is not much information about tree topology in the data because even a sample of size 1 million fails to find any tree topology that is more plausible than the others. Given that a flat prior distribution for unrooted trees of 12 taxa distributes probability evenly over all 654,729,075 possible tree topologies, the fact that the posterior sample is concentrated over a tiny fraction (0.001527) of the possible tree topologies suggests the opposite: that there is a tremendous amount of information in the data. In order to confidently conclude that there is no information in this data set, one would need to have a posterior sample size several times larger than the number of possible trees. While it is possible to ameliorate the problem by adjusting the maximum entropy to equal the log of the sample size rather than the log of the number of possible tree topologies, or by smoothing the empirical posterior probability distribution of tree topologies using conditional clade distributions (Lewis et al., 2016; Berling et al., 2025), this overestimation of information content due to the impossibility of adequately approximating the posterior distribution is a complication. The problem only gets much, much worse if the number of taxa is in the hundreds or thousands as opposed to 12.

This paper describes a very different approach to measuring phylogenetic information content that pays attention to edge lengths as well as topology and scales much better with larger numbers of taxa. In addition to measuring information content, we describe a corresponding measure of dissonance.

## Background

Brown and Owen (2020) discussed how to estimate the (Fréchet) mean and variance of a set of phylogenetic trees. In the context of Bayesian phylogenetics, the variance of a posterior sample of trees relative to the variance of a prior sample is useful as a measure of information content. Consider two mean trees computed from samples of trees from the posterior and prior distributions, respectively (Fig. 1). The data used for the posterior sample comprise 1296 DNA sites from the RuBisCO large subunit (*rbc*L) locus from 5 taxa chosen to span the green plant phylogeny. The prior and posterior samples were obtained using RevBayes version 1.3.1 (Höhna et al., 2016) (for details of settings, see the *1-fig-rbclmean* directory in the Supplementary Materials). The trees shown were determined using the Sturm (2003) algorithm as described in Miller, Owen, and Provan (2015) and Brown and Owen (2020). The posterior mean tree would make perfect sense to someone familiar with green plant evolution. For example, it places the two seed plants, *Picea* and *Avena*, as sister taxa in a tree rooted at the green alga *Chara* as well as placing all vascular plants (*Asplenium, Picea*, and *Avena*) in a clade. On the other hand, the prior mean is completely unresolved, which is expected because the prior treats every unrooted tree topology as equally probable. Note the relative scales of these trees. The posterior mean tree length is smaller than the prior mean tree length by an order of magnitude. Thus, the data have information about both the topology of the tree as well as the edge lengths.

**Figure 1:**
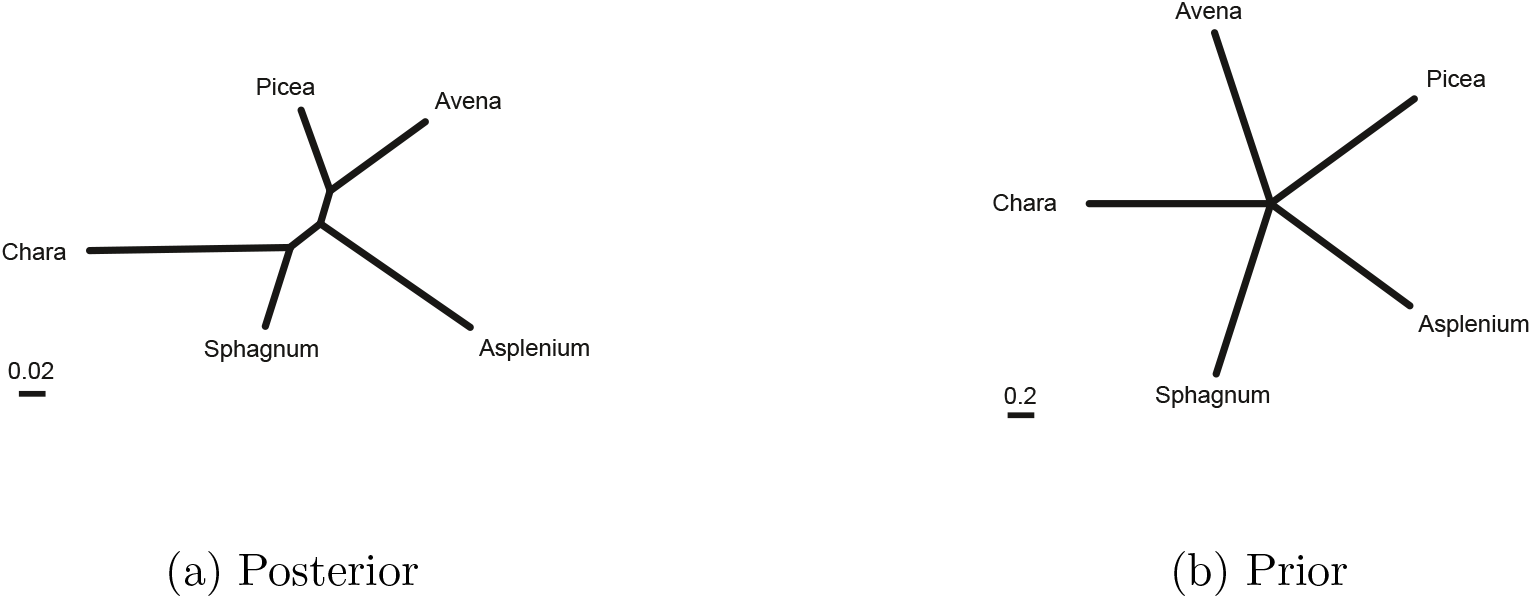
Fréchet mean trees computed from the (a) posterior and (b) prior samples using the *rbc*L dataset.

The mean tree is the tree minimizing the squared distance to all trees in a sample of phylogenetic trees. The variance of the sample is therefore obtained as a free byproduct of determining the mean tree. The distance used is the geodesic distance of Owen and Provan (2010) in the treespace characterized by Billera et al. (2001). For the example in Fig. 1, the variance of the posterior sample is 0.000728, while the variance of the prior sample (40.1) is 55 thousand times larger. In this paper, we propose using the difference between the prior and posterior sample phylogenetic variance as a measure of information content. A posterior variance equal to zero corresponds to complete information, whereas a posterior variance equal to the prior variance implies there is no information about the phylogeny in the observed data (although there may well be considerable information about other components of the model, such as nucleotide frequencies).

## Materials and Methods

### Information measure

Our method requires two samples of phylogenetic trees: a sample of size *N*_0_ from the prior distribution and a sample of size *N* from the posterior distribution. In this paper, we assume *N*_0_ = *N* and that both prior and posterior samples are obtained using MCMC. In RevBayes 1.3.1, for example, a sample from the prior may be obtained by calling the `ignoreAllData()` function of the model.

Shi et al. (2022) proposed the information content measure LCR (Log Concentration Ratio):

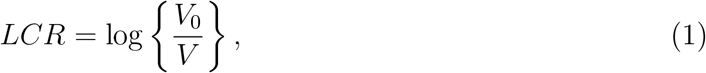

where *V*_0_ and *V* are (in Shi et al., 2022) the volumes of the 95% credible regions for the prior and posterior distributions, respectively. We equate volume (*V* and *V*_0_) with the 95% radius. To compute the 95% radius, a mean tree is obtained from a sample of trees, and the distance between that mean tree and each sampled tree is calculated. From these distances, the radius of the smallest hypersphere containing at least 95% of the distances from the mean tree is used as the “volume”. The danger of measuring volume in a single dimension is that information will be underestimated. For example, if the radius is reduced from 2 to 1, the volume is halved if assuming 1 dimension but is reduced by a factor of 4 if assuming even 2 dimensions. Nevertheless, we show that our 1-dimensional *LCR* behaves in a sensible and intuitive way for simulated data.

In addition to the 95% radius (hereafter denoted the RAD method), we explored two other ways of measuring dispersion:

- Letting *V* equal the standard deviation of distances from the mean tree (the STD method) and
- Letting *V* equal the radius associated with the 95% highest posterior density credible set of trees (the HPD method)

The HPD method results in a larger radius if the posterior is asymmetric (Fig. S1, Supplementary Materials). We found (Figs. S2 and S3, Supplementary Materials) that, while both alternative ways of measuring information largely agree with information measured using the RAD method, we prefer the RAD method because it is both more stable (avoiding the undue influence of occasional outliers that the STD approach would include) and more intuitive (avoiding most of the cases of negative information content that are often produced by the HPD method when information content is low). Ideally, *V* would equal the actual *volume* of treespace (as in the original LCR formulation by Shi et al. (2022)) but, unfortunately, treespace is only partially Euclidean and, lacking any way to accurately measure the volume of treespace regions, we default to the 95% radius as our measure of dispersion in this paper.

Some may find *LCR* difficult to interpret given the log scale. Transforming *LCR* to percent information (*I*) may be preferable:

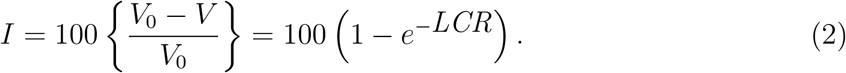

Whereas *LCR* ranges from 0 (no information^1^) to ∞ (complete information), *I* ranges from 0 (no information) to 100 (complete information) and thus has a similar interpretation to the measure of Lewis et al. (2016).

#### Scaling to a common tree length

Under normal circumstances, we expect *V* to be considerably smaller than *V*_0_, reflecting the fact that sequence data generally contains information relevant to both tree topology and edge lengths. Information content measured by *LCR* is often dominated by the difference in tree length between the posterior and prior sample. One way to reduce the influence of tree length on information content, giving the topological information more influence, is to scale trees in both prior and posterior samples such that the prior mean tree and the posterior mean tree both have a common tree length. This rescaling does not mean that the only information measured is topological; the data are expected to induce correlations among edge lengths in the posterior sample that are not present in the prior sample. Such correlations affect the posterior variance even though the prior sample is scaled to match the posterior mean tree length. Unless explicitly stated, information content using equation (1) scales sampled trees so that the prior mean tree and posterior mean tree each have the same total length of 1.0.

Both the Fréchet mean tree and tree distances were calculated in this paper using the open-source software op, available from https://github.com/plewis/op. For all analyses in this paper, we instructed the op program to stop when the maximum pairwise distance among the mean trees produced by the *N* = 10 most recent iterations first dropped to less than *ϵ* = 0.00001.

### Dissonance measure

Dissonance between two posterior distributions of trees is measured as a modified effect size. Let *d*_12_ be the geodesic distance between the scaled mean trees of the two data sets. Each mean tree is scaled to have tree length 1.0. Let *n*_1_ and *n*_2_ be the sample sizes and 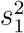 and 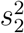 the scaled sample variances of the two posterior distributions, respectively. Variances are scaled using the same scaling factor used to scale mean trees: if the mean tree length for data set 1 is *L*, the mean tree is scaled by dividing each edge by *L* and the variance is scaled by dividing the unscaled variance by *L*^2^. The effect size may be computed as Cohen’s d:

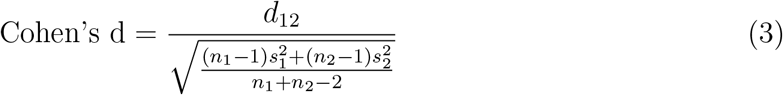

For compatibility with our information measure, we use a modified effect size that substitutes the 95% radius for *s*_1_ and *s*_2_. The formula for dissonance (*D*) is thus

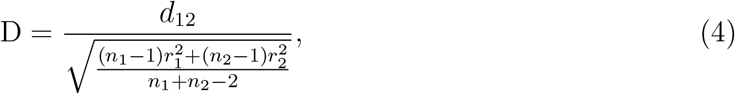

where *d*_12_ is the geodesic distance between the two (scaled) mean trees, and *r*_1_ and *r*_2_ are the 95% radii for the two (scaled) posterior samples.

Dissonance is expected to be zero if the two posterior distributions compared are identical (and thus have identical mean trees), and is expected to grow with increasing distance between the mean trees. Note that two data sets simulated independently from identical model trees are *not* expected to exhibit zero dissonance. Even though conditionally independent, the two data sets are not identical and thus have distinct posterior distributions.

### Simulation experiments

#### Information

We conducted a series of simulation experiments to characterize whether *LCR* performs as expected for differing substitution rates, sequence lengths, proportion missing data, and rate heterogeneity. Information content is expected to be low if the substitution rate is near zero. At the extreme of zero substitution rate, all sites would be constant and the data would therefore provide no evidence relevant for identifying historical groupings of taxa. At the ideal substitution rate, information content is expected to be 100% given that sequences are of sufficient length to provide complete evidence for every split. Increasing the substitution rate much beyond this ideal rate should result in a decrease in information because sequences eventually become saturated with substitutions (newer substitutions overwrite older substitutions that provided evidence of history). High among-site rate heterogeneity (ASRV) should also result in decreased phylogenetic information content, even if the mean substitution rate is ideal, because, with high ASRV, most sites are evolving at a rate less than the mean rate, and the remaining sites evolve at a high rate. Finally, a high proportion of missing data (missing-at-random) should decrease information content, and information should decrease with decreasing number of sites.

We measured information content on simulated DNA sequence data sets with 5 taxa and variable sequence lengths, mean substitution rates, amount of ASRV, and percent missing data. The tree used as the model tree for these simulations is that shown in Fig. 1a. DNA sequences on this tree were generated using Seq-Gen 1.3.4 (Rambaut and Grassly, 1997) and analyzed using RevBayes 1.3.1 using 50,000 iterations following a 2000-iteration burn-in period and a GTR substitution model with Gamma-Dirichlet tree length/edge proportion prior (Zhang et al., 2012). Trees were sampled every 5 iterations, yielding 10,000 sampled trees. Scripts for all analyses are provided in the supplementary materials (directory *2-fig-lcr*).

#### Dissonance

To evaluate dissonance, we conducted a simulation experiment in which 20 starting trees, each of 26 taxa, were drawn from the prior and moved via a random walk through treespace. Each random walk involved 500 steps and each step consisted of adding a Normal variate with mean zero and standard deviation 0.005 to each edge length. Pendant (leaf) edges that dropped below 1 × 10^−12^ were reflected back into the interval [1 × 10^−12^, ∞). Internal edges that dropped below zero resulted in a change in topology; one of the two possible alternative splits was chosen uniformly at random and received the residual edge length. Six trees were sampled at uniformly spaced points (stations) along this path (i.e. stations 1, 2, 3, 4, 5, 6 used trees sampled from steps 0, 100, 200, 300, 400, and 500). For each of the 6 stations along the path, for each of the 20 replicates, two data sets were simulated using Seq-Gen and posterior samples were obtained using RevBayes 1.3.1. The first simulated data set corresponded to the starting tree for that replicate (i.e. station 1); the second data set corresponded to the tree sampled at that particular station. The first station thus produced two replicate simulated data sets using the same model tree. Dissonance was measured for each pair of data sets at each of the 6 stations for all 20 replicates. The expectation is that dissonance will grow with increasing distance between model trees; however, as mentioned previously, we do not expect the dissonance between the two data sets simulated from the same model tree at station 1 to be zero because the data sets, while conditionally independent, are not identical. Scripts for all analyses are provided in the supplementary materials (directory *3-fig-dissonance*)

### Empirical analyses

#### Saturation

It is often assumed *a priori* that 3rd codon position sites in protein coding genes are saturated. Lewis et al. (2016) estimated topological information content in 2nd vs. 3rd position sites in the *psa*B locus in Spheroplealean green algae and found that the 3rd position sites were in fact not saturated and contained more information than 2nd position sites. We reanalyzed these data using equation (1). We also applied the saturation test in PhyloMAd (Duchêne et al., 2018) as an independent check for saturation. Scripts for all analyses are provided in the supplementary materials (directory *4-fig-saturation*)

#### Dissonance

We re-analyzed an example from Lewis et al. (2016) in which high dissonance is expected. The 5’ half of the mitochondrial *rps11* locus in Bloodroot (*Sanguinaria*, Papaveraceae) is vertically transferred, whereas the 3’ half was evidently horizontally transferred from a phylogenetically-distant monocot (Bergthorsson et al., 2003). We reanalyzed these data using equation (3). Scripts for all analyses are provided in the supplementary materials (directory *5-fig-bloodroot*)

## Results

### Simulation experiments

#### Information

Information content was highest when the relative substitution rate was 1 (i.e. the edge lengths used for simulating data were equal to those in Fig. 1a) and decreased when the data were simulated with either larger or smaller relative rates (Fig. 2a). Information content was zero when the number of sites was zero and generally increased as the sequence length increased, with the exception of the 1-site case, which, on average, resulted in slightly negative information (Fig. 2b). Information content decreased as the percentage of missing nucleotides grew from 10% to 95% (Fig. 2c). Finally, information content decreased with increasing variance in substitution rates across sites (Fig. 2d).

**Figure 2:**
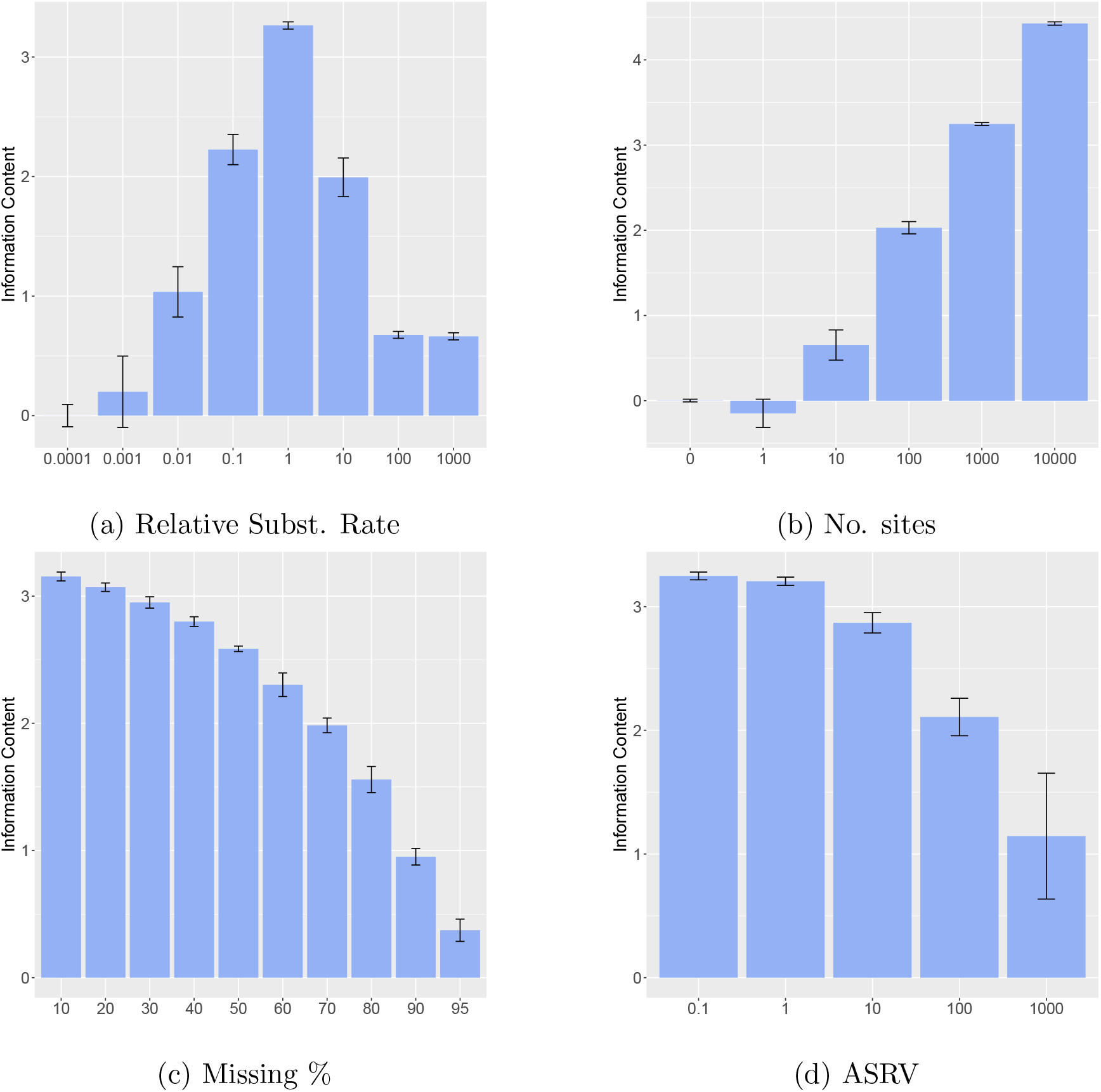
Information content (*LCR*) for 5-taxon datasets. a. Rate is relative to the edge lengths in the model tree. b. No. sites is the sequence length. c. Missing % is the percent of nucleotides missing at random. d. ASRV is measured as the variance in among-site relative rates. In each case, rate=1.0, ASRV=0.0, no. sites=1000, and missing = 0% except for the variable on the x-axis. Error bars show mean ± standard deviation across *n* = 10 replicates.

#### Dissonance

For random walks beginning at trees drawn from the prior, the distance between trees increased as a function of the number of random walk steps taken (Fig. 3a). The mean geodesic distance between starting and ending trees was 0.606209 (std. dev. 0.066439, *n* = 20). As expected, the dissonance between two data sets is strongly and positively correlated with the geodesic distance between the trees used to generate the data (Fig. 3b). Only 4 interstation segments (of 5 segments × 20 replicates = 100 total) failed to show increased dissonance. Dissonance was, on average, 0.124232 (std. dev. 0.011204, *n* = 20) when both data sets were simulated from the same tree and increased with increasing distance between model trees. A dissonance of 0.12 means that the distance between means represents 12% of the pooled radius. The total dissonance between the starting and ending data sets was 0.734085 (std. dev. 0.083814, *n* = 20). Thus, the random walks were not long enough to completely separate the starting and ending posterior distributions. There is a strong, positive correlation (0.96, *n* = 120, *P <* 0.0001) between the distance separating model trees and the dissonance of the corresponding posterior distributions.

**Figure 3:**
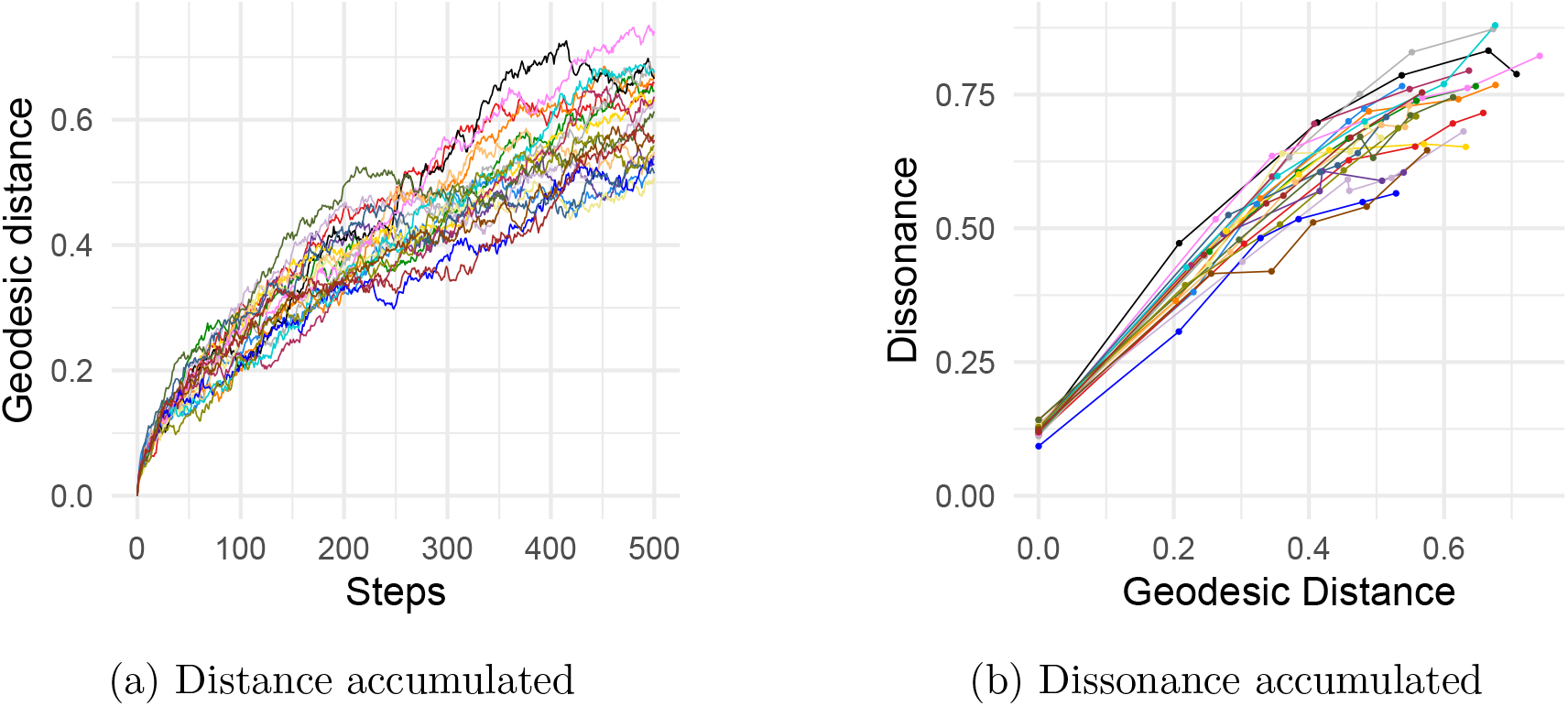
(a) Geodesic distance accumulated during random walk. (b) Dissonance as a function of geodesic distance.

### Empirical analyses

#### Saturation

Like Lewis et al. (2016), we found that 3rd position sites at the *psa*B locus in the green algal data set contained more information (*LCR* = 2.73, *I* = 93.5) than 2nd position sites (*LCR* = 1.75, *I* = 82.6). The Fréchet mean trees computed from posterior samples from the two sets of sites clearly show greater resolution in the 3rd-position sites (Fig. 4), which reflects the greater amount of phylogenetic information in 3rd-position sites.

**Figure 4:**
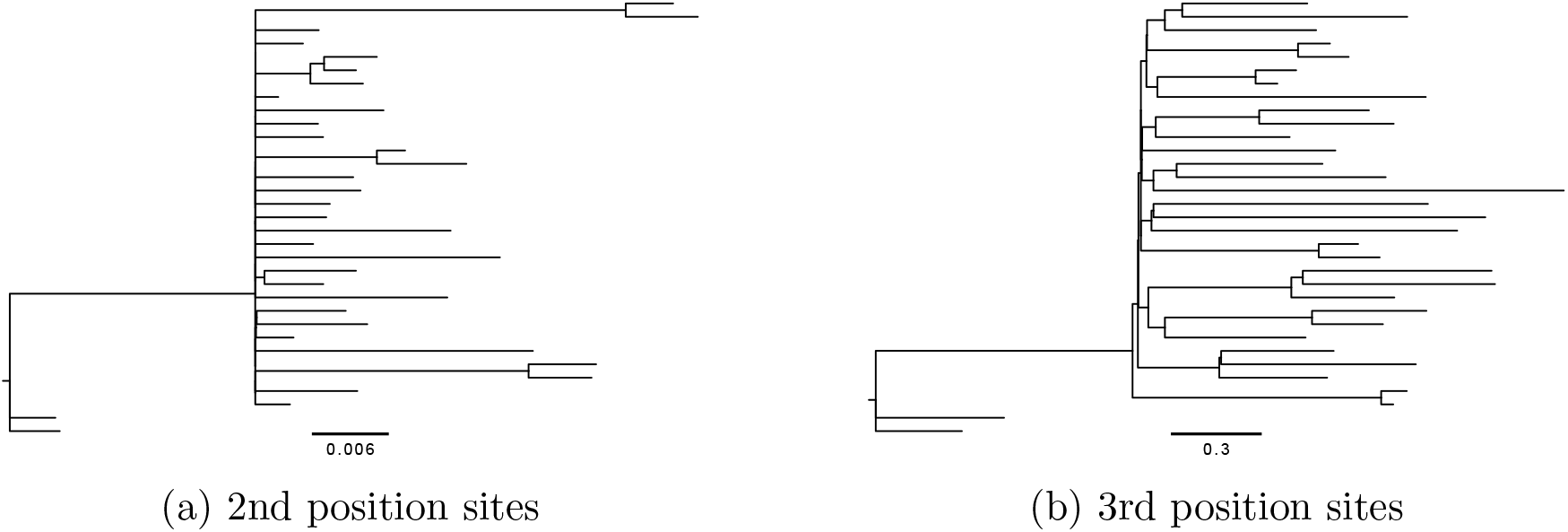
Fréchet mean trees computed from the *psa*B locus: (a) 2nd-codon-position sites only, and (b) 3rd-codon-position sites only.

3rd-position sites also had more information than 1st-position sites (*LCR* = 2.36, *I* = 90.5) and even slightly more information than 1st and 2nd positions combined (*LCR* = 2.58, *I* = 92.5). As expected, combining sites from all three codon positions yielded the most information (*LCR* = 3.48, *I* = 96.9).

A natural question is “Is the information in 3rd-position sites truthful information or is it misinformation?” Using either BHV distances (BHV; Owen and Provan, 2010), or Robinson-Foulds distances(RF; Robinson and Foulds, 1981), the 3rd-position mean tree is closer to the all-sites mean tree than is the 2nd-position mean tree (Table 1). This may mean only that the 3rd-position sites, with their greater information content, dominate in analyses using all sites. Note, however, that the 3rd-position mean tree is also closer to the 1st-position mean tree than is the 2nd-position mean tree. These results are contradicted by Cluster Information Distance (CID; Smith, 2020), which is arguably a more sensitive topology-only tree distance than the Robinson-Foulds distance. At the very least, given the small differences between these distances, it would be difficult to argue that either 2nd-position or 3rd-position sites are saying something completely different than 1st-position sites.

**Table 1:** BHV distances (BHV), Cluster Information Distances (CID), and Robinson-Foulds Distances (RFD) between mean trees from Bayesian analyses of codon-based data partitions. Each distance is a simple average of 4 distances, one from each of the two independent runs.

| Comparison | BHV | CID | RF |
| --- | --- | --- | --- |
| 2nd vs 1st | 0.09228 | 0.47349 | 35.5 |
| 3rd vs 1st | 0.09080 | 0.48355 | 33.5 |
| 2nd vs all | 0.10893 | 0.35122 | 29 |
| 3rd vs all | 0.05265 | 0.36291 | 25 |

The saturation test in PhyloMAd concluded that all data subsets (1st-codon, 2nd-codon, and 3rd-codon, as well as the entire locus), for both empirical and all simulated data sets, were ‘low risk’. This test computes a t-statistic that measures how surprised we should be at the site patterns in the data if the data subset was saturated (i.e. observed nucleotides are distributed according to the global empirical nucleotide frequencies). The critical value of the t-statistic is determined not from the t distribution, but, instead, is the value that maximizes the ratio of true positives to false positives from simulations covering a diversity of parameter combinations. The test is thus conservative, not requiring the data to be truly saturated to be declared unsound, yet in all cases the test said that the data were not close to being saturated, confirming our conclusions based on LCR.

#### Dissonance

The mean tree from a sample of 10,000 trees from the posterior distribution of the 5’ subset of *rps11* shows *Sanguinaria* properly associated with other genera (*Bocconia* and *Eschscholzia*) in its eudicot family (Poppy Family, Papaveraceae), whereas the 3’ posterior mean tree shows *Sanguinaria* associated with unrelated monocot genera (*Oryza* and *Disporum*) (Fig. 5).

**Figure 5:**
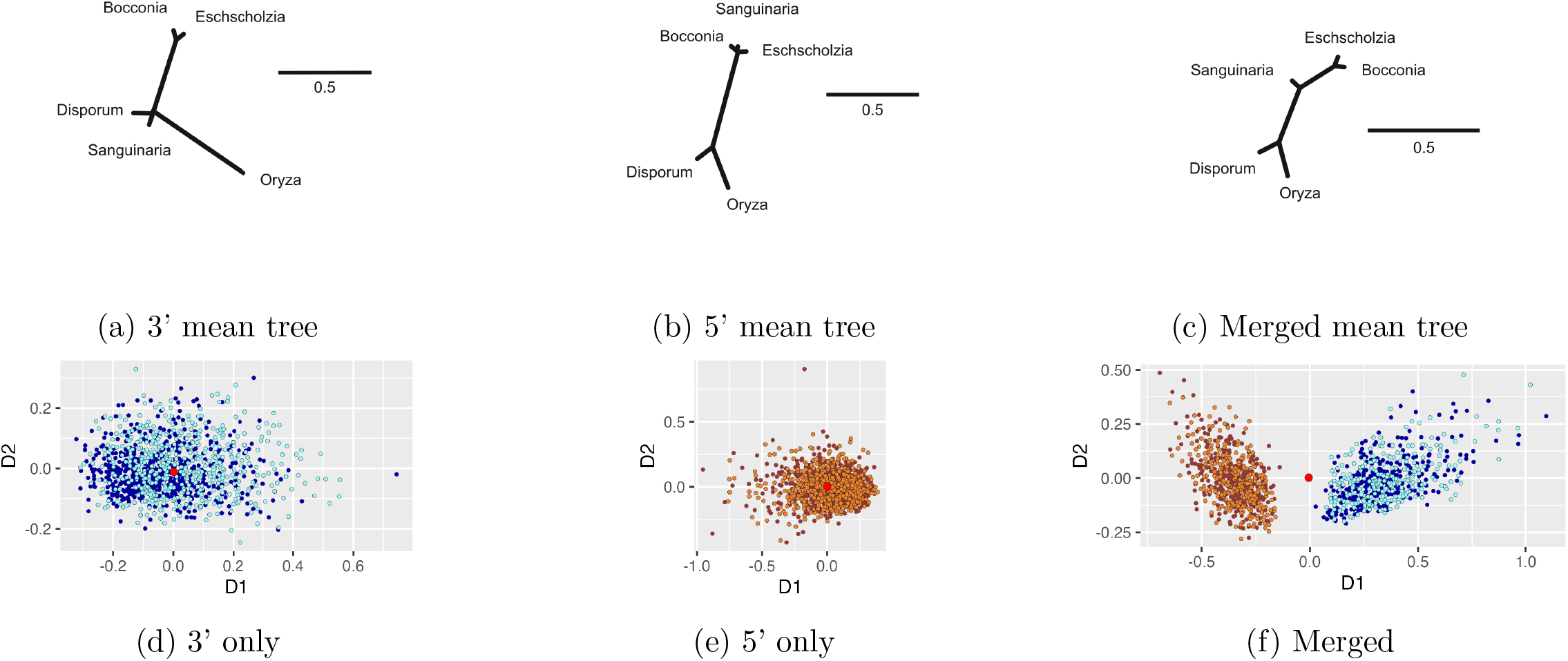
Comparisons of posterior samples from subsets of the *rps11* locus. (a-c) Mean trees from posterior samples conditioned on the 3’ subset, the 5’ subset, and the merged sample, respectively. (d-e) Multidimensional scaling (MDS) comparisons of 2 random subsamples, each of size 200, from independent analyses of the same 3’ and 5’ posterior distributions, respectively. (f) MDS comparison of merger of the 4 subsamples in (d) and (e). Larger red point in each plot is the mean tree.

Two independent MCMC analyses were conducted of both 3’ and 5’ data subsets, allowing comparison of dissonance measured between independent samples from the same posterior (i.e. 3’ vs. 3’ or 5’ vs. 5’) to dissonance measured between samples from different posteriors (e.g. 3’ vs 5’). Dissonance was consistently greater than 8 for comparisons between the 3’ and 5’ subsets but less than 0.2 for comparisons of independent samples from the same posterior distribution (Table 2).

**Table 2:**
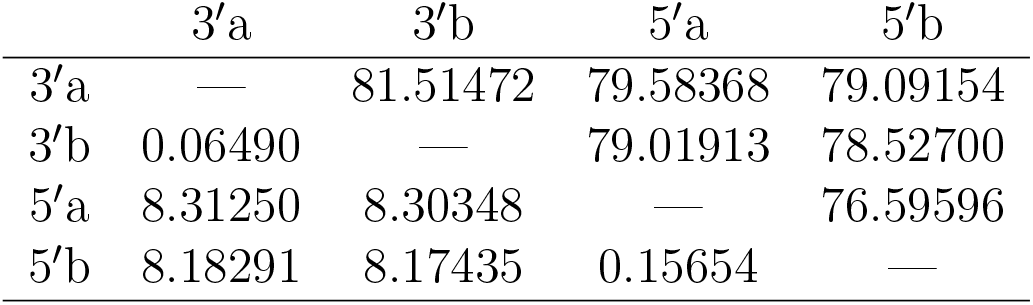
Dissonance between posterior samples (*D*; lower triangle) and average information (*I*; upper triangle). a and b denote independent samples of the same posterior distribution.

## Discussion

The methods presented in this paper use the difference in phylogenetic variance between prior and posterior tree samples to make inferences about the phylogenetic information content of data. This builds on previous work by Lewis et al. (2016), who measured information content using the difference between prior and posterior entropy in discrete topological probability distributions. The primary drawback of the Lewis et al. (2016) method was scalability. The methods we present here are very different, making use of recent algorithms for efficiently determining geodesic distances between trees to estimate the mean and variance of phylogenetic trees (Owen and Provan, 2010; Brown and Owen, 2020; Miller, Owen, and Provan, 2015). The new measure yields similar inferences of information content as the entropy-based measures if the prior sample is scaled to have the same mean tree length as the posterior sample. They have the advantage of being at least as scalable as the Bayesian sampling itself; that is, if it is possible to obtain a good sample from the posterior and prior distributions, then it is possible to apply our method.

In addition to measuring information content, we have also introduced a geodesic-distance-based measure of dissonance that is large and positive if data subsets conflict in their choice of tree and small if data subsets do not conflict.

One nice feature of the approach described by Lewis et al. (2016) is that information can be partitioned. For example, it is possible to say that 90% of the total information is attributable to a single well-supported clade. This additivity of information content is not currently possible with the method described here, although future work may reveal ways to assess the relative contribution of different subsets of taxa to the total information content.

More work is also needed to determine the best definition of “volume” to use in the LCR measure. Here, we’ve used as volume the 95% radius: that is, the distance from the mean tree to the most distant sampled tree that nevertheless lies within the 95% set of trees closest to the mean tree. It would be more appropriate to use a measure of the true volume of treespace enclosed by the 95% set of sampled trees closest to the mean, but the non-Euclidean component of tree space as defined by Billera et al. (2001) places this definition of volume currently out of reach.

One clear application of our information content measure is in phylogenomics. When data from many loci are available, it is not always advisable to include all loci (Duchêne et al., 2022). In particular, species tree methods such as BEAST2 (Bouckaert et al., 2019) that jointly estimate gene trees and the species tree could be made more computationally efficient by filtering out loci that have very little information to contribute. Species tree inference using ASTRAL, which treats gene trees as observed data, could be improved by using the mean tree from each locus as input. The mean tree is expected to be poorly resolved if information content is low, but the clades that are present in the mean tree are those that are supported by the information in the data for that locus. Thus, using the mean tree as input to a species tree method would be less likely to contribute false information than using the maximum likelihood or maximum *a posteriori* (MAP) tree and provides an alternative to collapsing nodes in gene trees using arbitrary rules.

With extremely fast tests for saturation available in software such as DAMBE (Xia, 2009) and PhyloMAd (Duchêne et al., 2018), why use the method described here, which is itself fast but requires a valid Bayesian posterior sample that may take hours or days to obtain? One reason is that saturation tests test only one end of the information spectrum: they do not reveal loci that exhibit low information due to a paucity of substitutions. Another reason is that our approach makes use of the exact Bayesian model being used for inference, whereas entropy t-tests such as those in PhyloMAd use critical values predicted from simulation experiments that, while exploring many relevant parameter combinations, cannot anticipate the behavior of the specific model being used. In particular, PhyloMAd may have difficulty accurately identifying problematic data sets when the model used is complex (Duchêne et al., 2022), for example, the Bayesian CAT model (Lartillot and Philippe, 2004). Finally, our approach directly measures the information content of data by comparing the posterior distribution, which is informed by the data, to the prior distribution, which serves as a reference point not influenced by the observed data.

## Acknowledgments

We would like to thank xxxxx for their constructive comments.

The computational work for this project was conducted using resources provided by the Storrs High-Performance Computing (HPC) cluster. We extend our gratitude to the UConn Storrs HPC and its team for their resources and support, which aided in achieving these results.

## Supplementary Material

Data available from the Dryad Digital Repository: http://datadryad.org/xxxxx

## Footnotes

1 but can be negative if the posterior has higher variance than the prior

