## Supplementary material for "Estimating Bayesian Phylogenetic Information Content Using Geodesic Distances": _readme.txt

To create fig-rbclmean:

Directory layout showing only directories and files mentioned in the instructions below:

  supplementary-materials
      calcinfo.py 
      1-fig-rbclmean
          data
            green5.nex
*         op-results
*           op-post.tre
*           op-prior.tre
*         rbcl
*           posterior
*           prior
          _readme.txt
          fig-rbclmean.Rev
*         op-results-output.txt
          rbcl-slurm.sh
    
* file/directory created as a result of these instructions

1. Edit the rbcl-slurm.sh and adapt the SBATCH commands at the top 
    to your cluster. Be sure the paths to the RevBayes and OP
    executables are correct.

    If you do not have these programs installed, here is where you can get them:
        RevBayes 1.3.1:   https://revbayes.github.io/download
        OP 1.9.4:         https://github.com/plewis/op/
                
2. Specify "hpc" as an alias to your remote cluster in  ~/.ssh/config and
   create a "work/rbclmean" directory inside your home directory on your cluster
   
   ssh hpc
   mkdir -p work/rbclmean
   mkdir -p work/rbclmean/data
    
3. Upload files to the "work" and "work/rbclmean" directories on your cluster:

    cd supplementary-materials
    scp calcinfo.py           hpc:work
    cd 1-fig-rbclmeans
    scp rbcl-slurm.sh         hpc:work/rbclmean
    scp fig-rbclmean.Rev      hpc:work/rbclmean
    scp data/green5.nex       hpc:work/rbclmean/data
    
4. Run the RevBayes and OP analyses on the cluster:

    ssh hpc
    cd work/rbclmean
    sbatch rbcl-slurm.sh

   This will run RevBayes and create a directory "rbcl" with subdirectories
   "prior" and "posterior" containing MCMC samples. It will also run 
   calcinfo.py, which runs OP to calculate information content
    
5. Bring results back to your local computer:

    cd supplementary-materials/1-fig-rbclmean
    rsync -azPv hpc:work/rbclmean/ ./

The op-results-output.txt file will contain estimated information as well as statistics reported in the paper such as the posterior and prior variance.

The op-results/op-post.tre file will contain the mean tree for the 
posterior sample.

The op-results/op-prior.tre file will contain the mean tree for the
prior sample.

Both trees can be visualized and saved as png files using FigTree.
