## Supplementary material for "Estimating Bayesian Phylogenetic Information Content Using Geodesic Distances": rbcl-slurm.sh

```
#!/bin/bash
#SBATCH --job-name=rbcl                         # job name
#SBATCH --partition=general                     # Name of Partition
#SBATCH --ntasks=1                              # Maximum CPU cores for each job
#SBATCH --nodes=1                               # Ensure all cores are from the same node for each job
#SBATCH --cpus-per-task=1                       # cpu-cores per task (>1 if multi-threaded tasks) for each job
#SBATCH --constraint='epyc128'                  # Target the AMD Epyc 128-core node architecture
#SBATCH --mem=20G                               # Request 20GB of available RAM
#SBATCH --mail-type=ALL                         # Event(s) that triggers email notification (BEGIN,END,FAIL,ALL)
#SBATCH --mail-user=        # Destination email address

# Set master random number seed
RANDOM=45971

# Specify paths to programs used
basedir=`pwd`
RB=$HOME/bin/rb131
OP=$HOME/bin/op194

# Load modules needed
module load python/3.12.5

# Generate seed for each analysis
postseed=("$RANDOM")
priorseed=("$RANDOM")

echo "RevBayes version:"
$RB -v

echo
echo "********************************"
echo "***** Creating directories *****"
echo "********************************"
# Create the main directory
mkdir -p $basedir/rbcl

# Create the posterior directory
mkdir -p $basedir/rbcl/posterior
echo "rnseed <- $postseed; run_under_prior <- FALSE" > rbcl/posterior/adhoc.Rev

# Create the prior directory
mkdir -p $basedir/rbcl/prior
echo "rnseed <- $priorseed; run_under_prior <- TRUE" > rbcl/prior/adhoc.Rev

echo
echo "****************************************"
echo "***** Running RevBayes (posterior) *****"
echo "****************************************"
cd $basedir/rbcl/posterior
$RB ../../fig-rbclmean.Rev

echo
echo "************************************"
echo "***** Running RevBayes (prior) *****"
echo "************************************"
cd $basedir/rbcl/prior
$RB ../../fig-rbclmean.Rev

echo
echo "****************************************"
echo "***** Running OP to calculate info *****"
echo "****************************************"
cd $basedir
python3 ../calcinfo.py --oppath $OP --dirname . --postfile $basedir/rbcl/posterior/output/algae.trees --priorfile $basedir/rbcl/prior/output/algae.trees --epsilon 0.00001
```
