## Supplementary material for "Estimating Bayesian Phylogenetic Information Content Using Geodesic Distances": _readme.txt

To create fig-lcr:

Directory layout showing only directories and files mentioned in the instructions below:

  supplementary-materials
      calcinfo.py
      2-fig-lcr
*         plots
*         results
*             argv
*             missing
*             nsites
*             rates
          _readme.txt
          addmissing.py
*         asrv-rbop-slurm.sh
          fig-lcr.Rev
          make-asrv-rbop-slurm.py
          make-missing-rbop-slurm.py
          make-nsites-rbop-slurm.py
          make-rates-rbop-slurm.py
*         missing-rbop-slurm.sh
          modeltree.txt
*         nsites-rbop-slurm.sh
*         rates-rbop-slurm.sh
          slurm-asrv-rbop-template.sh
          slurm-missing-rbop-template.sh
          slurm-nsites-rbop-template.sh
          slurm-rates-rbop-template.sh
          summarize.py        

* file/directory created as a result of these instructions

Note: The tree in modeltree.txt is identical to supplementary-materials/1-fig-rbclmean/op-results/op-post.tre
and supplementary-materials/1-fig-rbclmean/op-results/op-post.R

1. Edit make-rates-rbop-slurm.py and specify any changes you wish to make to these variables:

    __USEREMAIL__
    __NREPS__
    __MASTERSEED__

2. Edit the slurm-rates-rbop-template.sh and adapt the SBATCH commands at the top 
    to your cluster. Be sure the paths to the Seq-Gen and RevBayes 
    executables are correct.

    If you do not have these programs installed, here is where you can get them:
        Seq-Gen:    https://github.com/rambaut/Seq-Gen/releases/tag/1.3.4
        RevBayes:   https://revbayes.github.io/download

3. To create the rates-rbop-slurm.sh script for the analyses reported in Fig. 2a:

    python3 make-rates-rbop-slurm.py

4. Repeat steps 1 and 2 for make-nsites-rbop-slurm.py and slurm-nsites-rbop-template.sh 
   to create the nsites-rbop-slurm.sh script for the analyses reported in Fig. 2b:

    python3 make-nsites-rbop-slurm.py

5. Repeat steps 1 and 2 for make-missing-rbop-slurm.py and slurm-missing-rbop-template.sh 
   to create the missing-rbop-slurm.sh script for the analyses reported in Fig. 2c:

    python3 make-missing-rbop-slurm.py

6. Repeat steps 1 and 2 for make-asrv-rbop-slurm.py and slurm-asrv-rbop-template.sh 
   to create the asrv-rbop-slurm.sh script for the analyses reported in Fig. 2d:

    python3 make-asrv-rbop-slurm.py
    
7. Specify "hpc" as an alias to your remote cluster in  ~/.ssh/config and
   create a "work/lcrrad" directory inside your home directory on your cluster
   
   ssh hpc
   mkdir -p work/lcr
    
8. Upload files to the "work" and "work/sim" directories on your cluster:

    cd supplementary-materials/2-fig-lcr
    scp ../calcinfo.py        hpc:work
    scp rates-rbop-slurm.sh   hpc:work/lcr
    scp nsites-rbop-slurm.sh  hpc:work/lcr
    scp missing-rbop-slurm.sh hpc:work/lcr
    scp asrv-rbop-slurm.sh    hpc:work/lcr
    scp modeltree.txt         hpc:work/lcr
    scp fig-lcr.Rev           hpc:work/lcr
    scp addmissing.py         hpc:work/lcr
    
9. Run the analyses on the cluster:

    ssh hpc
    cd work/lcr
    sbatch rates-rbop-slurm.sh
    sbatch nsites-rbop-slurm.sh
    sbatch missing-rbop-slurm.sh
    sbatch asrv-rbop-slurm.sh
    
10. Bring results back to your local computer:

    cd supplementary-materials/2-fig-lcr
    mkdir results
    cd results
    rsync -azPv hpc:work/lcr/rates .
    rsync -azPv hpc:work/lcr/nsites .
    rsync -azPv hpc:work/lcr/missing .
    rsync -azPv hpc:work/lcr/asrv .

11. Summarize results (assume summarize.py is in parent directory of results):

    python3 summarize.py --dir results --categ rates
    python3 summarize.py --dir results --categ nsites
    python3 summarize.py --dir results --categ missing
    python3 summarize.py --dir results --categ asrv
    
    Files representing Figure 2a,b,c,d should now be in the "plots" directory.
