## Supplementary material for "Estimating Bayesian Phylogenetic Information Content Using Geodesic Distances": addmissing.py

```
import re,os,sys,random

narg = len(sys.argv)
assert narg == 5, 'Usage: python3 add-missing.py <input nexus file name> <output nexus file name> <missing percentage> <pseudorandom number seed>'
infilename = sys.argv[1]
outfilename = sys.argv[2]
missing_percent = float(sys.argv[3])
rnseed = int(sys.argv[4])

def readNexusFile(fn):
    '''
    Reads nexus file whose name is specified by fn and returns ntax, nchar, taxa, and a
    sequences dictionary with taxon names as keys. The values ntax and nchar are integers,
    while taxa is a list of taxon names in the order they were found in the taxa block or
    data block. Any underscores in taxon names are converted to spaces before being saved
    in the taxa list or as a key in the sequences dictionary. Also all nexus comments
    (text in square brackets) will be ignored.
    '''
    stuff = open(fn, 'r').read()
    mask = None

    # determine if taxa block exists
    taxa_block = None
    m = re.search(r'(?:BEGIN|Begin|begin)\s+(?:TAXA|Taxa|taxa)\s*;(.+?)(?:END|End|end)\s*;', stuff, re.M | re.S)
    if m is not None:
        taxa_block = m.group(1).strip()

    # determine if characters block exists
    characters_block = None
    m = re.search(r'(?:BEGIN|Begin|begin)\s+(?:CHARACTERS|Characters|characters)\s*;(.+?)(?:END|End|end)\s*;', stuff, re.M | re.S)
    if m is not None:
        characters_block = m.group(1).strip()

    # determine if data block exists
    data_block = None
    m = re.search(r'(?:BEGIN|Begin|begin)\s+(?:DATA|Data|data)\s*;(.+?)(?:END|End|end)\s*;', stuff, re.M | re.S)
    if m is not None:
        data_block = m.group(1).strip()

    if data_block is not None:
        # get ntax and nchar
        m = re.search(r'(?:DIMENSIONS|dimensions|Dimensions)\s+(?:NTAX|ntax|Ntax|NTax)\s*=\s*(\d+)\s+(?:NCHAR|nchar|Nchar|NChar)\s*=\s*(\d+)\s*;', data_block, re.M | re.S)
        assert m, 'Could not decipher dimensions statement in data block'
        ntax = int(m.group(1))
        nchar = int(m.group(2))

        # get matrix
        m = re.search(r'(?:MATRIX|matrix|Matrix)\s+(.+?)\s*;', data_block, re.M | re.S)
        assert m, 'Could not decipher matrix statement in data block'
        lines = m.group(1).strip().split('\n')
        taxa = []
        sequences = {}
        for line in lines:
            m = re.match(r'\[([-*]+)\]', line.strip())
            if m is not None:
                mask = m.group(1)
            else:
                stripped_line = re.sub(r'\[.+?\]', '', line).strip()
                if len(stripped_line) > 0:
                    parts = line.split()
                    assert len(parts) == 2, 'Found more than 2 parts to this line:\n%s' % line
                    taxon_name = re.sub(r'_', ' ', parts[0]).strip()
                    taxa.append(taxon_name)
                    sequences[taxon_name] = parts[1]
    else:
        assert characters_block is not None and taxa_block is not None, 'Assuming nexus file contains either a data block or a taxa block and characters block'

        # get ntax from taxa block
        m = re.search(r'(?:DIMENSIONS|dimensions|Dimensions)\s+(?:NTAX|ntax|Ntax|NTax)\s*=\s*(\d+)\s*;', taxa_block, re.M | re.S)
        assert m, 'Could not decipher dimensions statement in taxa block'
        ntax = int(m.group(1))

        # get nchar from characters block
        m = re.search(r'(?:DIMENSIONS|dimensions|Dimensions)\s+(?:NCHAR|nchar|Nchar|NChar)\s*=\s*(\d+)\s*;', characters_block, re.M | re.S)
        assert m, 'Could not decipher dimensions statement in characters block'
        nchar = int(m.group(1))

        # get matrix from characters block
        m = re.search(r'(?:MATRIX|matrix|Matrix)\s+(.+?)\s*;', characters_block, re.M | re.S)
        assert m, 'Could not decipher matrix statement in characters block'
        lines = m.group(1).strip().split('\n')
        taxa = []
        sequences = {}
        for line in lines:
            m = re.match(r'\[([-*]+)\]', line.strip())
            if m is not None:
                mask = m.group(1)
            else:
                stripped_line = re.sub(r'\[.+?\]', '', line).strip()
                if len(stripped_line) > 0:
                    parts = stripped_line.split()
                    assert len(parts) == 2, 'Found more than 2 parts to this line:\n%s' % line
                    taxon_name = re.sub(r'_', ' ', parts[0]).strip()
                    taxa.append(taxon_name)
                    sequences[taxon_name] = parts[1]

    return (ntax, nchar, mask, taxa, sequences)

import os,re

def writeNexusFile(fn, ntax, nchar, mask, taxa, sequences):
    if os.path.exists(fn):
        os.rename(fn, '%s.bak' % fn)
    longest = max([len(t) for t in taxa])
    taxonfmt = '  %%%ds' % longest
    f = open(fn, 'w')
    f.write('#nexus\n\n')
    f.write('begin data;\n')
    f.write('  dimensions ntax=%d nchar=%d;\n' % (ntax, nchar))
    f.write('  format datatype=dna gap=-;\n')
    f.write('  matrix\n')
    if mask is not None:
        f.write(taxonfmt % ' ')
        f.write('[%s]\n' % mask)
    for t in taxa:
        taxon_name = re.sub(r'\s+', '_', t)
        f.write(taxonfmt % taxon_name)
        f.write(' %s\n' % sequences[t])
    f.write('  ;\n')
    f.write('end;\n')
    f.close()

if __name__ == '__main__':
    random.seed(rnseed)
    prob_missing = missing_percent/100.0
    ntax, nchar, mask, taxa, sequences = readNexusFile(infilename)
    for t in taxa:
        newseq = ''
        for s in sequences[t]:
            u = random.uniform(0,1)
            if u < prob_missing:
                newseq += '?'
            else:
                newseq += s
        sequences[t] = newseq            
    writeNexusFile(outfilename, ntax, nchar, mask, taxa, sequences)
```
