## Supplementary material for "Estimating Bayesian Phylogenetic Information Content Using Geodesic Distances": make-nsites-rbop-slurm.py

```
import re, os, sys

stuff = open('slurm-nsites-rbop-template.sh', 'r').read()

# These are settings you can change
stuff,n = re.subn('__USEREMAIL__', '', stuff)
assert n == 1, 'Expecting 1 substitution for __USEREMAIL__, got %d' % n

stuff,n = re.subn('__NREPS__', '10', stuff)
assert n == 1, 'Expecting 1 substitution for __NREPS__, got %d' % n

stuff,n = re.subn('__MASTERSEED__', '12345', stuff)
assert n == 1, 'Expecting 1 substitution for __MASTERSEED__, got %d' % n

stuff,n = re.subn('__NARRAYTASKS__', '60', stuff)
assert n == 1, 'Expecting 1 substitution for __NARRAYTASKS__, got %d' % n

stuff,n = re.subn('__NVALUES__', '6', stuff)
assert n == 1, 'Expecting 1 substitution for __NVALUES__, got %d' % n

stuff,n = re.subn('__SGSEQLEN__', '-l$nsites', stuff)
assert n == 1, 'Expecting 1 substitution for __SGSEQLEN__, got %d' % n

outf = open('nsites-rbop-slurm.sh', 'w')
outf.write(stuff)
outf.close()
```
