## Supplementary material for "Estimating Bayesian Phylogenetic Information Content Using Geodesic Distances": slurm-rates-rbop-template.sh

```
#!/bin/bash
#SBATCH --job-name=rates             # job name
#SBATCH --array=1-__NARRAYTASKS__    # 1 to nelements*nreplicates as defined below
#SBATCH --partition=general          # Name of Partition
#SBATCH --ntasks=1                   # Maximum CPU cores for each job
#SBATCH --nodes=1                    # Ensure all cores are from the same node for each job
#SBATCH --cpus-per-task=1            # cpu-cores per task (>1 if multi-threaded tasks) for each job
#SBATCH --constraint='epyc128'       # Target the AMD Epyc 128-core node architecture
#SBATCH --mem=20G                    # Request 20GB of available RAM
#SBATCH --output=rates%a.out         # stdout: %a becomes the array id
#SBATCH --error=rates%a.err          # stderr: %a becomes the array id
#SBATCH --mail-type=END              # Event(s) that triggers email notification (BEGIN,END,FAIL,ALL)
#SBATCH --mail-user=__USEREMAIL__    # Destination email address

# Specify paths to programs used
basedir=`pwd`
RB=$HOME/bin/rb131
OP=$HOME/bin/op194
SG=$HOME/bin/seq-gen

# Set number of different values and number of replicates
nvalues=__NVALUES__
nreplicates=__NREPS__
ntotal=$(( $nvalues*$nreplicates ))

# Set master random number seed
RANDOM=__MASTERSEED__

# Load modules needed
module load python/3.12.5

# Generate seed and number of sites for each analysis
declare -a SEEDS
declare -a RATES
VALUES=(0.0001 0.001 0.01 0.1 1 10 100 1000)
for (( i=0; i<ntotal; i++ )); do
    SEEDS+=("$RANDOM")
    j=$(( $i % $nvalues ))
    RATES+=("${VALUES[$j]}")
done

idx=$(( $SLURM_ARRAY_TASK_ID - 1 ))
seed=${SEEDS[$idx]}
rate=${RATES[$idx]}
rep=$(( $idx / $nvalues ))

mkdir -p $basedir/rates
echo "***** $SLURM_ARRAY_TASK_ID: index = $idx, replicate = $rep, rate = $rate, seed = $seed *****" >> ./rates/repinfo.txt
echo "***** $SLURM_ARRAY_TASK_ID: index = $idx, replicate = $rep, rate = $rate, seed = $seed *****"

# Create the data directory
mkdir -p $basedir/rates/rep-$rep/rate-$rate/data

# Create the posterior directory
mkdir -p $basedir/rates/rep-$rep/rate-$rate/posterior
echo "rnseed <- $seed; run_under_prior <- FALSE" > rates/rep-$rep/rate-$rate/posterior/adhoc.Rev

# Create the prior directory
mkdir -p $basedir/rates/rep-$rep/rate-$rate/prior
echo "rnseed <- $seed; run_under_prior <- TRUE" > rates/rep-$rep/rate-$rate/prior/adhoc.Rev

# Simulate the data
echo
echo "***************************"
echo "***** Running seq-gen *****"
echo "***************************"
cd $basedir/rates/rep-$rep/rate-$rate/data
$SG -mGTR __SGSEQLEN__ -n1 -p1 -f 0.25 0.25 0.25 0.25 -r 1.0 4.0 1.0 1.0 4.0 1.0 -z$seed -on -s$rate < ../../../../modeltree.txt > sim.nex
cd ..

# Sample the posterior and compute the posterior mean tree and variance
echo
echo "****************************************"
echo "***** Running RevBayes (posterior) *****"
echo "****************************************"
cd $basedir/rates/rep-$rep/rate-$rate/posterior
$RB ../../../../fig-lcr.Rev

# Sample the prior and compute the prior mean tree
echo
echo "************************************"
echo "***** Running RevBayes (prior) *****"
echo "************************************"
cd $basedir/rates/rep-$rep/rate-$rate/prior
$RB ../../../../fig-lcr.Rev

echo
echo "*********************************"
echo "***** Calculate information *****"
echo "*********************************"
cd $basedir/rates/rep-$rep/rate-$rate
rm -rf results

python3 $basedir/../calcinfo.py --oppath $OP --hpdrad --radpct 95 --dirname results --prefix infohpd --skip 1 \
    --postfile posterior/output/simdata.trees --priorfile prior/output/simdata.trees 

python3 $basedir/../calcinfo.py --oppath $OP --radpct 95 --dirname results --prefix infostd --skip 1 \
    --postfile posterior/output/simdata.trees --priorfile prior/output/simdata.trees 
```
