## Supplementary material for "Estimating Bayesian Phylogenetic Information Content Using Geodesic Distances": summarize.py

```
from scipy.stats import describe
import argparse, os, sys, math, subprocess, glob, re

parser = argparse.ArgumentParser(
    formatter_class=argparse.RawDescriptionHelpFormatter,
    description='''Summarizes simulation results in the form of bar plots. LCR is computed using the 95% radius (RAD), 95% HPD radius (HPD), or the standard deviation (STD).''')
parser.add_argument('--dirname', required=True, help='directory containing asrv, missing, nsites, and rates directories')
parser.add_argument('--categname', required=True, choices=["asrv", "missing", "nsites", "rates"], help='specify which results to summarize')
parser.add_argument('--precision', type=int, default=5, help='number of digits precision')
settings = vars(parser.parse_args())

# Save settings into more convenient variables
dirname   = settings['dirname']
categname = settings['categname']
precision = settings['precision']

# Determine the current working directory
current_working_dir = os.getcwd()
print('current_working_dir = "%s"' % current_working_dir)

# Output function sends string argument to both screen and log file
outf = None
def output(s = None):
    global outf
    if s is None:
        print()
        outf.write('\n')
    else:
        print(s)
        outf.write('%s\n' % s)

# Dictionaries into which results are saved for plotting
valueSTDdict = {}
valueHPDdict = {}
valueRADdict = {}
    
# PlotJustOne function creates R script and produces a plot in the form of a PDF file  
# that has bars for just one of the three modes (RAD, HPD, or STD)
def PlotJustOne(fnprefix, values, means, stdevs):
    Rfn = '%s.R' % fnprefix
    PDFfn = '%s.pdf' % fnprefix
    output('  writing file "%s" and creating "%s"' % (Rfn, PDFfn))
    
    # Open the R script file for writing
    rfile = open(Rfn, 'w')
    
    # Import ggplot2
    rfile.write('library(ggplot2)\n')
    
    # Create a format string with the specified precision
    #fmtstr = '"%%.%df"' % precision
    
    # Set the current working directory
    rfile.write('cwd = system(\'cd "$( dirname "$0" )" && pwd\', intern = TRUE)\n')
    rfile.write('setwd(cwd)\n')
    
    # Open a PDF file for writing
    rfile.write('pdf("%s")\n' % PDFfn)

    # Let variable x hold the values (e.g. 0.0001, 0.001, 0.01, 0.1, 1, 10, 100, 1000)
    #xstr = [fmtstr % x for x in values]
    xstr = ['"%s"' % x for x in values]
    rfile.write('x <- c(%s)\n' % ','.join(xstr))

    # Let nvalues hold the number of values (e.g. 8)
    nvalues = len(values)

    # Let the variable ystd hold the mean info
    ystr = ['%g' % y for y in means]
    rfile.write('ystd <- c(%s)\n' % ','.join(ystr))

    # Let the variable zstd hold the standard deviation info
    sdstr = ['%g' % sd for sd in stdevs]
    rfile.write('zstd <- c(%s)\n' % ','.join(sdstr))
    
    # Construct the data frame, casting mode strings as factors
    rfile.write('df = data.frame(%s=x, mean=ystd, sd=zstd)\n' % categname)
    
    # Construct the minimum and maximum value of the error bars for each mean
    rfile.write('df$ymin <- df$mean - df$sd\n')
    rfile.write('df$ymax <- df$mean + df$sd\n')

	# Run ggplot, specifying values for x and means for y
    rfile.write('p <- ggplot(df, aes(x = %s, y = mean))\n' % categname)

	# Add the bars showing the means
	# stat="identity" plots pre-calculated summary statistics (could use geom_col which does this by default)
	# alpha = 0.75 adds transparency to the bars, which softens the colors
    rfile.write('p <- p + geom_col(fill = "#8eaff5", alpha=0.95)\n')

	# Add the error bars
	# width = 0.2 determines the width of the crossbars at the top and bottom of the error bars
    rfile.write('p <- p + geom_errorbar(aes(ymin = ymin, ymax = ymax), width = 0.2, position = position_dodge(width = 0.9))\n')

	# Add axis labels
    rfile.write('p <- p + labs(x = "", y = "Information Content")\n')
    
    # Specify the font size for the axis labels
    rfile.write('p <- p + theme(axis.text.x = element_text(size = 16))\n')
    rfile.write('p <- p + theme(axis.text.y = element_text(size = 16))\n')
    rfile.write('p <- p + theme(axis.title = element_text(size = 18))\n')

    # Print the plot
    rfile.write('print(p)\n')
    
    # Close the device
    rfile.write('dev.off()\n')
    
    # Close the R file
    rfile.close()
    
    # Use subprocess to run Rscript on the R file
    subprocess.Popen(['Rscript', Rfn]).communicate()
    
# PlotAllThree function creates R script and produces a plot in the form of a PDF file  
# that has bars for RAD, HPD, and STD modes  
def PlotAllThree(fnprefix, values, meansHPD, stdevsHPD, meansRAD, stdevsRAD, meansSTD, stdevsSTD):
    Rfn = '%s.R' % fnprefix
    PDFfn = '%s.pdf' % fnprefix
    output('  writing file "%s" and creating "%s"' % (Rfn, PDFfn))
    
    # Open the R script file for writing
    rfile = open(Rfn, 'w')
    
    # Import ggplot2
    rfile.write('library(ggplot2)\n')
    
    # Create a format string with the specified precision
    #fmtstr = '"%%.%df"' % precision
    
    # Set the current working directory
    rfile.write('cwd = system(\'cd "$( dirname "$0" )" && pwd\', intern = TRUE)\n')
    rfile.write('setwd(cwd)\n')
    
    # Open a PDF file for writing
    rfile.write('pdf("%s")\n' % PDFfn)

    # Let variable x hold the values (e.g. 0.0001, 0.001, 0.01, 0.1, 1, 10, 100, 1000)
    #xstr = [fmtstr % x for x in values]
    xstr = ['"%s"' % x for x in values]
    rfile.write('x <- c(%s)\n' % ','.join(xstr))

    # Let nvalues hold the number of values (e.g. 8)
    nvalues = len(values)

    # Let the variable yhpd hold the mean HPD info
    ystr = ['%g' % y for y in meansHPD]
    rfile.write('yhpd <- c(%s)\n' % ','.join(ystr))

    # Let the variable ystd hold the mean RAD info
    ystr = ['%g' % y for y in meansRAD]
    rfile.write('ystd <- c(%s)\n' % ','.join(ystr))

    # Let the variable yvar hold the mean STD info
    ystr = ['%g' % y for y in meansSTD]
    rfile.write('yvar <- c(%s)\n' % ','.join(ystr))

    # Let the variable zhpd hold the standard deviation for HPD info
    sdstr = ['%g' % sd for sd in stdevsHPD]
    rfile.write('zhpd <- c(%s)\n' % ','.join(sdstr))
    
    # Let the variable zstd hold the standard deviation for RAD info
    sdstr = ['%g' % sd for sd in stdevsRAD]
    rfile.write('zstd <- c(%s)\n' % ','.join(sdstr))
    
    # Let the variable zvar hold the standard deviation for STD info
    sdstr = ['%g' % sd for sd in stdevsSTD]
    rfile.write('zvar <- c(%s)\n' % ','.join(sdstr))
    
    # Column 1 of the data frame will hold nvalues of each mode
    rfile.write('col1 <- c(rep("HPD",%d), rep("RAD",%d), rep("STD",%d))\n' % (nvalues, nvalues, nvalues))

    # Column 2 of the data frame will hold a copy of the values for each mode
    rfile.write('col2 <- c(x, x, x)\n')

    # Column 3 of the data frame will hold means from each mode
    rfile.write('col3 <- c(yhpd, ystd, yvar)\n')

    # Column 4 of the data frame will hold standard deviations from each mode
    rfile.write('col4 <- c(zhpd, zstd, zvar)\n')
    
    # Construct the data frame, casting mode strings as factors
    rfile.write('df = data.frame(mode=factor(col1, levels=c("HPD", "RAD", "STD")), %s=col2, mean=col3, sd=col4)\n' % categname)
    
    # Construct the minimum and maximum value of the error bars for each mean
    rfile.write('df$ymin <- df$mean - df$sd\n')
    rfile.write('df$ymax <- df$mean + df$sd\n')

	# Run ggplot, specifying values for x, means for y, and color the bars by mode
    rfile.write('p <- ggplot(df, aes(x = %s, y = mean, fill = mode))\n' % categname)

	# Add the bars showing the means
	# stat="identity" plots pre-calculated summary statistics (could use geom_col which does this by default)
	# position = position_dodge means place bars site-by-side rather than stacked
	# alpha = 0.75 adds transparency to the bars, which softens the colors
    # rfile.write('p <- p + geom_bar(stat="identity", position=position_dodge(), alpha=0.75)\n')
    rfile.write('p <- p + geom_col(position=position_dodge(), alpha=0.75)\n')

	# Add the error bars
	# width = 0.2 determines the width of the crossbars at the top and bottom of the error bars
    rfile.write('p <- p + geom_errorbar(aes(ymin = ymin, ymax = ymax), width = 0.2, position = position_dodge(width = 0.9))\n')

	# Add axis labels
    rfile.write('p <- p + labs(x = "", y = "Information Content")\n')
    
    # Specify the font size for the axis labels
    rfile.write('p <- p + theme(axis.text.x = element_text(size = 16))\n')
    rfile.write('p <- p + theme(axis.text.y = element_text(size = 16))\n')
    rfile.write('p <- p + theme(axis.title = element_text(size = 18))\n')

    # Print the plot
    rfile.write('print(p)\n')
    
    # Close the device
    rfile.write('dev.off()\n')
    
    # Close the R file
    rfile.close()
    
    # Use subprocess to run Rscript on the R file
    subprocess.Popen(['Rscript', Rfn]).communicate()
    
def extractInfo(fn):
    stuff = open(fn, 'r').read()

    m = re.search(r'([0-9]+) = posterior sample size', stuff, re.M | re.S)
    assert m is not None, 'Could not extract posterior sample size from file "%s"' % fn
    post_sample_size = float(m.group(1))

    m = re.search(r'([0-9]+) = prior sample size', stuff, re.M | re.S)
    assert m is not None, 'Could not extract prior sample size from file "%s"' % fn
    prior_sample_size = float(m.group(1))

    m = re.search(r'([.0-9]+) = posterior mean tree length', stuff, re.M | re.S)
    assert m is not None, 'Could not extract posterior mean tree length from file "%s"' % fn
    post_mean_TL = float(m.group(1))

    m = re.search(r'([.0-9]+) = prior mean tree length', stuff, re.M | re.S)
    assert m is not None, 'Could not extract prior mean tree length from file "%s"' % fn
    prior_mean_TL = float(m.group(1))

    m = re.search(r'([.0-9]+) = posterior variance', stuff, re.M | re.S)
    assert m is not None, 'Could not extract posterior variance from file "%s"' % fn
    post_var = float(m.group(1))

    m = re.search(r'([.0-9]+) = prior variance', stuff, re.M | re.S)
    assert m is not None, 'Could not extract prior variance from file "%s"' % fn
    prior_var = float(m.group(1))

    m = re.search(r'([.0-9]+) = posterior scaler', stuff, re.M | re.S)
    assert m is not None, 'Could not extract posterior scaler from file "%s"' % fn
    post_scaler = float(m.group(1))

    m = re.search(r'([.0-9]+) = prior scaler', stuff, re.M | re.S)
    assert m is not None, 'Could not extract prior scaler from file "%s"' % fn
    prior_scaler = float(m.group(1))

    m = re.search(r'([.0-9]+) = posterior 95% radius', stuff, re.M | re.S)
    assert m is not None, 'Could not extract posterior 95% radius from file "%s"' % fn
    post_radius = float(m.group(1))

    m = re.search(r'([.0-9]+) = prior 95% radius', stuff, re.M | re.S)
    assert m is not None, 'Could not extract prior 95% radius from file "%s"' % fn
    prior_radius = float(m.group(1))

    m = re.search(r'([-.0-9]+) = LCRrad', stuff, re.M | re.S)
    assert m is not None, 'Could not extract LCRrad from file "%s"' % fn
    LCRrad = float(m.group(1))

    m = re.search(r'([-.0-9]+) = information as percentage', stuff, re.M | re.S)
    assert m is not None, 'Could not extract information as percentage from file "%s"' % fn
    Irad = float(m.group(1))

    return {
        'post_sample_size': post_sample_size,
        'prior_sample_size': prior_sample_size,
        'post_mean_TL': post_mean_TL,
        'prior_mean_TL': prior_mean_TL,
        'post_var': post_var,
        'prior_var': prior_var,
        'post_scaler': post_scaler,
        'prior_scaler': prior_scaler,
        'post_radius': post_radius,
        'prior_radius': prior_radius,
        'LCRrad': LCRrad,
        'Irad': Irad
    }
        
def summaryTable(what):
    output('\nSummary of %s for %s:' % (what,categname))
    
    # Get row headers and sort
    # Arbitrarily using valueHPDdict here, valueRADdict and valueSTDdict 
    # would also work because each have the same keys
    value_strings_in_order = []
    for value in valueHPDdict.keys():
        value_strings_in_order.append(value)
    value_strings_in_order.sort()
    
    # Convert row headers to floats
    values_in_order = []
    for valuestr in value_strings_in_order:
        values_in_order.append(float(valuestr))
    
    assert len(value_strings_in_order) > 0
    assert len(values_in_order) > 0
    
    # Column headers
    output('%12s %12s %12s %12s %12s %12s %12s' % ('mode', 'values','mean','sd','nobs','min','max'))
    
    # Create table
    meansHPD = []
    stdevsHPD = []
    meansRAD = []
    stdevsRAD = []
    meansSTD = []
    stdevsSTD = []
    for k in value_strings_in_order:
        dHPD = describe(valueHPDdict[k][what], 0, 1)  # 2nd arg 0 is axis, 3rd arg 1 is df
        meansHPD.append(dHPD.mean)
        stdevsHPD.append(math.sqrt(dHPD.variance))

        dRAD = describe(valueRADdict[k][what], 0, 1)  # 2nd arg 0 is axis, 3rd arg 1 is df
        meansRAD.append(dRAD.mean)
        stdevsRAD.append(math.sqrt(dRAD.variance))

        dSTD = describe(valueSTDdict[k][what], 0, 1)  # 2nd arg 0 is axis, 3rd arg 1 is df
        meansSTD.append(dSTD.mean)
        stdevsSTD.append(math.sqrt(dSTD.variance))

        fmtstr = '%%12s %%12.%df %%12.%df %%12.%df %%12d %%12.%df %%12.%df' % (precision,precision,precision,precision,precision)
        output(fmtstr % ('HPD', float(k), dHPD.mean, math.sqrt(dHPD.variance), dHPD.nobs, dHPD.minmax[0], dHPD.minmax[1]))
        output(fmtstr % ('RAD', float(k), dRAD.mean, math.sqrt(dRAD.variance), dRAD.nobs, dRAD.minmax[0], dRAD.minmax[1]))
        output(fmtstr % ('STD', float(k), dSTD.mean, math.sqrt(dSTD.variance), dSTD.nobs, dSTD.minmax[0], dSTD.minmax[1]))
        
    PlotJustOne(os.path.join('plots','%s-%s-STD-info' % (categname, what)), value_strings_in_order, meansSTD, stdevsSTD)
    PlotJustOne(os.path.join('plots','%s-%s-RAD-info' % (categname, what)), value_strings_in_order, meansRAD, stdevsRAD)
    PlotJustOne(os.path.join('plots','%s-%s-HPD-info' % (categname, what)), value_strings_in_order, meansHPD, stdevsHPD)
    PlotAllThree(os.path.join('plots','%s-%s-STD-RAD-HPD-info' % (categname, what)), value_strings_in_order, meansHPD, stdevsHPD, meansRAD, stdevsRAD, meansSTD, stdevsSTD)

if __name__ == '__main__':
    # Create plots directory if it does not already exist
    if not os.path.exists('plots'):
        os.mkdir('plots')

    # Open an output file in the plots directory
    outf = open(os.path.join('plots', '%s-output.txt' % categname), 'w')

    # Show the specified settings
    output('Settings:')
    output('  dir       = "%s"' % dirname)
    output('  categ     = %s' % categname)
    output('  precision = %d' % precision)
    
    # Compile a list (repdirs) of all replicate directories
    # Using rates as an example:
    # rates
    #     rep-0
    #         rate-0.1
    #         rate-0.01
    #         rate-0.001
    #         rate-0.0001
    #         rate-1
    #         rate-10
    #         rate-100
    #         rate-1000
    #     rep-1
    #         rate-0.1
    #         rate-0.01
    #         rate-0.001
    #         rate-0.0001
    #         rate-1
    #         rate-10
    #         rate-100
    #         rate-1000
    #      .
    #      .
    #      .
    #     rep-9
    #         rate-0.1
    #         rate-0.01
    #         rate-0.001
    #         rate-0.0001
    #         rate-1
    #         rate-10
    #         rate-100
    #         rate-1000
    #
    # repdirs = ['results/rates/rep-0', 'results/rates/rep-1', ..., 'results/rates/rep-9']        
    repdirs = glob.glob(os.path.join(dirname, categname,'rep-*'))
    for repdir in repdirs:
        # Compile a list of all category-specific directories inside repdir
        categdirglob = ''
        if categname == 'asrv':
            categdirglob = 'ratevar-*'
        elif categname == 'missing':
            categdirglob = 'missing-*'
        elif categname == 'rates':
            categdirglob = 'rate-*'
        elif categname == 'nsites':
            categdirglob = 'nsites-*'
        else:
            assert False, 'categname not valid'
            
        # Get list of directories matching specified category
        globname = os.path.join(repdir, categdirglob)
        categdirs = glob.glob(globname)
        
        # categdirs = ['results/rates/rep-0/rate-0.1', 'results/rates/rep-0/rate-0.01', ..., 'results/rates/rep-9/rate-1000']
        for catdir in categdirs:
            # Let valuestr equal the value of the category embedded in the directory name
            # For example: valuestr="0.001" for catdir='results/rates/rep-0/rate-0.001'
            # In this case, categdirglob = 'rate-*' so categdirglob[:-1] = 'rate-'
            regex = '.+' + categdirglob[:-1] + '([.0-9]+)$'
            m = re.match(regex, catdir)
            assert m is not None, 'regex "%s" did not match anything in "%s"' % (regex,catdir)
            valuestr = m.group(1)
            
            # Gather HPD info
            fn = os.path.join(catdir, 'results', 'infohpd-output.txt')
            infoHPDdict = extractInfo(fn)
            if valuestr in valueHPDdict.keys():
                v = valueHPDdict[valuestr]
                for k in infoHPDdict.keys():
                    v[k].append(infoHPDdict[k])
            else:
                valueHPDdict[valuestr] = {
                    'post_sample_size':[infoHPDdict['post_sample_size']],
                    'prior_sample_size':[infoHPDdict['prior_sample_size']],
                    'post_mean_TL':[infoHPDdict['post_mean_TL']],
                    'prior_mean_TL':[infoHPDdict['prior_mean_TL']],
                    'post_var':[infoHPDdict['post_var']],
                    'prior_var':[infoHPDdict['prior_var']],
                    'post_scaler':[infoHPDdict['post_scaler']],
                    'prior_scaler':[infoHPDdict['prior_scaler']],
                    'post_radius':[infoHPDdict['post_radius']],
                    'prior_radius':[infoHPDdict['prior_radius']],
                    'LCRrad':[infoHPDdict['LCRrad']],
                    'Irad':[infoHPDdict['Irad']]
                }

            # Gather RAD info
            fn = os.path.join(catdir, 'results', 'infostd-output.txt')
            infoRADdict = extractInfo(fn)
            if valuestr in valueRADdict.keys():
                v = valueRADdict[valuestr]
                for k in infoRADdict.keys():
                    v[k].append(infoRADdict[k])
            else:
                valueRADdict[valuestr] = {
                    'post_sample_size':[infoRADdict['post_sample_size']],
                    'prior_sample_size':[infoRADdict['prior_sample_size']],
                    'post_mean_TL':[infoRADdict['post_mean_TL']],
                    'prior_mean_TL':[infoRADdict['prior_mean_TL']],
                    'post_var':[infoRADdict['post_var']],
                    'prior_var':[infoRADdict['prior_var']],
                    'post_scaler':[infoRADdict['post_scaler']],
                    'prior_scaler':[infoRADdict['prior_scaler']],
                    'post_radius':[infoRADdict['post_radius']],
                    'prior_radius':[infoRADdict['prior_radius']],
                    'LCRrad':[infoRADdict['LCRrad']],
                    'Irad':[infoRADdict['Irad']]
                }

            # Gather STD info
            # Yes, a little redundant, but safer to just call extractInfo again 
            # rather than to write confusing conditional code
            fn = os.path.join(catdir, 'results', 'infostd-output.txt')
            infoSTDdict = extractInfo(fn)

            # Recalculate LCRrad and Irad based on standard deviation rather than radius
            newLCRrad  = 0.5*math.log(infoSTDdict['prior_var']) + math.log(infoSTDdict['prior_scaler'])
            newLCRrad -= 0.5*math.log(infoSTDdict['post_var']) + math.log(infoSTDdict['post_scaler'])
            newIrad = 100.0*(1.0 - math.exp(-newLCRrad))

            # # Recalculate LCRrad and Irad based on variance rather than radius
            # newLCRrad  = math.log(infoSTDdict['prior_var']) + 2.0*math.log(infoSTDdict['prior_scaler'])
            # newLCRrad -= math.log(infoSTDdict['post_var']) + 2.0*math.log(infoSTDdict['post_scaler'])
            # newIrad = 100.0*(1.0 - math.exp(-newLCRrad))
            
            # Replace values
            infoSTDdict['LCRrad'] = newLCRrad
            infoSTDdict['Irad'] = newIrad
            
            if valuestr in valueSTDdict.keys():
                v = valueSTDdict[valuestr]
                for k in infoSTDdict.keys():
                    v[k].append(infoSTDdict[k])
            else:
                valueSTDdict[valuestr] = {
                    'post_sample_size':[infoSTDdict['post_sample_size']],
                    'prior_sample_size':[infoSTDdict['prior_sample_size']],
                    'post_mean_TL':[infoSTDdict['post_mean_TL']],
                    'prior_mean_TL':[infoSTDdict['prior_mean_TL']],
                    'post_var':[infoSTDdict['post_var']],
                    'prior_var':[infoSTDdict['prior_var']],
                    'post_scaler':[infoSTDdict['post_scaler']],
                    'prior_scaler':[infoSTDdict['prior_scaler']],
                    'post_radius':[infoSTDdict['post_radius']],
                    'prior_radius':[infoSTDdict['prior_radius']],
                    'LCRrad':[infoSTDdict['LCRrad']],
                    'Irad':[infoSTDdict['Irad']]
                }

    summaryTable('LCRrad')
    outf.close()
```
