## Supplementary material for "Estimating Bayesian Phylogenetic Information Content Using Geodesic Distances": _readme.txt

To create fig-dissonance:

Directory layout showing only directories and files mentioned in the instructions below:

  supplementary-materials
      calcinfo.py
      3-fig-dissonance
*         diss
*           bhvdist
*           dissonancde
*           rep1
*             data
*             output
*             sim1
*             sim2
*             .
*             .
*             .
*             sim6
*             startingTree.Rev
*           rep2
*             data
*             output
*             sim1
*             sim2
*             .
*             .
*             .
*             sim6
*             startingTree.Rev
*           .
*           .
*           .
*           rep20
*             data
*             output
*             sim1
*             sim2
*             .
*             .
*             .
*             sim6
*             startingTree.Rev
*           bhv-vs-step.pdf
*           bhv-vs-step.R
*           diss-vs-bhv.pdf
*           diss-vs-bhv.R
          _readme.txt
          deploy.py
*         deploysettings.py
          fig-dissonance.Rev
          slurm-template.sh
          summarize.py

* file/directory created as a result of these instructions

1. Edit slurm-template.sh and adapt the SBATCH commands at the top 
    to your cluster. Be sure the paths to the Seq-Gen, RevBayes, 
    and OP executables are correct.

    If you do not have these programs installed, here is where you can get them:
        Seq-Gen:    https://github.com/rambaut/Seq-Gen/releases/tag/1.3.4
        OP:         https://github.com/plewis/OP
        RevBayes:   https://revbayes.github.io/download

2. Run deploy.py:

    python3 deploy.py --dirname diss
    
    The deploy.py script creates the directory structure (sim directories within each
    rep directory). It calls RevBayes to draw a starting tree from the prior and then
    OP to follow a random path starting with that tree. It then calls Seq-Gen to 
    simulate data for the starting and ending trees. 

3. Specify "hpc" as an alias to your remote cluster in  ~/.ssh/config and
   create a "work/diss" directory inside your home directory on your cluster
   
   ssh hpc
   mkdir -p work
    
4. Upload files to the "work/diss" directories on your cluster:

    rsync -azPv diss hpc:work
        
9. Run the analyses on the cluster:

    ssh hpc
    cd work/diss
    sbatch slurm.sh
    
    The slurm script calls RevBayes to obtain a sample from both prior and posterior
    of both start and end data sets for each sim within each rep. The calcinfo.py script
    is called to calculate information content, and this script in turn calls OP to do
    estimate the mean tree and compute variances and radii. The information is stored 
    in the results directory within each sim directory.
    
10. Bring results back to your local computer:

    rsync -azPv hpc:work/diss/ diss/

11. Summarize results:

    python3 summarize.py --mode hpd
    python3 summarize.py --mode rad
    
    The summarize.py script visits each sim directory within each replicated directory,
    computing the dissonance between the start and end datasets. OP is called to 
    calculate the distance between mean trees, but the remainder of the dissonance
    calculation is done in python.
    
    If mode is "hpd" then the dissonance is based on the 95% credible set of trees
    from each posterior. If mode is "rad", the dissonance is based on the 95% of trees
    closest to the mean tree.
    
    Two plots are created during each invocation of summarize.py. One plot shows 
    dissonance in units of pooled standard deviation (diss-vs-bhv-hpd-stddev.pdf and
    diss-vs-bhv-rad-stddev.pdf). The other plot shows dissonance in units of pooled 
    radius (diss-vs-bhv-hpd-radius.pdf and diss-vs-bhv-rad-radius.pdf).
    
    The paper reports the mean geodesic distance and std. dev. between starting and 
    ending trees (mean = 0.606208850, s.d. = 0.066439036, n = 20), the dissonance when 
    both data sets are simulated from the same starting tree (mean = 0.124231645, 
    s.d. = 0.011204318, n = 20), the total dissonance between starting and ending 
    data sets (mean = 0.734085101, s.d. = 0.083814257, n = 10), the number of segments 
    that failed to show an increase (4), and the correlation between geodesic distance 
    and dissonance (correlation = 0.961075611, pvalue = 0.000000000, n = 120). These
    numbers are all reported in the file 3-fig-dissonance/diss/diss-vs-bhv-rad-radius.R.
    
