## Supplementary material for "Estimating Bayesian Phylogenetic Information Content Using Geodesic Distances": deploy.py

```
import argparse, sys, os, random, subprocess, re, math

# Uses RevBayes to choose random starting tree from prior.
# NNI moves in RevBayes are used to move some distance (determined
# by niters) away from starting tree.
# This path from starting tree to ending tree is divided into nstations=5 chunks
# Data are simulated from each station defined by the vector stations
# Dissonance is calculated between starting data set and the data set 
# generated for each station. Note that the station at 0.0 is included so
# that dissonance is determined for two independent data sets simulated
# from the same starting model tree. All this is repeated nreps times.
# Plot shows true distance between model trees on x-axis and
# dissonance on y-axis. Expectation is that dissonance will rise
# monotonically with increasing geodesic distance between model trees.

use_ggplot = True

parser = argparse.ArgumentParser(
    formatter_class=argparse.RawDescriptionHelpFormatter,
    description='''Calculates dissonance for a series of increasingly distant model trees.''',
    epilog='''Be sure OPpath is the correct path to the OP executable. OP may be obtained from https://github.com/plewis/OP.''')

parser.add_argument('--dirname',   required=True,             help='directory to be created for storing results')
parser.add_argument('--oppath',    default='op',              help='path to OP executable')
parser.add_argument('--nreps',     default=20,    type=int,   help='number of independent replicate simulations')
parser.add_argument('--nsteps',    default=500,   type=int,   help='number of steps to take in random walk for each replicate')
parser.add_argument('--niters',    default=10000, type=int,   help='number of steps to run posterior MCMC')
parser.add_argument('--rnseed',    default=12345, type=int,   help='pseudorandom number seed')
parser.add_argument('--tlshape',   default=5.0,   type=float, help='edge length proportion prior dirichlet parameter')
parser.add_argument('--dirparam',  default=5.0,   type=float, help='edge length proportion prior dirichlet parameter')
parser.add_argument('--stepmu',    default=0.0,   type=float, help='mean of normal variate used to modify edge length in random walk')
parser.add_argument('--stepsigma', default=0.005, type=float, help='standard deviation of normal variate used to modify edge length in random walk')

# Example of a setting that has a default of None
parser.add_argument('--postfile', default=None, help='file name of posterior tree file')

settings = vars(parser.parse_args())

masterdir = settings['dirname']
OPpath    = settings['oppath']
nreps     = settings['nreps']
nsteps    = settings['nsteps']
niters    = settings['niters']
rnseed    = settings['rnseed']
tlshape   = settings['tlshape']
dirparam  = settings['dirparam']
stepmu    = settings['stepmu']
stepsigma = settings['stepsigma']

# These settings fixed an not modifiable from the command line
stations  = [0.0, 0.2, 0.4, 0.6, 0.8, 1.0] # stops along the path from starting to ending tree
nstations = len(stations) # number of stops along the path from starting to ending tree

# Save settings to a file to be imported by harvest.py
settingsf = open('deploysettings.py', 'w')
settingsf.write("masterdir = '%s'\n" % masterdir)
settingsf.write("OPpath    = '%s'\n" % OPpath   )
settingsf.write('nreps     = %d\n' % nreps    )
settingsf.write('nsteps    = %d\n' % nsteps   )
settingsf.write('niters    = %d\n' % niters   )
settingsf.write('rnseed    = %d\n' % rnseed   )
settingsf.write('tlshape   = %g\n' % tlshape  )
settingsf.write('dirparam  = %g\n' % dirparam )
settingsf.write('stepmu    = %g\n' % stepmu   )
settingsf.write('stepsigma = %g\n' % stepsigma)
settingsf.write('nstations = %d\n' % nstations   )
settingsf.write('stations  = [0.0, 0.2, 0.4, 0.6, 0.8, 1.0]\n')
settingsf.close()

random.seed(rnseed)
dissdir = os.path.join(masterdir, 'dissonance')
bhvdir  = os.path.join(masterdir, 'bhvdist')

if __name__ == '__main__':
    # Program paths
    current_working_dir = os.getcwd()
    #scriptdir = os.path.dirname(os.path.realpath(sys.argv[0]))
    RB = 'rb132'     # Used only to sample model trees from the prior, not to sample posteriors
    OP = 'op'        # Used to determine geodesic distance between trees
    SG = 'seq-gen'   # Used to simulate data from a specified model tree
    
    # Sanity check to make sure current working directory is what we thought
    if not os.path.exists(os.path.join(current_working_dir, '..', 'calcinfo.py')):
        sys.exit('I was assuming you would call %(prog)s from a directory one level above calcinfo.py; current directory is "%s"' % current_working_dir)
    
    # Determine number of cores needed for slurm array job
    array_size = nreps*nstations

    # Create master directory (refuse to continue if it already exists)
    if os.path.exists(masterdir):
        sys.exit('Please remove/rename directory "%s" and run again' % masterdir)
    os.mkdir(masterdir)
    
    # Create <masterdir>/dissonance subdirectory
    os.mkdir(dissdir)
    
    # Create <masterdir>/bhvdist subdirectory
    os.mkdir(bhvdir)

    # Generate model trees and calculate BHV distances
    for rep in range(nreps):
        rep_plus_one = rep + 1
        
        # Create rep directory
        repdir = os.path.join(masterdir, 'rep%d' % rep_plus_one)
        os.mkdir(repdir)
        
        # Create output directory
        outputdir = os.path.join(repdir, 'output')
        os.mkdir(outputdir)
                
        # Use RevBayes to draw a random starting tree from the prior
        startf = open(os.path.join(repdir, 'startingTree.Rev'), 'w')
        startf.write('seed(330607)\n')
        startf.write('num_taxa <- 26\n')
        startf.write('num_branches <- 2 * num_taxa - 3\n')
        startf.write('taxa <- [taxon("A"),taxon("B"),taxon("C"),taxon("D"),taxon("E"),taxon("F"),taxon("G"),taxon("H"),taxon("I"),taxon("J"),taxon("K"),taxon("L"),taxon("M"),taxon("N"),taxon("O"),taxon("P"),taxon("Q"),taxon("R"),taxon("S"),taxon("T"),taxon("U"),taxon("V"),taxon("W"),taxon("X"),taxon("Y"),taxon("Z")]\n')
        startf.write('outgroup <- clade("A")\n')
        startf.write('print("#nexus")\n')
        startf.write('print("begin trees;")\n')
        startf.write('starting_topology ~  dnUniformTopology(taxa, outgroup=outgroup)\n')
        startf.write('starting_TL       ~  dnGamma(shape=5.0, rate=5.0/2.66809)  # mean TL is 2.66809\n')
        startf.write('starting_edgeprop ~  dnDirichlet( rep(5.0, num_branches) )\n')
        startf.write('starting_edgelens := starting_edgeprop * starting_TL\n')
        startf.write('starting_psi      := treeAssembly(starting_topology, starting_edgelens)\n')
        startf.write('print("  tree starttree = ", starting_psi)\n')
        startf.write('print("end;")\n')
        startf.close()
    
        # Run RevBayes to generate random prior path through treespace
        outfile = open(os.path.join(outputdir, 'starting.tre'), 'w')
        ok = True
        try:
            subprocess.run([RB, 'startingTree.Rev'], stdout=outfile, cwd=repdir, check=True)
        except subprocess.CalledProcessError as e:
            print('RevBayes failed to create a starting tree for rep %d' % rep_plus_one)
            ok = False
        except FileNotFoundError:
            print('Error: Executable or file not found.')
            ok = False
        outfile.close()
        if not ok:
            sys.exit('\nAborted.')

        # Run OP to perform a random walk starting with the tree generated by RevBayes
        opf = open(os.path.join(outputdir, 'op.conf'),'w')
        opf.write('prefix = randompath\n')
        opf.write('seed = %d\n' % random.randint(1, 99999))
        opf.write('reftree = starting.tre\n')
        opf.write('refskip = 0\n')
        #opf.write('refrooted = yes\n')
        opf.write('refscale = 1.0\n')
        opf.write('nsteps = %d\n' % nsteps)
        opf.write('stepmu = %g\n' % stepmu)
        opf.write('stepsigma = %g\n' % stepsigma)
        opf.close()
        subprocess.run([OP], cwd=outputdir)
                
    # Harvest BHV distances and tree descriptions 
    # Sort both by MCMC iteration   
    bhvrep = []     # contains nreps lists, each containing nstations (tree index, bhv) tuples
    newicksrep = [] # contains nreps lists, each containing nstations (tree index, newick) tuples
    xmax = None
    ymax = None
    for rep in range(nreps):
        rep_plus_one = rep + 1
        repdir = os.path.join(masterdir, 'rep%d' % rep_plus_one)
    
        # # Read in nsteps + 1 tree distances from randompath.R
        # bhvdists = [] # list of (tree index, bhv distance) tuples
        # stuff = open(os.path.join(repdir, 'output', 'randompath.R'),'r').read()
        # m = re.search(r'bhvdist <- c\(.+?\)', stuff, re.M | re.S)
        # assert m is not None
        # tmp = m.group(1))
        # parts = tmp.split(',')
        # for p in parts:
        #     bhvdists.append(float(p.strip()))
        # assert len(bhvdists) == nsteps + 1

        # Read in nsteps + 1 tree distances and tree descriptions from randompath.tre
        # t0 is the starting tree
        bhvdists = [] # list of (tree index, bhv distance) tuples
        newicks = []  # list of (tree index, newick) tuples
        lines = open(os.path.join(repdir, 'output', 'randompath.tre'),'r').readlines()
        for line in lines:
            m = re.match(r'\s*tree t(\d+) \[bhvdist = ([.0-9]+), nNNIs = ([0-9]+)\]\s*=\s*\[&\S\]\s*(.+)', line)
            if m is not None:
                t = int(m.group(1))
                d = float(m.group(2))
                nni = float(m.group(3))
                newick = m.group(4)
                if xmax is None or t > xmax:
                    xmax = t
                if ymax is None or d > ymax:
                    ymax = d
                bhvdists.append((t,d))
                newicks.append((t,newick))
        nbhv = len(bhvdists)
        print('Found %d BHV distances for rep %d' % (nbhv, rep_plus_one))  
                
        # Sort to ensure both vectors are sorted by tree index (should be true anyway)
        bhvrep.append(sorted(bhvdists))
        newicksrep.append(sorted(newicks))
                    
    # Create <masterdir>/slurm.sh from slurm-template.sh
    slurm_contents = open(os.path.join('slurm-template.sh'), 'r').read()
    slurmf = open(os.path.join(masterdir, 'slurm.sh'), 'w')
    slurm_contents, nsub = re.subn('__ARRAY_SIZE__', '%d' % array_size, slurm_contents, re.M | re.S)
    assert nsub == 2, 'Expecting 2 subs, found $d' % nsub
    slurm_contents, nsub = re.subn('__NSIMS__', '%d' % nstations, slurm_contents, re.M | re.S)
    assert nsub == 2, 'Expecting 2 subs, found %d' % nsub
    slurmf.write(slurm_contents)
    slurmf.close()
    
    # Create <masterdir>/fig-dissonance.Rev from fig-dissonance.Rev
    rev_contents = open(os.path.join('fig-dissonance.Rev'), 'r').read()
    revf = open(os.path.join(masterdir, 'fig-dissonance.Rev'), 'w')
    revf.write(rev_contents)
    revf.close()
    
    # Create <masterdir>/calcinfo.py from scripts/calcinfo.py
    calcinfo_contents = open(os.path.join(current_working_dir, '..', 'calcinfo.py'), 'r').read()
    calcinfof = open(os.path.join(masterdir, 'calcinfo.py'), 'w')
    calcinfof.write(calcinfo_contents)
    calcinfof.close()
    
    # Create <masterdir>/treedist.R
    rff = open(os.path.join(masterdir, 'treedist.R'), 'w')
    rff.write('library("TreeDist")\n')
    rff.write('rep <- 1:%d\n' % nreps)
    rff.write('f <- function(rep) {\n')
    rff.write('    fn <- paste("rep", rep[1], "/", "output", "/", "priorpath-lambda.tre", sep="")\n')
    rff.write('    cat(sprintf("  fn = %s\n", fn))\n')
    rff.write('    trees <- ape::read.nexus(fn)\n')
    rff.write('    TreeDistance(trees, trees)\n')
    rff.write('}\n')
    rff.write('lapply(X=rep, FUN=f)\n')
    rff.close()
    
    # Simulate datasets for nstations of the reps
    for rep in range(nreps):
        rep_plus_one = rep + 1
        repdir = os.path.join(masterdir, 'rep%d' % rep_plus_one)
        
        # Create a data directory for this rep
        datadir = os.path.join(repdir, 'data') 
        if not os.path.exists(datadir):
            os.mkdir(datadir)
            
        # Save the reference newick in the file tree-ref.txt
        assert newicksrep[rep][0][0] == 0, 'Expecting first tree index to be 0 in newicksrep[%d][0] list' % rep
        newick0 = newicksrep[rep][0][1] # newick string from 0th tuple in prior path
        reftreefile = 'tree-ref.txt'
        treef = open(os.path.join(datadir, reftreefile), 'w')
        treef.write('%s\n' % newick0)
        treef.close()
        
        for j in range(nstations):
            sim = j
            sim_plus_one = j + 1
            station = int(stations[j]*nsteps)
            assert station <= len(newicksrep[rep]), 'station (%d) should be less than or equal to length of newicksrep (%d)' % (station, len(newicksrep[rep]))
            assert len(newicksrep[rep]) == len(bhvrep[rep]), 'Length of newicksrep (%d) should equal the length of bhvrep (%d)' % (len(newicksrep[rep]), len(bhvrep[rep]))
            
            # Retrieve stationth newick from ith replicate
            try:
                newick = newicksrep[rep][station][1]
            except IndexError:
                print('IndexError: station = %d' % station)
                print('  newicksrep has length %d' % (len(newicksrep),))
                print('  nstations is %d' % nstations)
                sys.exit('Aborted.')
    
            # Save true BHV distance between reference tree and end tree
            bhvf = open(os.path.join(bhvdir, 'bhv-rep%d-sim%d.txt' % (rep_plus_one, sim_plus_one)), 'w')
            bhvf.write('rep\tsim\taddress\tbhv\n')
            bhvf.write('%d\t%d\t%d\t%g\n' % (rep_plus_one, sim_plus_one, station, bhvrep[rep][station][1]))
            bhvf.close()
            
            # Save tree in file
            treefile = 'tree-%d.txt' % sim_plus_one
            treef = open(os.path.join(datadir, treefile), 'w')
            treef.write('%s\n' % newick)
            treef.close()
            
            # Run Seq-gen to create data using the reference tree
            seqgen_command = [SG, 
                '-mGTR', 
                '-l1000', 
                '-n1', 
                '-p1', 
                '-on', 
                '-f', '0.25', '0.25', '0.25', '0.25', 
                '-r', '1.0', '1.0', '1.0', '1.0', '1.0', '1.0', 
                '-z%d' % random.randint(1,99999)]
            refseqfile = 'data0-%d.nex' % sim_plus_one
            
            infile  = open(os.path.join(datadir, reftreefile), 'r')
            outfile = open(os.path.join(datadir, refseqfile), 'w')
                    
            try:
                # Run the command, redirecting stdin from infile and stdout to outfile
                subprocess.run(seqgen_command, stdin=infile, stdout=outfile, check=True)
            except subprocess.CalledProcessError as e:
                print('Seq-gen failed to simulate reference data for iter %d, rep %d {e.returncode}' % (j, rep_plus_one))
                sys.exit('\nAborted.')
            except FileNotFoundError:
                print('Error: Executable or file not found.')
                sys.exit('\nAborted.')
                
            infile.close()
            outfile.close()
                        
            # Run Seq-gen to create data using the tree from iteration j
            seqgen_command = [SG, 
                '-mGTR', 
                '-l1000', 
                '-n1', 
                '-p1', 
                '-on', 
                '-f', '0.25', '0.25', '0.25', '0.25', 
                '-r', '1.0', '1.0', '1.0', '1.0', '1.0', '1.0', 
                '-z%d' % random.randint(1,99999)]
            seqfile = 'data-%d.nex' % sim_plus_one
            
            infile  = open(os.path.join(datadir, treefile), 'r')
            outfile = open(os.path.join(datadir, seqfile), 'w')
                    
            try:
                # Run the command, redirecting stdin from infile and stdout to outfile
                subprocess.run(seqgen_command, stdin=infile, stdout=outfile, check=True)
            except subprocess.CalledProcessError as e:
                print('Seq-gen failed to simulate data for iter %d, rep %d {e.returncode}' % (j, rep_plus_one))
                sys.exit('\nAborted.')
            except FileNotFoundError:
                print('Error: Executable or file not found.')
                sys.exit('\nAborted.')
                        
            infile.close()
            outfile.close()
            
            # Create directory for this segment of the path
            simdir = 'sim%d' % sim_plus_one
            os.mkdir(os.path.join(repdir, simdir))
            
            # Create start directory
            repsimstartdir = os.path.join(repdir, simdir, 'start')
            os.mkdir(repsimstartdir)

            # Create post directory inside start directory
            repsimstartpostdir = os.path.join(repsimstartdir, 'post')
            os.mkdir(repsimstartpostdir)
            adhocf = open(os.path.join(repsimstartpostdir, 'adhoc.Rev'), 'w')
            adhocf.write('rnseed       <- %d\n' % random.randint(1,99999))
            adhocf.write('niter        <- %d\n' % niters)
            adhocf.write('tlshape      <- %d\n' % tlshape)
            adhocf.write('dirparam     <- %d\n' % dirparam)
            adhocf.write('samplefreq   <- 1\n')
            adhocf.write('printfreq    <- 100\n')
            adhocf.write('datafile     <- "../../../data/data0-%d.nex"\n' % sim_plus_one)
            adhocf.write('treefile     <- "post.trees"\n')
            adhocf.write('logfile      <- "post.log"\n')
            adhocf.write('prior_path   <- FALSE\n')
            adhocf.write('sample_prior <- FALSE\n')
            adhocf.close()

            # Create prior directory inside start directory
            repsimstartpriordir = os.path.join(repsimstartdir, 'prior')
            os.mkdir(repsimstartpriordir)
            adhocf = open(os.path.join(repsimstartpriordir, 'adhoc.Rev'), 'w')
            adhocf.write('rnseed       <- %d\n' % random.randint(1,99999))
            adhocf.write('niter        <- %d\n' % niters)
            adhocf.write('tlshape      <- %d\n' % tlshape)
            adhocf.write('dirparam     <- %d\n' % dirparam)
            adhocf.write('samplefreq   <- 1\n')
            adhocf.write('printfreq    <- 100\n')
            adhocf.write('datafile     <- "../../../data/data0-%d.nex"\n' % sim_plus_one)
            adhocf.write('treefile     <- "prior.trees"\n')
            adhocf.write('logfile      <- "prior.log"\n')
            adhocf.write('prior_path   <- FALSE\n')
            adhocf.write('sample_prior <- TRUE\n')
            adhocf.close()
            
            # Create end directory
            repsimenddir = os.path.join(repdir, simdir, 'end')
            os.mkdir(repsimenddir)
            
            # Create post directory inside end directory
            repsimendpostdir = os.path.join(repsimenddir, 'post')
            os.mkdir(repsimendpostdir)
            adhocf = open(os.path.join(repsimendpostdir, 'adhoc.Rev'), 'w')
            adhocf.write('rnseed       <- %d\n' % random.randint(1,99999))
            adhocf.write('niter        <- %d\n' % niters)
            adhocf.write('tlshape      <- %d\n' % tlshape)
            adhocf.write('dirparam     <- %d\n' % dirparam)
            adhocf.write('samplefreq   <- 1\n')
            adhocf.write('printfreq    <- 100\n')
            adhocf.write('datafile     <- "../../../data/data-%d.nex"\n' % sim_plus_one)
            adhocf.write('treefile     <- "post.trees"\n')
            adhocf.write('logfile      <- "post.log"\n')
            adhocf.write('prior_path   <- FALSE\n')
            adhocf.write('sample_prior <- FALSE\n')
            adhocf.close()
            
            # Create prior directory inside end directory
            repsimendpriordir = os.path.join(repsimenddir, 'prior')
            os.mkdir(repsimendpriordir)
            adhocf = open(os.path.join(repsimendpriordir, 'adhoc.Rev'), 'w')
            adhocf.write('rnseed       <- %d\n' % random.randint(1,99999))
            adhocf.write('niter        <- %d\n' % niters)
            adhocf.write('tlshape      <- %d\n' % tlshape)
            adhocf.write('dirparam     <- %d\n' % dirparam)
            adhocf.write('samplefreq   <- 1\n')
            adhocf.write('printfreq    <- 100\n')
            adhocf.write('datafile     <- "../../../data/data-%d.nex"\n' % sim_plus_one)
            adhocf.write('treefile     <- "prior.trees"\n')
            adhocf.write('logfile      <- "prior.log"\n')
            adhocf.write('prior_path   <- FALSE\n')
            adhocf.write('sample_prior <- TRUE\n')
            adhocf.close()
            
            # Create calcdiss.sh file inside simdir to calculate dissonance for this simulation
            calcdissf = open(os.path.join(repdir, simdir, 'calcdiss.sh'), 'w')
            calcdissf.write('#!/bin/bash\n')
            calcdissf.write('\n')
            calcdissf.write('calcinfo="%s/../calcinfo.py"\n' % current_working_dir)
            calcdissf.write('\n')
            calcdissf.write('python3 $calcinfo --hpdrad --radpct 95 --dirname results --prefix infostart --skip 1 \\\n')
            calcdissf.write('    --postfile start/post.trees --priorfile start/prior.trees\n')
            calcdissf.write('\n')
            calcdissf.write('python3 $calcinfo --hpdrad --radpct 95 --dirname results --prefix infoend --skip 1 \\\n')
            calcdissf.write('    --postfile end/post.trees --priorfile end/prior.trees\n')
            calcdissf.write('\n')
            calcdissf.write("START_TL=`cat results/infostart/op-post.R | grep -Eo 'tree length = ([-.0-9e]+)' | awk -F ' = ' '{print $2}'`\n")
            calcdissf.write("END_TL=`cat results/infoend/op-post.R | grep -Eo 'tree length = ([-.0-9e]+)' | awk -F ' = ' '{print $2}'`\n")
            calcdissf.write('\n')
            calcdissf.write('# Scale everything to a tree length of 1\n')
            calcdissf.write('START_SCALER=`echo "1.0/$START_TL" | bc -l`\n')
            calcdissf.write('END_SCALER=`echo "1.0/$END_TL" | bc -l`\n')
            calcdissf.write('\n')
            calcdissf.write("START_VAR=`cat results/infostart/op-post.R | grep -Eo 'variance = ([-.0-9e]+)' | awk -F ' = ' '{print $2}'`\n")
            calcdissf.write("END_VAR=`cat results/infoend/op-post.R | grep -Eo 'variance = ([-.0-9e]+)' | awk -F ' = ' '{print $2}'`\n")
            calcdissf.write('START_SD=`echo "$START_SCALER * sqrt($START_VAR)" | bc -l`\n')
            calcdissf.write('END_SD=`echo "$END_SCALER * sqrt($END_VAR)" | bc -l`\n')
            calcdissf.write('\n')
            calcdissf.write('op --reftree ./results/infostart/op-post.tre \\\n')
            calcdissf.write('   --refskip 0 \\\n')
            calcdissf.write('   --refscale $START_SCALER \\\n')
            calcdissf.write('   --treefile ./results/infoend/op-post.tre \\\n')
            calcdissf.write('   --skip 0 \\\n')
            calcdissf.write('   --scale $END_SCALER \\\n')
            calcdissf.write('   --prefix results/meandiff\n')
            calcdissf.write('\n')
            calcdissf.write("MEAN_DIFF=`cat results/meandiff.tre | grep -Eo 'bhvdist = ([-.0-9e]+)' | awk -F ' = ' '{print $2}'`\n")
            calcdissf.write("START_EFFECT_SIZE=`echo \"$MEAN_DIFF / $START_SD\" | bc -l`\n")
            calcdissf.write("END_EFFECT_SIZE=`echo \"$MEAN_DIFF / $END_SD\" | bc -l`\n")
            calcdissf.write('\n')
            calcdissf.write('printf "\\n\\nTree length for start mean tree: %g" "$POST_TL_START"\n')
            calcdissf.write('printf "\\nTree length for end mean tree: %g" "$POST_TL_END"\n')
            calcdissf.write('printf "\\n\\nScaler for start: %g" "$START_SCALER"\n')
            calcdissf.write('printf "\\nScaler for end: %g" "$END_SCALER"\n')
            calcdissf.write('printf "\\n\\nStd. dev. (start): %g" "$START_SD"\n')
            calcdissf.write('printf "\\nStd. dev. (end): %g" "$END_SD"\n')
            calcdissf.write('printf "\\n\\nMean difference: %g" "$MEAN_DIFF"\n')
            calcdissf.write('printf "\\nEffect size (start): %g" "$START_EFFECT_SIZE"\n')
            calcdissf.write('printf "\\nEffect size (end): %g" "$END_EFFECT_SIZE"\n')
            calcdissf.write('printf "\\n"\n')
            calcdissf.write('\n')
            calcdissf.close()
            
    # Create R file for plot
    plotRfn = os.path.join(masterdir, 'bhv-vs-step.R')
    plotf = open(plotRfn, 'w')
    plotf.write('cwd = system(\'cd "$( dirname "$0" )" && pwd\', intern = TRUE)\n')
    plotf.write('setwd(cwd)\n')
    plotf.write('pdf("bhv-vs-step.pdf")\n')
        
    if use_ggplot:
        # Save data used for plotting
        rstr = []
        xstr = []
        ystr = []
        for r in range(nreps):
            n = len(bhvrep[r])
            for s in range(n):
                rstr.append('%d' % (r+1,))
                xstr.append('%g' % bhvrep[r][s][0])
                ystr.append('%g' % bhvrep[r][s][1])
                
        plotf.write('library(ggplot2)\n')
        plotf.write('library(RColorBrewer)\n')
        plotf.write('# https://stackoverflow.com/questions/9563711/r-color-palettes-for-many-data-classes\n')
        plotf.write('c20 <- c(\n')
        plotf.write('  "dodgerblue2",    # red    \n')
        plotf.write('  "#E31A1C",        # red    \n')
        plotf.write('  "green4",         # green  \n')
        plotf.write('  "#6A3D9A",        # purple \n')
        plotf.write('  "#FF7F00",        # orange \n')
        plotf.write('  "black",          # black  \n')
        plotf.write('  "gold1",          # gold   \n')
        plotf.write('  "#CAB2D6",        # purple \n')
        plotf.write('  "#FDBF6F",        # orange \n')
        plotf.write('  "gray70",         # gray   \n')
        plotf.write('  "khaki2",         # brown  \n')
        plotf.write('  "maroon",         # red    \n')
        plotf.write('  "orchid1",        # purple \n')
        plotf.write('  "blue1",          # blue   \n')
        plotf.write('  "steelblue4",     # blue   \n')
        plotf.write('  "darkturquoise",  # purple \n')
        plotf.write('  "darkolivegreen", # green  \n')
        plotf.write('  "yellow4",        # yellow \n')
        plotf.write('  "darkorange4",    # orange \n')
        plotf.write('  "brown"           # brown  \n')
        plotf.write(')\n')
        plotf.write('r <- factor(c(%s))\n' % ','.join(rstr))
        plotf.write('x <- c(%s)\n' % ','.join(xstr))
        plotf.write('y <- c(%s)\n' % ','.join(ystr))
        plotf.write('df <- data.frame(r, x, y)\n')
        plotf.write('ggplot(data = df, aes(x = x, y = y, color = r, group = r)) +\n')
        plotf.write('  scale_color_manual(values = c20) + \n')
        plotf.write('  geom_line() +\n')
        plotf.write('  labs(\n')
        plotf.write('    title = "",\n')
        plotf.write('    x = "Steps",\n')
        plotf.write('    y = "Geodesic distance"\n')
        plotf.write('  ) + \n')
        plotf.write('  coord_cartesian(xlim = c(0.0, %g), ylim = c(0.0, %g)) +\n' % (xmax, ymax))
        plotf.write('  theme_minimal() +\n')
        plotf.write('  theme(legend.position = "none")\n')
    else:
        plotf.write('cols <- terrain.colors(%d)\n' % nreps)
        
        # Save data used for plotting
        for rep in range(nreps):
            rep_plus_one = rep + 1
            repdir = os.path.join(masterdir, 'rep%d' % rep_plus_one)
            
            if rep_plus_one == 1:
                # Create a vector of step indices (same for all replicates)
                tstr = ','.join(['%d' % t for t,d in bhvrep[rep]])
    
                # Write the vector to plotf
                plotf.write('t <- c(%s)\n' % tstr)
    
            # Create a vector of geodesic distances for this replicate
            dstr = ','.join(['%g' % d for t,d in bhvrep[rep]])
            
            # Write the vector to plotf
            plotf.write('d%d <- c(%s)\n' % (rep_plus_one, dstr))
        
        # Add plot command to generate the line for the first replicate
        plotf.write('plot(t, d1, type="l", lwd=2, col=cols[1], ylim=c(0, %g), xlab="Steps", ylab="BHV distance")\n' % ymax)
        
        # Add additional lines commands for overplotting remaining lines
        for rep in range(1,nreps):
            rep_plus_one = rep + 1
            plotf.write('lines(t, d%d, lwd=2, col=cols[%d])\n' % (rep_plus_one, rep_plus_one))
            
    # Close the plot file
    plotf.write('dev.off()\n')
    plotf.close()
    
    # Process R file to create plot
    subprocess.run(['rscript', 'bhv-vs-step.R'], cwd=masterdir)
    
    # Open plot
    subprocess.run(['open', 'bhv-vs-step.pdf'], cwd=masterdir)
    
```
