## Supplementary material for "Estimating Bayesian Phylogenetic Information Content Using Geodesic Distances": slurm-template.sh

```
#!/bin/bash
#SBATCH --job-name=diss                         # Job name
#SBATCH --output=diss%a.out                     # stdout: %a will be replaced by array job number (-o)
#SBATCH --error=diss%a.err                      # stderr: %a will be replaced by array job number (-e)
#SBATCH --array=1-__ARRAY_SIZE__%36             # nreps*nstations analyses (max 36 running simultaneously)
#SBATCH --partition=general                     # Name of Partition
#SBATCH --ntasks=1                              # Maximum CPU cores for each job
#SBATCH --nodes=1                               # Ensure all cores are from the same node for each job
#SBATCH --cpus-per-task=1                       # CPU-cores per task (>1 if multi-threaded tasks) for each job
#SBATCH --constraint='epyc128'                  # Target the AMD Epyc 128-core node architecture
#SBATCH --mem=20G                               # Request 20GB of available RAM
#SBATCH --mail-type=ALL                         # Event(s) that triggers email notification (BEGIN,END,FAIL,ALL)
#SBATCH --mail-user=        # Destination email address

# Set master random number seed
RANDOM=45971

# Load modules needed
module load python/3.12.5

# Specify paths
basedir=`pwd`
RB=$HOME/bin/rb131
OP=$HOME/bin/op194

# Generate arrays containing random number seeds and data file names
# i = $SLURM_ARRAY_TASK_ID - 1
# rep = 1 + i / __SIMS_PER_REP__
# sim = 1 + i % __SIMS_PER_REP__
# $SLURM_ARRAY_TASK_ID    i     rep    sim
#          1              0       1      1
#          2              1       1      2
#          3              2       1      3
#          4              3       2      1
#          5              4       2      2
#          6              5       2      3
rep=()
sim=()
for (( i=0; i<__ARRAY_SIZE__; i++ )); do
    rep+=($(( 1 + $i / __NSIMS__ )))
    k=$(( 1 + $i % __NSIMS__ ))
    sim+=($k)
done

j=$(( $SLURM_ARRAY_TASK_ID - 1 ))
myrep=${rep[$j]}
mysim=${sim[$j]}

echo
echo "SLURM_ARRAY_TASK_ID = $SLURM_ARRAY_TASK_ID"
echo "myrep   = $myrep"
echo "mysim   = $mysim"

echo
echo "************************************************"
echo "***** Run RevBayes for 'starting' data set *****"
echo "************************************************"
cd $basedir/rep$myrep/sim$mysim/start/post
$RB $basedir/fig-dissonance.Rev
cd $basedir/rep$myrep/sim$mysim/start/prior
$RB $basedir/fig-dissonance.Rev

echo
echo "**********************************************"
echo "***** Run RevBayes for 'ending' data set *****"
echo "**********************************************"
cd $basedir/rep$myrep/sim$mysim/end/post
$RB $basedir/fig-dissonance.Rev
cd $basedir/rep$myrep/sim$mysim/end/prior
$RB $basedir/fig-dissonance.Rev
cd $basedir
    
echo
echo "**********************************************"
echo "***** Run calcinfo to compute dissonance *****"
echo "**********************************************"

cd $basedir/rep$myrep/sim$mysim
rm -rf results

python3 $basedir/calcinfo.py --oppath $OP --hpdrad --radpct 95 --dirname results --prefix infoHPDstart --skip 1 \
    --postfile start/post/post.trees --priorfile start/prior/prior.trees

python3 $basedir/calcinfo.py --oppath $OP --hpdrad --radpct 95 --dirname results --prefix infoHPDend --skip 1 \
    --postfile end/post/post.trees --priorfile end/prior/prior.trees

python3 $basedir/calcinfo.py --oppath $OP --radpct 95 --dirname results --prefix infoRADstart --skip 1 \
    --postfile start/post/post.trees --priorfile start/prior/prior.trees

python3 $basedir/calcinfo.py --oppath $OP --radpct 95 --dirname results --prefix infoRADend --skip 1 \
    --postfile end/post/post.trees --priorfile end/prior/prior.trees

cd $basedir
```
