## Supplementary material for "Estimating Bayesian Phylogenetic Information Content Using Geodesic Distances": summarize.py

```
import argparse,sys,re,os,glob,math,subprocess,deploysettings
from scipy.stats import describe,pearsonr

parser = argparse.ArgumentParser(
    formatter_class=argparse.RawDescriptionHelpFormatter,
    description='''Summarizes dissonance for a series of increasingly distant model trees. Specify "--mode hpd" to summarize the HPD results or "--mode rad" to summarize the RAD results.''',
    epilog='''Be sure OPpath is the correct path to the OP executable. OP may be obtained from https://github.com/plewis/OP.''')

parser.add_argument('--oppath',    default='op',                          help='path to OP executable')
parser.add_argument('--mode',      required=True, choices=["hpd", "rad"], help='specify which results to summarize')

settings = vars(parser.parse_args())
oppath = settings['oppath']
mode   = settings['mode']

masterdir = deploysettings.masterdir
nreps     = deploysettings.nreps
nstations = deploysettings.nstations
stations  = deploysettings.stations

current_working_dir = os.getcwd()

# If true, uses ggplot2 to create the plots; 
# otherwise, uses the built-in R plot command
use_ggplot = True

print('mode      = "%s"' % mode)
print('masterdir = "%s"' % masterdir)
print('nreps     = %d' % nreps)
print('nstations = %d' % nstations)
print('stations = ', stations)

def extractStatsFrom(fn):
    if not os.path.exists(fn):
        print('extractStatsFrom failed because file "%s" does not exist' % fn)
        return None
        
    stuff = open(fn, 'r').read()
    
    # Extract sample size
    m = re.search(r'sample size = ([-0-9]+)', stuff, re.S | re.M)
    assert m is not None, 'Failed to extract sample size from file "%s"' % fn
    sample_size = int(m.group(1))
    
    # Extract tree length
    m = re.search(r'tree length = ([-.0-9e]+)', stuff, re.S | re.M)
    assert m is not None, 'Failed to extract tree length from file "%s"' % fn
    tree_length = float(m.group(1))
    
    # Extract variance
    m = re.search(r'variance = ([-.0-9e]+)', stuff, re.S | re.M)
    assert m is not None, 'Failed to extract variance from file "%s"' % fn
    variance = float(m.group(1))
    assert variance >= 0.0, 'variance was negative for "%s"' % fn
    
    # Extract radius
    m = re.search(r'95% radius = ([-.0-9e]+)', stuff, re.S | re.M)
    assert m is not None, 'Failed to extract radius from file "%s"' % fn
    radius = float(m.group(1))
    assert radius >= 0.0, 'radius was negative for "%s"' % fn
        
    # Calculate scaler
    assert tree_length >= 0.0, 'tree length was negative for "%s"' % fn
    scaler = 1.0/tree_length
    
    # Calculate standard deviation
    stddev = math.sqrt(variance)

    return {'sample_size':sample_size, 'tree_length':tree_length, 'variance':variance, 'radius':radius, 'scaler':scaler, 'stddev':stddev}   

def extractMeanDiffFrom(fn):
    stuff = open(fn, 'r').read()
    m = re.search(r'bhvdist = ([-.0-9e]+)', stuff, re.S | re.M)
    assert m is not None, 'Failed to extract bhvdist from file "%s"' % fn
    return float(m.group(1))

def createPlot(use_radius):
    # Start by gathering statistics that will be saved as comments in the R plot file
    # These statistics include the mean and standard deviation of the ending BHV
    # distance, the starting dissonance, and the ending dissonance, as well as the 
    # number of replicates (and segments) that were monotoically increasing
    ystartingdiss = []
    yendingdiss = []
    nmonotonic = 0
    nmonosegments = 0
    totalsegments = 0
    distances = []
    dissonances = []
    nexcludedreps = len(excluded_reps)
    for rep in range(nreps):
        if not rep in excluded_reps:
            is_monotonic = True
            if use_radius:
                ystartingdiss.append(replicate[rep]['yradius'][0])
                yendingdiss.append(replicate[rep]['yradius'][nstations - 1])
                yprev = replicate[rep]['yradius'][0]
                distances.append(replicate[rep]['x'][0])
                dissonances.append(replicate[rep]['yradius'][0])
                for station in range(1,nstations):
                    distances.append(replicate[rep]['x'][station])
                    dissonances.append(replicate[rep]['yradius'][station])
                    ycurr = replicate[rep]['yradius'][station]
                    if ycurr is not None:
                        totalsegments += 1
                        if ycurr >= yprev:
                            nmonosegments += 1
                        else:
                            is_monotonic = False
                        yprev = ycurr
                if is_monotonic:
                    nmonotonic += 1
            else:
                ystartingdiss.append(replicate[rep]['ystddev'][0])
                yendingdiss.append(replicate[rep]['ystddev'][nstations - 1])
                yprev = replicate[rep]['ystddev'][0]
                distances.append(replicate[rep]['x'][0])
                dissonances.append(replicate[rep]['ystddev'][0])
                for station in range(1,nstations):
                    distances.append(replicate[rep]['x'][station])
                    dissonances.append(replicate[rep]['ystddev'][station])
                    ycurr = replicate[rep]['ystddev'][station]
                    if ycurr is not None:
                        totalsegments += 1
                        if ycurr >= yprev:
                            nmonosegments += 1
                        else:
                            is_monotonic = False
                        yprev = ycurr
                if is_monotonic:
                    nmonotonic += 1
                    
    # Calculate correlation between distances and dissonances
    assert len(distances) == len(dissonances)
    xycorr = pearsonr(distances, dissonances)
    distance_dissonance_correlation = xycorr.statistic
    distance_dissonance_pvalue = xycorr.pvalue
    distance_dissonance_n = len(distances)
    
    ystartingdescribe = describe(ystartingdiss, 0, 1)  # 2nd arg 0 is axis
    ystartingmean = ystartingdescribe.mean
    ystartingsd = math.sqrt(ystartingdescribe.variance)
    
    yendingdescribe = describe(yendingdiss, 0, 1)  # 2nd arg 0 is axis
    yendingmean = yendingdescribe.mean
    yendingsd = math.sqrt(yendingdescribe.variance)
    
    percent_segments_monotonic = 100.0*nmonosegments/totalsegments
    
    if use_radius:
        harvestRfn = os.path.join(masterdir, 'diss-vs-bhv-%s-radius.R' % mode)
        harvestPDFfn = os.path.join(masterdir, 'diss-vs-bhv-%s-radius.pdf' % mode)
    else:
        harvestRfn = os.path.join(masterdir, 'diss-vs-bhv-%s-stddev.R' % mode)
        harvestPDFfn = os.path.join(masterdir, 'diss-vs-bhv-%s-stddev.pdf' % mode)
    harvestf = open(harvestRfn, 'w')
    if use_radius:
        harvestf.write('# Dissonance: %s-radius\n' % mode)
    else:
        harvestf.write('# Dissonance: %s-stddev\n' % mode)
    harvestf.write('# %d excluded replicates\n' % nexcludedreps)
    if nexcludedreps > 0:
        for i in range(nexcludedreps):
            harvestf.write('# rep %d was excluded\n' % excluded_reps[i])
    harvestf.write('# %.9f pooled-radius/pooled-stddev\n' % (pooled_radius/pooled_stddev,))
    harvestf.write('# %.9f mean ending distance\n' % xmean)
    harvestf.write('# %.9f sd ending distance\n' % xsd)
    harvestf.write('# %.9f mean starting dissonance\n' % ystartingmean)
    harvestf.write('# %.9f sd starting dissonance\n' % ystartingsd)
    harvestf.write('# %.9f mean ending dissonance\n' % yendingmean)
    harvestf.write('# %.9f sd ending dissonance\n' % yendingsd)
    harvestf.write('# %d curves are monotonically increasing\n' % nmonotonic)
    harvestf.write('# %d (%.5f %%) segments (of %d total) are increasing\n' % (nmonosegments, percent_segments_monotonic, totalsegments))
    harvestf.write('# %.9f correlation between distance and dissonance\n' % distance_dissonance_correlation)
    harvestf.write('#   %.9f pvalue\n' % distance_dissonance_pvalue)
    harvestf.write('#   %d sample size\n' % distance_dissonance_n)
    harvestf.write('# %15s %15s %15s %15s %15s\n' % ('Replicate', 'Station', 'Distance', 'Dissonance', 'Change'))
    for r in range(nreps):
        if not r in excluded_reps:
            for s in range(nstations):
                if use_radius:
                    if s == 0:
                        harvestf.write('# %15d %15d %15.5f %15.5f %15s\n' % (r, s, replicate[r]['x'][s], replicate[r]['yradius'][s], '---'))
                    else:
                        harvestf.write('# %15d %15d %15.5f %15.5f %15.5f\n' % (r, s, replicate[r]['x'][s], replicate[r]['yradius'][s], replicate[r]['yradius'][s] - replicate[r]['yradius'][s-1]))
                else:
                    if s == 0:
                        harvestf.write('# %15d %15d %15.5f %15.5f %15s\n' % (r, s, replicate[r]['x'][s], replicate[r]['ystddev'][s], '---'))
                    else:
                        harvestf.write('# %15d %15d %15.5f %15.5f %15.5f\n' % (r, s, replicate[r]['x'][s], replicate[r]['ystddev'][s], replicate[r]['ystddev'][s] - replicate[r]['ystddev'][s-1]))
    harvestf.write('cwd = system(\'cd "$( dirname "$0" )" && pwd\', intern = TRUE)\n')
    harvestf.write('setwd(cwd)\n')
    harvestf.write('pdf("%s")\n' % harvestPDFfn)
    
    if use_ggplot:
        rstr = []
        sstr = []
        xstr = []
        ystr = []
        for r in range(nreps):
            if not r in excluded_reps:
                for s in range(nstations):
                    rstr.append('%d' % (r+1,))
                    sstr.append('%g' % stations[s])
                    xstr.append('%g' % replicate[r]['x'][s])
                    if use_radius:
                        ystr.append('%g' % replicate[r]['yradius'][s])
                    else:
                        ystr.append('%g' % replicate[r]['ystddev'][s])
                    
        harvestf.write('library(ggplot2)\n')
        harvestf.write('library(RColorBrewer)\n')
        harvestf.write('# https://stackoverflow.com/questions/9563711/r-color-palettes-for-many-data-classes\n')
        harvestf.write('c20 <- c(\n')
        harvestf.write('  "dodgerblue2",    # red    \n')
        harvestf.write('  "#E31A1C",        # red    \n')
        harvestf.write('  "green4",         # green  \n')
        harvestf.write('  "#6A3D9A",        # purple \n')
        harvestf.write('  "#FF7F00",        # orange \n')
        harvestf.write('  "black",          # black  \n')
        harvestf.write('  "gold1",          # gold   \n')
        harvestf.write('  "#CAB2D6",        # purple \n')
        harvestf.write('  "#FDBF6F",        # orange \n')
        harvestf.write('  "gray70",         # gray   \n')
        harvestf.write('  "khaki2",         # brown  \n')
        harvestf.write('  "maroon",         # red    \n')
        harvestf.write('  "orchid1",        # purple \n')
        harvestf.write('  "blue1",          # blue   \n')
        harvestf.write('  "steelblue4",     # blue   \n')
        harvestf.write('  "darkturquoise",  # purple \n')
        harvestf.write('  "darkolivegreen", # green  \n')
        harvestf.write('  "yellow4",        # yellow \n')
        harvestf.write('  "darkorange4",    # orange \n')
        harvestf.write('  "brown"           # brown  \n')
        harvestf.write(')\n')
        harvestf.write('r = factor(c(%s))\n' % ','.join(rstr))
        #harvestf.write('s = c(%s)\n' % ','.join(sstr))
        harvestf.write('x = c(%s)\n' % ','.join(xstr))
        harvestf.write('y = c(%s)\n' % ','.join(ystr))
        #harvestf.write('df <- data.frame(r, s, x, y)\n')
        harvestf.write('df <- data.frame(r, x, y)\n')
        harvestf.write('ggplot(data = df, aes(x = x, y = y, color = r, group = r)) +\n')
        harvestf.write('  scale_color_manual(values = c20) + \n')
        harvestf.write('  geom_line() +\n')
        harvestf.write('  geom_point() +\n')
        harvestf.write('  labs(\n')
        harvestf.write('    title = "",\n')
        harvestf.write('    x = "Geodesic Distance",\n')
        if use_radius:
            harvestf.write('    y = "Dissonance"\n')
        else:
            harvestf.write('    y = "Dissonance"\n')
        harvestf.write('  ) + \n')
        if use_radius:
            harvestf.write('  coord_cartesian(xlim = c(0.0, %g), ylim = c(0.0, %g)) +\n' % (xmax, ymax_radius))
        else:
            harvestf.write('  coord_cartesian(xlim = c(0.0, %g), ylim = c(0.0, %g)) +\n' % (xmax, ymax_stddev))
        harvestf.write('  theme_minimal(base_size = 28) +\n')
        harvestf.write('  theme(legend.position = "none")\n')
    else:
        harvestf.write('cols <- terrain.colors(%d)\n' % nreps)
        
        xstr = ['%g' % x for x in replicate[0]['x']]
        harvestf.write('x0 = c(%s)\n' % ','.join(xstr))
        
        if use_radius:
            ystr = ['%g' % y for y in replicate[0]['yradius']]
        else:
            ystr = ['%g' % y for y in replicate[0]['ystddev']]
        harvestf.write('y0 = c(%s)\n' % ','.join(ystr))
        
        if use_radius:
            harvestf.write('plot(x0, y0, type="b", pch=19, lwd=2, col=cols[1], xlim=c(0.0, %g), ylim=c(0.0, %g), main="", xlab="BHV distance", ylab="Dissonance")\n' % (xmax, ymax))
        else:
            harvestf.write('plot(x0, y0, type="b", pch=19, lwd=2, col=cols[1], xlim=c(0.0, %g), ylim=c(0.0, %g), main="", xlab="BHV distance", ylab="Dissonance")\n' % (xmax, ymax))
        
        print('rep = %d' % 0)
        print(replicate[0]['x'])
        if use_radius:
            print(replicate[0]['yradius'])
        else:
            print(replicate[0]['ystddev'])
        
        for r in range(1, nreps):
            if not r in excluded_reps:
                print('~~~~~~~~~~~~')
                print('rep = %d' % r)
                print(replicate[r]['x'])
                if use_radius:
                    print(replicate[r]['yradius'])
                else:
                    print(replicate[r]['ystddev'])
                
                xstr = ['%g' % x for x in replicate[r]['x']]
                harvestf.write('x%d = c(%s)\n' % (r, ','.join(xstr)))
            
                if use_radius:
                    ystr = ['%g' % y for y in replicate[r]['yradius']]
                else:
                    ystr = ['%g' % y for y in replicate[r]['ystddev']]
                harvestf.write('y%d = c(%s)\n' % (r, ','.join(ystr)))
            
                harvestf.write('lines(x%d, y%d, lwd=2, col=cols[%d])\n' % (r, r, r+1))
                harvestf.write('points(x%d, y%d, pch=19, col=cols[%d])\n' % (r, r, r+1))
        
    harvestf.write('dev.off()\n')
    harvestf.close()
    
    subprocess.run(['rscript', harvestRfn])

if __name__ == '__main__':
    if mode == 'rad':
        info_start_dir = 'infoRADstart'
        info_end_dir   = 'infoRADend'
    else:
        info_start_dir = 'infoHPDstart'
        info_end_dir   = 'infoHPDend'
    
    # Get a list of all the rep* directories inside masterdir
    repdirs = sorted(glob.glob(os.path.join(masterdir,'rep*')))
    
    # Stores dissonance statistics
    dissonance = []
    
    # Keep track of replicates excluded because not all stations were completed
    excluded_reps = []
    
    # Open file to store details
    outf = open(os.path.join(masterdir, 'dissonance', 'details-%s.txt' % mode), 'w')
    outf.write('rep\tsim\tmode\tcohens_stddev\tcohens_radius\tmean_diff\tpooled_radius\tpooled_stddev\tsd_start\tsd_end\tn_start\tn_end\ttl_start\ttl_end\tscaler_start\tscaler_end\n')
    
    # Visit each rep* directory
    for repdir in repdirs:
        print(repdir)
        
        # For each directory in repdirs, get a list of each sim* directory
        simdirs = sorted(glob.glob(os.path.join(repdir,'sim*')))
        
        # Count number of stations that failed
        nfails = 0
        
        # Visit each sim* directory
        for simdir in simdirs:
            # Extract replicate number and simulation number from directory name
            m = re.search(r'rep(\d+)/sim(\d+)', simdir)
            assert m is not None, 'Failed to extract replicate and simulation number from "%s"' % simdir
            rep = int(m.group(1)) - 1
            sim = int(m.group(2)) - 1
    
            print('  %s (rep = %d, sim = %d)' % (simdir, rep, sim))
            
            # Get starting sample size, tree length, variance
            stats_start = extractStatsFrom(os.path.join(simdir, 'results', info_start_dir, 'op-post.R'))
            
            # Get ending sample size, tree length, variance
            stats_end = extractStatsFrom(os.path.join(simdir, 'results', info_end_dir, 'op-post.R'))
    
            if stats_start is None or stats_end is None:
                nHPDfails += 1
            else:
                n_start      = stats_start['sample_size']
                tl_start     = stats_start['tree_length']
                radius_start = stats_start['radius']
                scaler_start = stats_start['scaler']
                stddev_start = stats_start['stddev']
                sd_start     = scaler_start*stddev_start
                
                n_end      = stats_end['sample_size']
                tl_end     = stats_end['tree_length']
                radius_end = stats_end['radius']
                scaler_end = stats_end['scaler']
                stddev_end = stats_end['stddev']
                sd_end     = scaler_end*stddev_end
                
                var_start = math.pow(sd_start, 2)
                var_end   = math.pow(sd_end, 2)
                
                # Calculated the pooled standard deviation
                pooled_numer = var_start*(n_start - 1) + var_end*(n_end - 1)
                pooled_denom = n_start + n_end - 2
                pooled_stddev = math.sqrt(pooled_numer/pooled_denom)
                
                # Calculated the pooled radius
                pooled_numer = radius_start*(n_start - 1) + radius_end*(n_end - 1)
                pooled_denom = n_start + n_end - 2
                pooled_radius = math.sqrt(pooled_numer/pooled_denom)
                
                cmds = [oppath,
                    '--reftree', os.path.join(simdir, 'results', 'infoHPDstart', 'op-post.tre'),
                    '--refskip', '0',
                    '--refscale', '%g' % scaler_start,
                    '--treefile', os.path.join(simdir, 'results', 'infoHPDend', 'op-post.tre'),       
                    '--skip', '0',
                    '--scale', '%g' % scaler_end,
                    '--prefix', os.path.join(simdir, 'results', 'meanHPDdiff')
                ]
                completed_process = subprocess.run(cmds, cwd=current_working_dir, capture_output=True)
                try:
                    completed_process.check_returncode()
                except sub.CalledProcessError:
                    print('\nAn error occurred while running OP for "%s"' % simdir)
                    print('Standard error:')
                    print(completed_process.stderr)
                    print('Standard output:')
                    print(completed_process.stdout)
                    sys.exit('Aborted.')
                
                mean_diff = extractMeanDiffFrom(os.path.join(simdir, 'results', 'meanHPDdiff.tre'))
                cohens_stddev = mean_diff/pooled_stddev
                cohens_radius = mean_diff/pooled_radius
                
                # Write details to the details file
                outf.write('%d\t%d\t%s\t%g\t%g\t%g\t%g\t%g\t%g\t%g\t%d\t%d\t%g\t%g\t%g\t%g\n' % 
                (rep, sim, mode, cohens_stddev, cohens_radius, mean_diff, pooled_radius, pooled_stddev, sd_start, sd_end, n_start, n_end, tl_start, tl_end, scaler_start, scaler_end))
                
                # Create dissonance entry
                dissmap = {
                    'rep':rep,
                    'sim':sim,
                    'mode':mode,
                    'cohens_stddev':cohens_stddev,
                    'cohens_radius':cohens_radius,
                    'mean_diff':mean_diff,
                    'pooled_radius':pooled_radius,
                    'pooled_stddev':pooled_stddev,
                    'n_start':n_start,
                    'n_end':n_end,
                    'tl_start':tl_start,
                    'tl_end':tl_end,
                    'scaler_start':scaler_start,
                    'scaler_end':scaler_end,
                    'stddev_start':stddev_start,
                    'stddev_end':stddev_end,
                    'sd_start':sd_start,
                    'sd_end':sd_end
                }
                
                dissonance.append(dissmap)
                
        if nfails > 0:
            excluded_reps.append(rep)     
    
    outf.close()
    
    if len(excluded_reps) > 0:        
        print('Replicates excluded because at least one station failed:')           
        for r in excluded_reps:
            print('  %d' % r)
        input('Press return to continue...')
        
    # Read true BHV distances between model trees
    # These values will form the x axis of the plot
    bhvdists = []
    bhv_filenames = glob.glob(os.path.join(masterdir,'bhvdist', 'bhv*.txt'))
    assert len(bhv_filenames) > 0, 'Oops, not expecting bhv_filenames to be empty'
    for fn in bhv_filenames:
        lines = open(fn, 'r').readlines()
        headers = lines[0].strip().split('\t')
        content = lines[1].strip().split('\t')
        
        # bhvmap has these keys:
        #  'rep'     = replicate
        #  'sim'     = station along random walk
        #  'address' = iteration this station
        #  'bhv'     = BHV distance between start and this station
        assert len(headers) == 4
        assert headers[0] == 'rep'
        assert headers[1] == 'sim'
        assert headers[2] == 'address'
        assert headers[3] == 'bhv'
        assert len(content) == 4
        rep     = int(content[0]) - 1
        assert rep >= 0
        sim     = int(content[1]) - 1
        assert sim >= 0
        address = int(content[2])
        bhv     = float(content[3])
        
        bhvmap = {}
        bhvmap['rep']     = rep
        bhvmap['sim']     = sim
        bhvmap['address'] = address
        bhvmap['bhv']     = bhv
        bhvdists.append(bhvmap)
    
    # Create the replicate array with slots for nstations x, ysd, and yrad values
    # replicate[rep][station]['x'] stores the BHV distance between the start and this station
    # replicate[rep][station]['ystddev'] stores the dissonance in pooled standard deviation units
    # replicate[rep][station]['yradius'] stores the dissonance in pooled radius units
    replicate = []
    for r in range(nreps):
        replicate.append({'x': [None]*nstations, 'ystddev': [None]*nstations, 'yradius': [None]*nstations})
    
    # Populate the x values in the replicate array    
    assert len(bhvdists) > 0, 'Oops, not expecting bhvdists to be empty'
    xmin = None
    xmax = None
    for b in bhvdists:
        rep = b['rep']
        sim = b['sim']
        bhv = b['bhv']
        if xmin is None or bhv < xmin:
            xmin = bhv
        if xmax is None or bhv > xmax:
            xmax = bhv
        replicate[rep]['x'][sim] = bhv
    
    # Calculate the total distance between the start and the last station
    xtotaldist = []
    for rep in range(nreps):
        xtotaldist.append(replicate[rep]['x'][nstations - 1])
    
    # Calculate the mean and standard deviation of the nreps total distances
    xdescribe = describe(xtotaldist, 0, 1)  # 2nd arg 0 is axis
    xmean = xdescribe.mean
    xsd = math.sqrt(xdescribe.variance)
    
    # Populate the ystdev and yradius values in the replicate array    
    ymin_stddev = None
    ymax_stddev = None
    ymin_radius = None
    ymax_radius = None
    for d in dissonance:
        rep = d['rep']
        sim = d['sim']
        cohens_stddev = d['cohens_stddev']
        cohens_radius = d['cohens_radius']
        if ymin_stddev is None or cohens_stddev < ymin_stddev:
            ymin_stddev = cohens_stddev
        if ymax_stddev is None or cohens_stddev > ymax_stddev:
            ymax_stddev = cohens_stddev
        if ymin_radius is None or cohens_radius < ymin_radius:
            ymin_radius = cohens_radius
        if ymax_radius is None or cohens_radius > ymax_radius:
            ymax_radius = cohens_radius
        replicate[rep]['ystddev'][sim] = cohens_stddev
        replicate[rep]['yradius'][sim] = cohens_radius
                
    createPlot(True)   # use_radius = True
    createPlot(False)  # use_radius = False
```
