## Supplementary material for "Estimating Bayesian Phylogenetic Information Content Using Geodesic Distances": _readme.txt

To create fig-saturation:

Note: sites 1-3 and 1405-. were excluded from the original rbcl data file to avoid
including the start codon and trailing missing data. Also, only 21 of the original 43
taxa were included to facilitate MCMC convergence.

Directory layout showing only directories and files mentioned in the instructions below:

  supplementary-materials
      calcinfo.py
      4-fig-saturation
          data
              rbcl.nex
              rbcl1st.nex
              rbcl2nd.nex
              rbcl3rd.nex
          iqtree
              iqtree.sh
              partition.nex
*             partition.nex.best_model.nex
*             partition.nex.bionj
*             partition.nex.ckp.gz
*             partition.nex.iqtree
*             partition.nex.log
*             partition.nex.mldist
*             partition.nex.treefile
*             truetree.tre
          phylomad
              commandLineFile.Rscript
*             saturation.test.results.csv
*         reps
*             rep0
*             rep1
*             rep2
*             rep3
*             rep4
*             rep5
*             rep6
*             rep7
*             rep8
*             rep9
*             rep10
*             calcinfo.py
*             go.sh
          seqgen
              seqgen.sh
              concat.py
          _readme.txt
          fig-saturation.Rev
          make-slurm.py
*         output.txt          
          slurm-template.sh
          summarize.py

* file/directory created as a result of these instructions

1.  Run IQ-TREE on data/rbcl.nex to estimate the model and relative rates 
    for the three codon-specific subsets. 
    
    cd supplementary-materials/4-fig-saturation/iqtree
    . iqtree.sh
    
    The file partition.nex.iqtree contains
    the estimated relative rates, kappa values, and nucleotide frequencies for
    each partition subset.
    
    ID  Model           Speed  Parameters
     1  HKY+F          0.3313  HKY{2.14183}+F{0.236464,0.191802,0.388702,0.183033}
     2  HKY+F          0.1454  HKY{1.95213}+F{0.301213,0.238401,0.207913,0.252473}
     3  HKY+F          2.5233  HKY{7.54411}+F{0.3005,0.155705,0.123789,0.420006}
   
2.  Edit the seqgen/seqgen.sh file, replacing the three seq-gen commands in the for
    loop to specify the estimated tratios, base frequencies, and relative rates estimated
    by IQ-TREE. Note that IQ-TREE reports kappa but Seq-Gen requires tratio, so
    conversion from kappa+frequencies to tratio is necessary. Run Seg-Gen to 
    simulate sim1st.nex, sim2nd.nex, and sim3rd.nex.
    
    cd supplementary-materials/4-fig-saturation/seqgen
    chmod +x seqgen.sh
    ./seqgen.sh
    
    This creates the directory "reps" containing one directory for analyzing
    the empirical data (rep0) and 10 directories (rep1...rep10) for analyzing 
    simulated data.
             
3. Edit the slurm-template.sh and adapt the SBATCH commands at the top to your
    cluster. Be sure the paths to the RevBayes and OP executables are correct.

    If you do not have RevBayes installed, here is where you can get it:
        RevBayes 1.3.1:   https://revbayes.github.io/download
        OP 1.9.4:         https://plewis.github.io/op
        
    Run make-slurm.py to create a sim-slurm.sh file within each of
    the simulation directories and a sat-slurm.sh file inside rep0:
    
    cd supplementary-materials/4-fig-saturation
    python3 make-slurm.py
        
4. Specify "hpc" as an alias to your remote cluster in  ~/.ssh/config and
   create a "work/sat" directory inside your home directory on your cluster
   
   ssh hpc
    
5. Upload files to the "work" directories on your cluster:

    cd supplementary-materials/4-fig-saturation
    rsync -azPv reps hpc:work
    scp ../calcinfo.py  hpc:work/reps
    
6. Run the RevBayes and OP analyses on the cluster for both the empirical and
   simulated data:

    ssh hpc
    cd ~/work/reps
    . go.sh

   This will run RevBayes and create a directory "rbcl" with subdirectories
   "prior" and "posterior" containing MCMC samples. It will also run 
   calcinfo.py, which runs OP to calculate information content. It will do 
   these things both for the original data (rep0 directory) and the simulated
   data (rep1..rep10 directories).
    
7. Run summarize.py to compile the information results into tables
   that are saved in the file output.txt

    cd supplementary-materials/4-fig-saturation
    scp summarize.py hpc:work
    ssh hpc
    cd ~/work
    python3 summarize.py
    
8. Run phylomad/commandLineFile.Rscript to perform saturation analyses on empirical
   as well as simulated data sets (all sites as well as 1st-, 2nd-, and 3rd-codon sites)

    cd supplementary-materials/4-fig-saturation/phylomad
    Rscript commandLineFile.Rscript
    
9. Bring results back to your local computer (if desired)

    cd supplementary-materials/4-fig-saturation
    rsync -azPv hpc:work/reps/ reps/
