## Supplementary material for "Estimating Bayesian Phylogenetic Information Content Using Geodesic Distances": make-slurm.py

```
import os,re,glob

# Import the contents of the slurm template
slurm_contents = open(os.path.join('slurm-template.sh'), 'r').read()

# Get a list of all the replicated directories inside the "reps" directory
glob_pattern = os.path.join('reps', 'rep*')
repdirs = glob.glob(glob_pattern)

# Open a master run script that will start all analyses
gof = open(os.path.join('reps', 'go.sh'), 'w')
gof.write('#!/bin/bash\n\n')

for dirname in repdirs:
    # Get the replicate index from dirname
    m = re.match(r'reps/rep(\d+)', dirname)
    assert m is not None, 'Could not extract replicate index from dirname "%s"' % dirname
    i = int(m.group(1))
    
    print('Working on replicate %d...' % i)
    
    if i == 0:    
        # Create sat-slurm.sh to analyze the empirical data
        
        gof.write('cd rep%d; sbatch sat-slurm.sh; cd ..\n' % i)
        
        slurmf = open(os.path.join(dirname, 'sat-slurm.sh'), 'w')
        
        sat_slurm, nsub = re.subn('__JOBNAME__', 'sat', slurm_contents, re.M | re.S)
        assert nsub == 3, 'Expecting 3 subs, found $d' % nsub
        
        sat_slurm, nsub = re.subn('__DATAFIRST__', 'rbcl1st', sat_slurm, re.M | re.S)
        assert nsub == 1, 'Expecting 1 sub, found %d' % nsub
        
        sat_slurm, nsub = re.subn('__DATASECOND__', 'rbcl2nd', sat_slurm, re.M | re.S)
        assert nsub == 1, 'Expecting 1 sub, found %d' % nsub
        
        sat_slurm, nsub = re.subn('__DATATHIRD__', 'rbcl3rd', sat_slurm, re.M | re.S)
        assert nsub == 1, 'Expecting 1 sub, found %d' % nsub
        
        sat_slurm, nsub = re.subn('__DATAALL__', 'rbcl', sat_slurm, re.M | re.S)
        assert nsub == 1, 'Expecting 1 sub, found %d' % nsub
        
        slurmf.write(sat_slurm)
        slurmf.close()
        
    else:
        # Create a sim-slurm.sh file to analyze simulated data
        
        gof.write('cd rep%d; sbatch sim-slurm.sh; cd ..\n' % i)
        
        slurmf = open(os.path.join(dirname, 'sim-slurm.sh'), 'w')
        
        sim_slurm, nsub = re.subn('__JOBNAME__', 'sim%d' % i, slurm_contents, re.M | re.S)
        assert nsub == 3, 'Expecting 3 subs, found $d' % nsub
        
        sim_slurm, nsub = re.subn('__DATAFIRST__', 'sim1st', sim_slurm, re.M | re.S)
        assert nsub == 1, 'Expecting 1 sub, found %d' % nsub
        
        sim_slurm, nsub = re.subn('__DATASECOND__', 'sim2nd', sim_slurm, re.M | re.S)
        assert nsub == 1, 'Expecting 1 sub, found %d' % nsub
        
        sim_slurm, nsub = re.subn('__DATATHIRD__', 'sim3rd', sim_slurm, re.M | re.S)
        assert nsub == 1, 'Expecting 1 sub, found %d' % nsub
        
        sim_slurm, nsub = re.subn('__DATAALL__', 'sim', sim_slurm, re.M | re.S)
        assert nsub == 1, 'Expecting 1 sub, found %d' % nsub
        
        slurmf.write(sim_slurm)
        slurmf.close()

gof.close()
```
