## Supplementary material for "Estimating Bayesian Phylogenetic Information Content Using Geodesic Distances": concat.py

```
import argparse, re, os

parser = argparse.ArgumentParser(
    prog='concat',
    formatter_class=argparse.RawDescriptionHelpFormatter,
    description='''Concatenates the specified nexus-formatted input data sets to create a concatenated nexus-formatted output data set.''',
    epilog='''Assumes that all input data sets have the same taxa in the same order!''')

parser.add_argument('--in', action='append', help='the name of an input data set in nexus format')
parser.add_argument('--out', required=True, help='the desired name of the output data set')

settings = vars(parser.parse_args())

infilenames  = settings['in']
outfilename = settings['out']

print('out = %s' % outfilename)
if infilenames is None:
    sys.exit('Must specify at least one input file name')
else:
    print('in:')
    for fn in infilenames:
        print('  %s' % fn)

def readNexusFile(fn):
    '''
    Reads nexus file whose name is specified by fn and returns ntax, nchar, taxa, and a
    sequences dictionary with taxon names as keys. The values ntax and nchar are integers,
    while taxa is a list of taxon names in the order they were found in the taxa block or
    data block. Any underscores in taxon names are converted to spaces before being saved
    in the taxa list or as a key in the sequences dictionary. Also all nexus comments
    (text in square brackets) will be ignored.
    '''
    stuff = open(fn, 'r').read()
    mask = None

    # determine if taxa block exists
    taxa_block = None
    m = re.search(r'(?:BEGIN|Begin|begin)\s+(?:TAXA|Taxa|taxa)\s*;(.+?)(?:END|End|end)\s*;', stuff, re.M | re.S)
    if m is not None:
        taxa_block = m.group(1).strip()

    # determine if characters block exists
    characters_block = None
    m = re.search(r'(?:BEGIN|Begin|begin)\s+(?:CHARACTERS|Characters|characters)\s*;(.+?)(?:END|End|end)\s*;', stuff, re.M | re.S)
    if m is not None:
        characters_block = m.group(1).strip()

    # determine if data block exists
    data_block = None
    m = re.search(r'(?:BEGIN|Begin|begin)\s+(?:DATA|Data|data)\s*;(.+?)(?:END|End|end)\s*;', stuff, re.M | re.S)
    if m is not None:
        data_block = m.group(1).strip()

    if data_block is not None:
        # get ntax and nchar
        m = re.search(r'(?:DIMENSIONS|dimensions|Dimensions)\s+(?:NTAX|ntax|Ntax|NTax)\s*=\s*(\d+)\s+(?:NCHAR|nchar|Nchar|NChar)\s*=\s*(\d+)\s*;', data_block, re.M | re.S)
        assert m, 'Could not decipher dimensions statement in data block'
        ntax = int(m.group(1))
        nchar = int(m.group(2))

        # get matrix
        m = re.search(r'(?:MATRIX|matrix|Matrix)\s+(.+?)\s*;', data_block, re.M | re.S)
        assert m, 'Could not decipher matrix statement in data block'
        lines = m.group(1).strip().split('\n')
        taxa = []
        sequences = {}
        for line in lines:
            m = re.match(r'\[([-*]+)\]', line.strip())
            if m is not None:
                mask = m.group(1)
            else:
                stripped_line = re.sub(r'\[.+?\]', '', line).strip()
                if len(stripped_line) > 0:
                    parts = line.split()
                    assert len(parts) == 2, 'Found more than 2 parts to this line:\n%s' % line
                    taxon_name = re.sub(r'_', ' ', parts[0]).strip()
                    taxa.append(taxon_name)
                    sequences[taxon_name] = parts[1]
    else:
        assert characters_block is not None and taxa_block is not None, 'Assuming nexus file contains either a data block or a taxa block and characters block'

        # get ntax from taxa block
        m = re.search(r'(?:DIMENSIONS|dimensions|Dimensions)\s+(?:NTAX|ntax|Ntax|NTax)\s*=\s*(\d+)\s*;', taxa_block, re.M | re.S)
        assert m, 'Could not decipher dimensions statement in taxa block'
        ntax = int(m.group(1))

        # get nchar from characters block
        m = re.search(r'(?:DIMENSIONS|dimensions|Dimensions)\s+(?:NCHAR|nchar|Nchar|NChar)\s*=\s*(\d+)\s*;', characters_block, re.M | re.S)
        assert m, 'Could not decipher dimensions statement in characters block'
        nchar = int(m.group(1))

        # get matrix from characters block
        m = re.search(r'(?:MATRIX|matrix|Matrix)\s+(.+?)\s*;', characters_block, re.M | re.S)
        assert m, 'Could not decipher matrix statement in characters block'
        lines = m.group(1).strip().split('\n')
        taxa = []
        sequences = {}
        for line in lines:
            m = re.match(r'\[([-*]+)\]', line.strip())
            if m is not None:
                mask = m.group(1)
            else:
                stripped_line = re.sub(r'\[.+?\]', '', line).strip()
                if len(stripped_line) > 0:
                    parts = stripped_line.split()
                    assert len(parts) == 2, 'Found more than 2 parts to this line:\n%s' % line
                    taxon_name = re.sub(r'_', ' ', parts[0]).strip()
                    taxa.append(taxon_name)
                    sequences[taxon_name] = parts[1]

    return (ntax, nchar, mask, taxa, sequences)

def writeNexusFile(outfn, infilenames, input_dict):
    # Make sure we do not overwrite an existing output file
    if os.path.exists(outfn):
        sys.exit('Please move or rename the file "%s"' % outfn)
        
    # Combine the sequences
    numtax  = None
    numchar = None
    taxon_list = None
    sequences = None
    for fn in infilenames:
        ntax, nchar, mask, taxa, seq = input_dict[fn]
        
        if numtax is None:
            numtax = ntax
        else:
            assert ntax == numtax, 'number of taxa specified in the file "%s" differs from other files' % fn
            
        if numchar is None:
            numchar = nchar
        else:
            numchar += nchar
        
        if taxon_list is None:
            taxon_list = taxa[:]
        else:
            # Check to ensure taxa are the same and in the same order
            for i in range(numtax):
                if taxon_list[i] != taxa[i]:
                    sys.exit('taxon "%s" in file "%s" differs from corresponding taxon "%s" in other files' % (taxa[i], fn, taxon_list[i]))
            
        if sequences is None:
            sequences = {}
            for t in taxon_list:
                sequences[t] = seq[t][:]
        else:
            for t in taxon_list:
                sequences[t] = sequences[t] + seq[t][:]
            
    longest = max([len(t) for t in taxon_list])
    taxonfmt = '  %%%ds' % longest
    f = open(outfn, 'w')
    f.write('#nexus\n\n')
    f.write('begin data;\n')
    f.write('  dimensions ntax=%d nchar=%d;\n' % (numtax, numchar))
    f.write('  format datatype=dna gap=-;\n')
    f.write('  matrix\n')
    for t in taxon_list:
        taxon_name = re.sub(r'\s+', '_', t)
        f.write(taxonfmt % taxon_name)
        f.write(' %s\n' % sequences[t])
    f.write('  ;\n')
    f.write('end;\n')
    f.close()

indata = {}
for fn in infilenames:
    indata[fn] = readNexusFile(fn)
writeNexusFile(outfilename, infilenames, indata)    
```
