## Supplementary material for "Estimating Bayesian Phylogenetic Information Content Using Geodesic Distances": seqgen.sh

```
#!/bin/bash

BASEDIR="$( cd .. && pwd )"
echo "$BASEDIR"

rm -rf $BASEDIR/reps
mkdir $BASEDIR/reps

cp $BASEDIR/../calcinfo.py $BASEDIR/reps

#python3 ~/bin/kappa2tratio.py --kappa=2.14183  --piA=0.236464  --piC=0.191802 --piG=0.388702
#python3 ~/bin/kappa2tratio.py --kappa=1.95213  --piA=0.301213  --piC=0.238401 --piG=0.207913
#python3 ~/bin/kappa2tratio.py --kappa=7.54411  --piA=0.300500  --piC=0.155705 --piG=0.123789

# kappa  = 2.14183
# piA    = 0.23646
# piC    = 0.19180
# piG    = 0.38870
# piT    = 0.18303
# tratio = 1.16097
# 
# kappa  = 1.95213
# piA    = 0.30121
# piC    = 0.23840
# piG    = 0.20791
# piT    = 0.25247
# tratio = 0.95933
# 
# kappa  = 7.54411
# piA    = 0.30050
# piC    = 0.15571
# piG    = 0.12379
# piT    = 0.42001
# tratio = 3.16862

# Set master random number seed
RANDOM=45971  # used for reps 1-20

# Let $N hold number of simulation replicates
N=20

# Generate seeds for generating each simulated data set
SEEDFIRST=()
SEEDSECOND=()
SEEDTHIRD=()
for ((i=0; i<$N; i++)) do
    SEEDFIRST+=($RANDOM)
    SEEDSECOND+=($RANDOM)
    SEEDTHIRD+=($RANDOM)
done

# Create a directory rep0 for the empirical analysis
mkdir -p $BASEDIR/reps/rep0/data
cp $BASEDIR/fig-saturation.Rev $BASEDIR/reps/rep0
cp $BASEDIR/data/rbcl.nex    $BASEDIR/reps/rep0/data
cp $BASEDIR/data/rbcl1st.nex $BASEDIR/reps/rep0/data
cp $BASEDIR/data/rbcl2nd.nex $BASEDIR/reps/rep0/data
cp $BASEDIR/data/rbcl3rd.nex $BASEDIR/reps/rep0/data

# Create a directory for each of the $N replicates and simulate data for it
for ((idx=0; idx<$N; idx++)) do
    repidx=$((idx + 1))
    echo "********** simulation replicate $repidx **********"
    seedfirst=${SEEDFIRST[$idx]}
    seedsecond=${SEEDSECOND[$idx]}
    seedthird=${SEEDTHIRD[$idx]}
    mkdir -p $BASEDIR/reps/rep$repidx/data
    cd $BASEDIR/reps/rep$repidx
    cp $BASEDIR/fig-saturation.Rev .
    cd data
    seq-gen -mHKY -l467 -n1 -p1 -f 0.236464 0.191802 0.388702 0.183033 -t1.16097 -z$seedfirst  -on -s0.3313 < $BASEDIR/iqtree/partition.nex.treefile > sim1st.nex
    seq-gen -mHKY -l467 -n1 -p1 -f 0.301213 0.238401 0.207913 0.252473 -t0.95933 -z$seedsecond -on -s0.1454 < $BASEDIR/iqtree/partition.nex.treefile > sim2nd.nex
    seq-gen -mHKY -l467 -n1 -p1 -f 0.300500 0.155705 0.123789 0.420006 -t3.16862 -z$seedthird  -on -s2.5233 < $BASEDIR/iqtree/partition.nex.treefile > sim3rd.nex
    python3 $BASEDIR/seqgen/concat.py --in sim1st.nex --in sim2nd.nex --in sim3rd.nex --out sim.nex
done
```
