## Supplementary material for "Estimating Bayesian Phylogenetic Information Content Using Geodesic Distances": slurm-template.sh

```
#!/bin/bash

#SBATCH --job-name=__JOBNAME__                          # Job name
#SBATCH --output=__JOBNAME__%a.out                      # stdout: %a becomes the array id
#SBATCH --error=__JOBNAME__%a.err                       # stderr: %a becomes the array id
#SBATCH --array=1-8                             # Two reps each of 1st, 2nd, 3rd, all sites
#SBATCH --partition=general                     # Name of Partition
#SBATCH --ntasks=1                              # Maximum CPU cores for each job
#SBATCH --nodes=1                               # Ensure all cores are from the same node for each job
#SBATCH --cpus-per-task=1                       # CPU-cores per task (>1 if multi-threaded tasks) for each job
#SBATCH --constraint='epyc128'                  # Target the AMD Epyc 128-core node architecture
#SBATCH --mem=20G                               # Request 20GB of available RAM
#SBATCH --mail-type=ALL                         # Event(s) that triggers email notification (BEGIN,END,FAIL,ALL)
#SBATCH --mail-user=        # Destination email address

# Specify paths to RevBayes executable
basedir=`pwd`
RB=$HOME/bin/rb131
OP=$HOME/bin/op194

locus="rbcl"

echo "Creating directories and running revbayes"

# Set epsilon for Frechet mean stopping criterion
FRECHET_EPSILON="0.00001"

# Set master random number seed
RANDOM=13579

# Load modules needed
module load python/3.12.5

# Generate seed for each analysis
post1stseed1=("$RANDOM")
post1stseed2=("$RANDOM")
post2ndseed1=("$RANDOM")
post2ndseed2=("$RANDOM")
post3rdseed1=("$RANDOM")
post3rdseed2=("$RANDOM")
post12seed1=("$RANDOM")
post12seed1=("$RANDOM")
post123seed1=("$RANDOM")
post123seed1=("$RANDOM")

prior1stseed1=("$RANDOM")
prior1stseed2=("$RANDOM")
prior2ndseed1=("$RANDOM")
prior2ndseed2=("$RANDOM")
prior3rdseed1=("$RANDOM")
prior3rdseed2=("$RANDOM")
prior12seed1=("$RANDOM")
prior12seed2=("$RANDOM")
prior123seed1=("$RANDOM")
prior123seed2=("$RANDOM")

data1st="\"$basedir/data/__DATAFIRST__.nex\""
data2nd="\"$basedir/data/__DATASECOND__.nex\""
data3rd="\"$basedir/data/__DATATHIRD__.nex\""
data123="\"$basedir/data/__DATAALL__.nex\""

echo
echo "********************************"
echo "***** Creating directories *****"
echo "********************************"
# Create the main directory
mkdir -p $basedir/$locus

if [[ $SLURM_ARRAY_TASK_ID -eq 1 ]]; then
    # Create the 1st1-codon-position sites posterior directory
    mkdir -p $basedir/$locus/1st1/posterior
    echo "datafile <- $data1st; rnseed <- $post1stseed1; run_under_prior <- FALSE" > $locus/1st1/posterior/adhoc.Rev
    
    # Create the 1st1-codon-position prior directory
    mkdir -p $basedir/$locus/1st1/prior
    echo "datafile <- $data1st; rnseed <- $prior1stseed1; run_under_prior <- TRUE" > $locus/1st1/prior/adhoc.Rev
elif [[ $SLURM_ARRAY_TASK_ID -eq 2 ]]; then
    # Create the 1st2-codon-position sites posterior directory
    mkdir -p $basedir/$locus/1st2/posterior
    echo "datafile <- $data1st; rnseed <- $post1stseed2; run_under_prior <- FALSE" > $locus/1st2/posterior/adhoc.Rev
    
    # Create the 1st-codon-position prior directory
    mkdir -p $basedir/$locus/1st2/prior
    echo "datafile <- $data1st; rnseed <- $prior1stseed2; run_under_prior <- TRUE" > $locus/1st2/prior/adhoc.Rev
elif [[ $SLURM_ARRAY_TASK_ID -eq 3 ]]; then
    # Create the 2nd1-codon-position sites posterior directory
    mkdir -p $basedir/$locus/2nd1/posterior
    echo "datafile <- $data2nd; rnseed <- $post2ndseed1; run_under_prior <- FALSE" > $locus/2nd1/posterior/adhoc.Rev
    
    # Create the 2nd1-codon-position prior directory
    mkdir -p $basedir/$locus/2nd1/prior
    echo "datafile <- $data2nd; rnseed <- $prior2ndseed1; run_under_prior <- TRUE" > $locus/2nd1/prior/adhoc.Rev
elif [[ $SLURM_ARRAY_TASK_ID -eq 4 ]]; then
    # Create the 2nd2-codon-position sites posterior directory
    mkdir -p $locus/2nd2/posterior
    echo "datafile <- $data2nd; rnseed <- $post2ndseed2; run_under_prior <- FALSE" > $locus/2nd2/posterior/adhoc.Rev
    
    # Create the 2nd2-codon-position prior directory
    mkdir -p $basedir/$locus/2nd2/prior
    echo "datafile <- $data2nd; rnseed <- $prior2ndseed2; run_under_prior <- TRUE" > $locus/2nd2/prior/adhoc.Rev
elif [[ $SLURM_ARRAY_TASK_ID -eq 5 ]]; then
    # Create the 3rd1-codon-position sites posterior directory
    mkdir -p $basedir/$locus/3rd1/posterior
    echo "datafile <- $data3rd; rnseed <- $post3rdseed1; run_under_prior <- FALSE" > $locus/3rd1/posterior/adhoc.Rev
    
    # Create the 3rd1-codon-position prior directory
    mkdir -p $basedir/$locus/3rd1/prior
    echo "datafile <- $data3rd; rnseed <- $prior3rdseed1; run_under_prior <- TRUE" > $locus/3rd1/prior/adhoc.Rev
elif [[ $SLURM_ARRAY_TASK_ID -eq 6 ]]; then
    # Create the 3rd2-codon-position sites posterior directory
    mkdir -p $basedir/$locus/3rd2/posterior
    echo "datafile <- $data3rd; rnseed <- $post3rdseed2; run_under_prior <- FALSE" > $locus/3rd2/posterior/adhoc.Rev
    
    # Create the 3rd2-codon-position prior directory
    mkdir -p $basedir/$locus/3rd2/prior
    echo "datafile <- $data3rd; rnseed <- $prior3rdseed2; run_under_prior <- TRUE" > $locus/3rd2/prior/adhoc.Rev
elif [[ $SLURM_ARRAY_TASK_ID -eq 7 ]]; then
    # Create the all sites posterior directory
    mkdir -p $basedir/$locus/all1/posterior
    echo "datafile <- $data123; rnseed <- $post123seed1; run_under_prior <- FALSE" > $locus/all1/posterior/adhoc.Rev
    
    # Create the all prior directory
    mkdir -p $basedir/$locus/all1/prior
    echo "datafile <- $data123; rnseed <- $prior123seed1; run_under_prior <- TRUE" > $locus/all1/prior/adhoc.Rev
else
    # Create the all sites posterior directory
    mkdir -p $locus/all2/posterior
    echo "datafile <- $data123; rnseed <- $post123seed2; run_under_prior <- FALSE" > $locus/all2/posterior/adhoc.Rev
    
    # Create the all prior directory
    mkdir -p $basedir/$locus/all2/prior
    echo "datafile <- $data123; rnseed <- $prior123seed2; run_under_prior <- TRUE" > $locus/all2/prior/adhoc.Rev
fi

if [[ $SLURM_ARRAY_TASK_ID -eq 1 ]]; then
    echo
    echo "*************************************************************"
    echo "***** Running RevBayes (1st-codon-position posterior 1) *****"
    echo "*************************************************************"
    # Sample the posterior and compute the prior mean tree
    cd $basedir/$locus/1st1/posterior 
    $RB $basedir/fig-saturation.Rev
    
    echo
    echo "*********************************************************"
    echo "***** Running RevBayes (1st-codon-position prior 1) *****"
    echo "*********************************************************"
    # Sample the prior and compute the prior mean tree
    cd $basedir/$locus/1st1/prior
    $RB $basedir/fig-saturation.Rev

    echo
    echo "***********************************"
    echo "***** Calculating information *****"
    echo "***********************************"
    cd $basedir/$locus/1st1
    rm -rf results
    python3 $basedir/../calcinfo.py --oppath $OP --dirname results --prefix info --skip 1 \
        --postfile posterior/output/landplant.trees --priorfile prior/output/landplant.trees 
    python3 $basedir/../calcinfo.py --oppath $OP --hpdrad --radpct 95 --dirname results --prefix info95 --skip 1 \
        --postfile posterior/output/landplant.trees --priorfile prior/output/landplant.trees 
elif [[ $SLURM_ARRAY_TASK_ID -eq 2 ]]; then
    echo
    echo "*************************************************************"
    echo "***** Running RevBayes (1st-codon-position posterior 2) *****"
    echo "*************************************************************"
    # Sample the posterior and compute the prior mean tree
    cd $basedir/$locus/1st2/posterior 
    $RB $basedir/fig-saturation.Rev
    
    echo
    echo "********************************************************"
    echo "***** Running RevBayes (1st-codon-position prior 2) *****"
    echo "********************************************************"
    # Sample the prior and compute the prior mean tree
    cd $basedir/$locus/1st2/prior
    $RB $basedir/fig-saturation.Rev

    echo
    echo "***********************************"
    echo "***** Calculating information *****"
    echo "***********************************"
    cd $basedir/$locus/1st2
    rm -rf results
    python3 $basedir/../calcinfo.py --oppath $OP --dirname results --prefix info --skip 1 \
        --postfile posterior/output/landplant.trees --priorfile prior/output/landplant.trees 
    python3 $basedir/../calcinfo.py --oppath $OP --hpdrad --radpct 95 --dirname results --prefix info95 --skip 1 \
        --postfile posterior/output/landplant.trees --priorfile prior/output/landplant.trees 
elif [[ $SLURM_ARRAY_TASK_ID -eq 3 ]]; then
    echo
    echo "*************************************************************"
    echo "***** Running RevBayes (2nd-codon-position posterior 1) *****"
    echo "*************************************************************"
    # Sample the posterior and compute the prior mean tree
    cd $basedir/$locus/2nd1/posterior 
    $RB $basedir/fig-saturation.Rev
    
    echo
    echo "*********************************************************"
    echo "***** Running RevBayes (2nd-codon-position prior 1) *****"
    echo "*********************************************************"
    # Sample the prior and compute the prior mean tree
    cd $basedir/$locus/2nd1/prior
    $RB $basedir/fig-saturation.Rev

    echo
    echo "***********************************"
    echo "***** Calculating information *****"
    echo "***********************************"
    cd $basedir/$locus/2nd1
    rm -rf results
    python3 $basedir/../calcinfo.py --oppath $OP --dirname results --prefix info --skip 1 \
        --postfile posterior/output/landplant.trees --priorfile prior/output/landplant.trees 
    python3 $basedir/../calcinfo.py --oppath $OP --hpdrad --radpct 95 --dirname results --prefix info95 --skip 1 \
        --postfile posterior/output/landplant.trees --priorfile prior/output/landplant.trees 
elif [[ $SLURM_ARRAY_TASK_ID -eq 4 ]]; then
    echo
    echo "*************************************************************"
    echo "***** Running RevBayes (2nd-codon-position posterior 2) *****"
    echo "*************************************************************"
    # Sample the posterior and compute the prior mean tree
    cd $basedir/$locus/2nd2/posterior 
    $RB $basedir/fig-saturation.Rev
    
    echo
    echo "*********************************************************"
    echo "***** Running RevBayes (2nd-codon-position prior 2) *****"
    echo "*********************************************************"
    # Sample the prior and compute the prior mean tree
    cd $basedir/$locus/2nd2/prior
    $RB $basedir/fig-saturation.Rev

    echo
    echo "***********************************"
    echo "***** Calculating information *****"
    echo "***********************************"
    cd $basedir/$locus/2nd2
    rm -rf results
    python3 $basedir/../calcinfo.py --oppath $OP --dirname results --prefix info --skip 1 \
        --postfile posterior/output/landplant.trees --priorfile prior/output/landplant.trees 
    python3 $basedir/../calcinfo.py --oppath $OP --hpdrad --radpct 95 --dirname results --prefix info95 --skip 1 \
        --postfile posterior/output/landplant.trees --priorfile prior/output/landplant.trees 
elif [[ $SLURM_ARRAY_TASK_ID -eq 5 ]]; then
    echo
    echo "*************************************************************"
    echo "***** Running RevBayes (3rd-codon-position posterior 1) *****"
    echo "*************************************************************"
    # Sample the posterior and compute the prior mean tree
    cd $basedir/$locus/3rd1/posterior
    $RB $basedir/fig-saturation.Rev
    
    echo
    echo "*********************************************************"
    echo "***** Running RevBayes (3rd-codon-position prior 1) *****"
    echo "*********************************************************"
    # Sample the prior and compute the prior mean tree
    cd $basedir/$locus/3rd1/prior
    $RB $basedir/fig-saturation.Rev

    echo
    echo "***********************************"
    echo "***** Calculating information *****"
    echo "***********************************"
    cd $basedir/$locus/3rd1
    rm -rf results
    python3 $basedir/../calcinfo.py --oppath $OP --dirname results --prefix info --skip 1 \
        --postfile posterior/output/landplant.trees --priorfile prior/output/landplant.trees 
    python3 $basedir/../calcinfo.py --oppath $OP --hpdrad --radpct 95 --dirname results --prefix info95 --skip 1 \
        --postfile posterior/output/landplant.trees --priorfile prior/output/landplant.trees 
elif [[ $SLURM_ARRAY_TASK_ID -eq 6 ]]; then
    echo
    echo "*************************************************************"
    echo "***** Running RevBayes (3rd-codon-position posterior 2) *****"
    echo "*************************************************************"
    # Sample the posterior and compute the prior mean tree
    cd $basedir/$locus/3rd2/posterior
    $RB $basedir/fig-saturation.Rev
    
    echo
    echo "*********************************************************"
    echo "***** Running RevBayes (3rd-codon-position prior 2) *****"
    echo "*********************************************************"
    # Sample the prior and compute the prior mean tree
    cd $basedir/$locus/3rd2/prior
    $RB $basedir/fig-saturation.Rev

    echo
    echo "***********************************"
    echo "***** Calculating information *****"
    echo "***********************************"
    cd $basedir/$locus/3rd2
    rm -rf results
    python3 $basedir/../calcinfo.py --oppath $OP --dirname results --prefix info --skip 1 \
        --postfile posterior/output/landplant.trees --priorfile prior/output/landplant.trees 
    python3 $basedir/../calcinfo.py --oppath $OP --hpdrad --radpct 95 --dirname results --prefix info95 --skip 1 \
        --postfile posterior/output/landplant.trees --priorfile prior/output/landplant.trees 
elif [[ $SLURM_ARRAY_TASK_ID -eq 7 ]]; then
    echo
    echo "****************************************************"
    echo "***** Running RevBayes (all-sites posterior 1) *****"
    echo "****************************************************"
    # Sample the posterior and compute the prior mean tree
    cd $basedir/$locus/all1/posterior
    $RB $basedir/fig-saturation.Rev
    
    echo
    echo "************************************************"
    echo "***** Running RevBayes (all-sites prior 1) *****"
    echo "************************************************"
    # Sample the prior and compute the prior mean tree
    cd $basedir/$locus/all1/prior
    $RB $basedir/fig-saturation.Rev

    echo
    echo "***********************************"
    echo "***** Calculating information *****"
    echo "***********************************"
    cd $basedir/$locus/all1
    rm -rf results
    python3 $basedir/../calcinfo.py --oppath $OP --dirname results --prefix info --skip 1 \
        --postfile posterior/output/landplant.trees --priorfile prior/output/landplant.trees 
    python3 $basedir/../calcinfo.py --oppath $OP --hpdrad --radpct 95 --dirname results --prefix info95 --skip 1 \
        --postfile posterior/output/landplant.trees --priorfile prior/output/landplant.trees 
else
    echo
    echo "****************************************************"
    echo "***** Running RevBayes (all-sites posterior 2) *****"
    echo "****************************************************"
    # Sample the posterior and compute the prior mean tree
    cd $basedir/$locus/all2/posterior
    $RB $basedir/fig-saturation.Rev
    
    echo
    echo "************************************************"
    echo "***** Running RevBayes (all-sites prior 2) *****"
    echo "************************************************"
    # Sample the prior and compute the prior mean tree
    cd $basedir/$locus/all2/prior
    $RB $basedir/fig-saturation.Rev

    echo
    echo "***********************************"
    echo "***** Calculating information *****"
    echo "***********************************"
    cd $basedir/$locus/all2
    rm -rf results
    python3 $basedir/../calcinfo.py --oppath $OP --dirname results --prefix info --skip 1 \
        --postfile posterior/output/landplant.trees --priorfile prior/output/landplant.trees 
    python3 $basedir/../calcinfo.py --oppath $OP --hpdrad --radpct 95 --dirname results --prefix info95 --skip 1 \
        --postfile posterior/output/landplant.trees --priorfile prior/output/landplant.trees 
fi
```
