## Supplementary material for "Estimating Bayesian Phylogenetic Information Content Using Geodesic Distances": summarize.py

```
import argparse,sys,os,subprocess,re,math,glob

locus_name = 'rbcl'

parser = argparse.ArgumentParser(
    formatter_class=argparse.RawDescriptionHelpFormatter,
    description='''Tabulates information for the saturation example.''',
)
parser.add_argument('--repsdir', default='reps', help='directory containing replicate directories rep0...rep10')
parser.add_argument('--hpdrad', default=False, action='store_true', help='use mean tree from HPD credible set of trees')
parser.add_argument('--oppath', default='op194', help='path to the OP program')
settings = vars(parser.parse_args())

repsdir = settings['repsdir']
hpdrad = settings['hpdrad']
op_path = settings['oppath']

repdirs = glob.glob(os.path.join(repsdir, 'rep*'))
nsimreps = len(repdirs) - 1

scriptdir = os.path.dirname(os.path.realpath(sys.argv[0]))
outf = None

def output(s = '', line_ending='\n'):
    global outf
    if line_ending is None:
        print(s, end='')
        outf.write('%s' % s)
    else:
        print(s, end=line_ending)
        outf.write('%s%s' % (s, line_ending))
        
def averageInfo(info1, info2):
    info = {}
    for s in info1.keys():
        info[s] = 0.5*(info1[s] + info2[s])
    return info

def extractInfo(fn):
    if not os.path.exists(fn):
        print('Could not exxtractInfo from "%s" because it does not exist' % fn)
        return {
            'LCRrad':None,
            'Ipct':None,
            'post_samplesize':None,
            'prior_samplesize':None,
            'post_treelength':None,
            'prior_treelength':None,
            'post_variance':None,
            'prior_variance':None,
            'post_scaler':None,
            'prior_scaler':None,
            'post_radius':None,
            'prior_radius':None
        }
        
    stuff = open(fn, 'r').read()

    m = re.search(r'([.0-9]+) = posterior sample size', stuff, re.M | re.S)
    assert m is not None, 'Could not extract post_samplesize from file "%s"' % fn
    post_samplesize = float(m.group(1))

    m = re.search(r'([.0-9]+) = prior sample size', stuff, re.M | re.S)
    assert m is not None, 'Could not extract prior_samplesize from file "%s"' % fn
    prior_samplesize = float(m.group(1))

    m = re.search(r'([.0-9]+) = posterior variance', stuff, re.M | re.S)
    assert m is not None, 'Could not extract post_variance from file "%s"' % fn
    post_variance = float(m.group(1))

    m = re.search(r'([.0-9]+) = prior variance', stuff, re.M | re.S)
    assert m is not None, 'Could not extract prior_variance from file "%s"' % fn
    prior_variance = float(m.group(1))

    m = re.search(r'([.0-9]+) = posterior mean tree length', stuff, re.M | re.S)
    assert m is not None, 'Could not extract post_treelength from file "%s"' % fn
    post_treelength = float(m.group(1))

    m = re.search(r'([.0-9]+) = prior mean tree length', stuff, re.M | re.S)
    assert m is not None, 'Could not extract prior_treelength from file "%s"' % fn
    prior_treelength = float(m.group(1))

    m = re.search(r'([.0-9]+) = LCRrad', stuff, re.M | re.S)
    assert m is not None, 'Could not extract LCRrad from file "%s"' % fn
    LCRrad = float(m.group(1))

    m = re.search(r'([.0-9]+) = information as percentage', stuff, re.M | re.S)
    assert m is not None, 'Could not extract Ipct from file "%s"' % fn
    Ipct = float(m.group(1))

    m = re.search(r'([.0-9]+) = posterior scaler', stuff, re.M | re.S)
    assert m is not None, 'Could not extract post_scaler from file "%s"' % fn
    post_scaler = float(m.group(1))

    m = re.search(r'([.0-9]+) = prior scaler', stuff, re.M | re.S)
    assert m is not None, 'Could not extract prior_scaler from file "%s"' % fn
    prior_scaler = float(m.group(1))

    m = re.search(r'([.0-9]+) = posterior 95% radius', stuff, re.M | re.S)
    assert m is not None, 'Could not extract post_radius from file "%s"' % fn
    post_radius = float(m.group(1))

    m = re.search(r'([.0-9]+) = prior 95% radius', stuff, re.M | re.S)
    assert m is not None, 'Could not extract prior_radius from file "%s"' % fn
    prior_radius = float(m.group(1))

    return {
        'LCRrad':LCRrad,
        'Ipct':Ipct,
        'post_samplesize':post_samplesize,
        'prior_samplesize':prior_samplesize,
        'post_treelength':post_treelength,
        'prior_treelength':prior_treelength,
        'post_variance':post_variance,
        'prior_variance':prior_variance,
        'post_scaler':post_scaler,
        'prior_scaler':prior_scaler,
        'post_radius':post_radius,
        'prior_radius':prior_radius
    }
    
def extractMeanDiffFrom(fn):
    stuff = open(fn, 'r').read()
    m = re.search(r'bhvdist = ([-.0-9e]+)', stuff, re.S | re.M)
    assert m is not None, 'Failed to extract bhvdist from file "%s"' % fn
    return float(m.group(1))

def calcDissForPair(s1, s2, is_hpd, mode):
    # mode is one of the following:
    #   radius: use squared radius as measure of dispersion
    #   stddev: use squared standard deviation as measure of dispersion
    infodict = info
    if is_hpd:
        infodict = info95
        
    # Calculate pooled standard deviation
    n1        = infodict[s1]['post_samplesize']
    scaler1   = infodict[s1]['post_scaler']
    variance1 = infodict[s1]['post_variance']
    radius1   = infodict[s1]['post_radius']
    stddev1   = math.sqrt(variance1) 
    sd1       = scaler1*stddev1
    rad1      = scaler1*radius1
    
    n2        = infodict[s2]['post_samplesize']
    scaler2   = infodict[s2]['post_scaler']
    variance2 = infodict[s2]['post_variance']
    radius2   = infodict[s2]['post_radius']
    stddev2   = math.sqrt(variance2) 
    sd2       = scaler2*stddev2
    rad2      = scaler2*radius2
    
    if mode == 'stddev':
        var1 = math.pow(sd1, 2)
        var2 = math.pow(sd2, 2)
        pooled_numer = var1*(n1 - 1) + var2*(n2 - 1)
        pooled_denom = n1 + n2 - 2
        pooled_sd = math.sqrt(pooled_numer/pooled_denom)
    else:
        rad1squared = math.pow(rad1, 2)
        rad2squared = math.pow(rad2, 2)
        pooled_numer = rad1squared*(n1 - 1) + rad2squared*(n2 - 1)
        pooled_denom = n1 + n2 - 2
        pooled_sd = math.sqrt(pooled_numer/pooled_denom)
    
    cmds = [op_path,
        '--reftree', os.path.join(scriptdir, locus_name, s1, 'results', is_hpd and 'info95' or 'info', 'op-post.tre'),
        '--refskip', '0',
        '--refscale', '%g' % scaler1,
        '--treefile', os.path.join(scriptdir, locus_name, s2, 'results', is_hpd and 'info95' or 'info', 'op-post.tre'),       
        '--skip', '0',
        '--scale', '%g' % scaler2,
        '--prefix', os.path.join(scriptdir, 'results', 'meandiff', 'meandiff-%s-%s-%s-%s' % (is_hpd and 'hpd' or 'rad', mode, s1, s2))
    ]
    completed_process = subprocess.run(cmds, cwd=scriptdir, capture_output=True)
    try:
        completed_process.check_returncode()
    except sub.CalledProcessError:
        print('\nAn error occurred while running OP for "%s"' % simdir)
        print('Standard error:')
        print(completed_process.stderr)
        print('Standard output:')
        print(completed_process.stdout)
        sys.exit('Aborted.')
    
    mean_diff = extractMeanDiffFrom(os.path.join(scriptdir, 'results', 'meandiff', 'meandiff-%s-%s-%s-%s.tre' % (is_hpd and 'hpd' or 'rad', mode, s1, s2)))
    #effect1 = mean_diff/sd1
    #effect2 = mean_diff/sd2
    cohens_d = mean_diff/pooled_sd
        
    return cohens_d
    
def distToTrueTree(dirname, which):
    global hpdrad, true_scaler
    
    # Find tree length and calculate scaler for "which" mean tree
    fn = None
    if hpdrad:
        fn = os.path.join(dirname, which, 'results','info95-output.txt')
    else:
        fn = os.path.join(dirname, which, 'results','info-output.txt')
    info = extractInfo(fn)
    TL = info['post_treelength']
    scaler = 1.0/TL
    
    # Find distance between the mean tree for "which" and the true tree
    which_prefix = os.path.join(results_dir, '%s-true' % which)
    op_cmds = [op_path, 
        '--reftree', os.path.join('iqtree', 'truetree.tre'), 
        '--refskip','0',
        '--refscale', '%.9f' % true_scaler,
        '--treefile', os.path.join(dirname, which, 'results', hpdrad and 'info95' or 'info','op-post.tre'),
        '--skip', '0',
        '--scale', '%.9f' % scaler,
        '--prefix', '%s' % which_prefix
    ]
    completed_process = subprocess.run(op_cmds, cwd=scriptdir, capture_output=True)
    try:
        completed_process.check_returncode()
    except subprocess.CalledProcessError:
        print('\nAn error occurred while running OP to find distance of %s mean tree to the true tree' % which)
        print('\nStandard error:')
        print(completed_process.stderr.decode('utf-8'))
        print('\nStandard output:')
        print(completed_process.stdout.decode('utf-8'))
        print('\nCommands:')
        print(' '.join(op_cmds))
        sys.exit('Aborted.')
    mean_diff = extractMeanDiffFrom('%s.tre' % which_prefix)
    return (mean_diff, scaler)
    
def calcDistToTrueTree(dirname):
    # dirname is e.g. 'reps/rep3/<locus_name>'
    truedist = {}
    scaler = {}
    
    dist11, scaler11 = distToTrueTree(dirname, '1st1')
    truedist['1st1'] = dist11
    scaler['1st1'] = scaler11
    
    dist12, scaler12 = distToTrueTree(dirname, '1st2')
    truedist['1st2'] = dist12
    scaler['1st2'] = scaler12
    
    dist21, scaler21 = distToTrueTree(dirname, '2nd1')
    truedist['2nd1'] = dist21
    scaler['2nd1'] = scaler21
    
    dist22, scaler22 = distToTrueTree(dirname, '2nd2')
    truedist['2nd2'] = dist22
    scaler['2nd2'] = scaler22
    
    dist31, scaler31 = distToTrueTree(dirname, '3rd1')
    truedist['3rd1'] = dist31
    scaler['3rd1'] = scaler31
    
    dist32, scaler32 = distToTrueTree(dirname, '3rd2')
    truedist['3rd2'] = dist32
    scaler['3rd2'] = scaler32
    
    dist1, scaler1 = distToTrueTree(dirname, 'all1')
    truedist['all1'] = dist1
    scaler['all1'] = scaler1

    dist2, scaler2 = distToTrueTree(dirname, 'all2')
    truedist['all2'] = dist2
    scaler['all2'] = scaler2

    return truedist, scaler

if __name__ == '__main__':
    results_dir = os.path.join(scriptdir, 'results')
    if os.path.exists(results_dir):
        sys.exit('Please move or rename the directory "%s"' % results_dir)
    os.mkdir(results_dir)
        
    # Open an output file 
    outf = open('output.txt', 'w')

    if hpdrad:
        output('Results based on 95% highest posterior sample.')
    else:
        output('Results below based on 95% samples closest to mean tree.')
        
    output('Number of simulation replicates: %d' % nsimreps)
    
    # Save the true tree (estimated by IQ-TREE from real data) to a nexus file
    true_tree = open(os.path.join('iqtree', 'partition.nex.treefile'), 'r').read()
    truef = open(os.path.join('iqtree', 'truetree.tre'), 'w')
    truef.write('#nexus\n\n')
    truef.write('begin trees;\n')
    truef.write('  tree truth = %s' % true_tree)
    truef.write('end;\n')
    truef.close()  
    
    # Get length of true_tree
    edgelens = re.findall(r':([-.e0-9]+)(?:[,)])', true_tree)
    nedgelens = len(edgelens)
    print('Number of edges in true tree: %d' % nedgelens)
    trueTL = 0.0
    for edgelen in edgelens:
        trueTL += float(edgelen)
    true_scaler = 1.0/trueTL
    output('True tree length: %12.5f' % trueTL)
    output('True tree scaler: %12.5f' % true_scaler)
    
    # Calculate distance to the true tree for every subset in each replicate
    truedistscaler = []
    for i in range(nsimreps):  
        rep = i + 1      
        dirname = os.path.join(repsdir, 'rep%d' % rep, locus_name)
        # truedict['1st1'] holds distance to true tree for subset 1st1
        # scalerdict['1st1'] holds scaler (1/TL) for mean tree of subset 1st1
        truedict, scalerdict = calcDistToTrueTree(dirname)
        truedistscaler.append((truedict, scalerdict))
    
    ####################################################################
    # First table shows distances to true tree for each rep and subset #
    ####################################################################
    output('\nGeodesic distances of mean tree to true tree:\n')
    subsets = ['1st1', '1st2', '2nd1', '2nd2', '3rd1', '3rd2', 'all1', 'all2']
    nsubsets = len(subsets)
    output('%12s %12s %12s %12s %12s %12s %12s %12s %12s' % ('replicate', '1st1', '1st2', '2nd1', '2nd2', '3rd1', '3rd2', 'all1', 'all2'))
    output('%12s %12s %12s %12s %12s %12s %12s %12s %12s' % ('-----------', '-----------', '-----------', '-----------', '-----------', '-----------', '-----------', '-----------', '-----------'))
    sums = [0.0]*nsubsets
    sumsqs = [0.0]*nsubsets
    for i in range(nsimreps):
        rep = i + 1
        output('%12d' % rep, None)
        truedict, scalerdict = truedistscaler[i]
        for j,which in enumerate(subsets):
            d = truedict[which]
            output(' %12.5f' % d, None)
            sums[j] += d
            sumsqs[j] += d*d
        output()
    output('%12s %12s %12s %12s %12s %12s %12s %12s %12s' % ('-----------', '-----------', '-----------', '-----------', '-----------', '-----------', '-----------', '-----------', '-----------'))
    means = [0.0]*nsubsets
    stdevs = [0.0]*nsubsets
    for i in range(nsubsets):
        mean = sums[i]/nsimreps
        var = (sumsqs[i] - mean*mean*nsimreps)/(nsimreps-1)
        sd = math.sqrt(var)    
        means[i] = mean
        stdevs[i] = sd
    output('%12s' % 'mean', None)
    for i in range(nsubsets):
        output(' %12.5f' % means[i], None)
    output()
    output('%12s' % 'sd', None)
    for i in range(nsubsets):
        output(' %12.5f' % stdevs[i], None)
    output()
    
    # Calculate information (LCR and Ipct) for each subset in each rep
    # Include rep0, which holds the results for the empirical data
    n = nsimreps + 1
    repinfo = []
    for rep in range(n):
        dirname = os.path.join(repsdir, 'rep%d' % rep, locus_name)
        info = {}
        for s in subsets:
            if hpdrad:
                info[s] = extractInfo(os.path.join(dirname, s, 'results', 'info95-output.txt'))
            else:
                info[s] = extractInfo(os.path.join(dirname, s, 'results', 'info-output.txt'))
        repinfo.append(info)
        
    ##################################################
    # Second table shows LCR for each rep and subset #
    ##################################################
    output('\nLCR for each replicate and subset:\n')
    output('%12s %12s %12s %12s %12s %12s %12s %12s %12s' % ('replicate', '1st1', '1st2', '2nd1', '2nd2', '3rd1', '3rd2', 'all1', 'all2'))
    output('%12s %12s %12s %12s %12s %12s %12s %12s %12s' % ('-----------', '-----------', '-----------', '-----------', '-----------', '-----------', '-----------', '-----------', '-----------'))

    sums = [0.0]*nsubsets
    sumsqs = [0.0]*nsubsets
    for i in range(nsimreps):
        rep = i + 1
        output('%12d' % rep, None)
        info = repinfo[rep]
        for j,which in enumerate(subsets):
            lcr = info[which]['LCRrad']
            output(' %12.5f' % lcr, None)
            sums[j] += lcr
            sumsqs[j] += lcr*lcr
        output()

    output('%12s %12s %12s %12s %12s %12s %12s %12s %12s' % ('-----------', '-----------', '-----------', '-----------', '-----------', '-----------', '-----------', '-----------', '-----------'))

    means = [0.0]*nsubsets
    stdevs = [0.0]*nsubsets
    for i in range(nsubsets):
        mean = sums[i]/nsimreps
        var = (sumsqs[i] - mean*mean*nsimreps)/(nsimreps-1)
        sd = math.sqrt(var)    
        means[i] = mean
        stdevs[i] = sd

    # Output line for rep0 (empirical data)
    output('%12s' % 'empirical', None)
    info = repinfo[0]
    for j,which in enumerate(subsets):
        lcr = info[which]['LCRrad']
        output(' %12.5f' % lcr, None)
    output()
        
    output('%12s' % 'mean', None)
    for i in range(nsubsets):
        output(' %12.5f' % means[i], None)
    output()

    output('%12s' % 'sd', None)
    for i in range(nsubsets):
        output(' %12.5f' % stdevs[i], None)
    output()

    ##################################################
    # Third table shows Ipct for each rep and subset #
    ##################################################
    output('\nIpct for each replicate and subset:\n')
    output('%12s %12s %12s %12s %12s %12s %12s %12s %12s' % ('replicate', '1st1', '1st2', '2nd1', '2nd2', '3rd1', '3rd2', 'all1', 'all2'))
    output('%12s %12s %12s %12s %12s %12s %12s %12s %12s' % ('-----------', '-----------', '-----------', '-----------', '-----------', '-----------', '-----------', '-----------', '-----------'))

    sums = [0.0]*nsubsets
    sumsqs = [0.0]*nsubsets
    for i in range(nsimreps):
        rep = i + 1
        output('%12d' % rep, None)
        info = repinfo[rep]
        for j,which in enumerate(subsets):
            ipct = info[which]['Ipct']
            output(' %12.5f' % ipct, None)
            sums[j] += ipct
            sumsqs[j] += ipct*ipct
        output()

    output('%12s %12s %12s %12s %12s %12s %12s %12s %12s' % ('-----------', '-----------', '-----------', '-----------', '-----------', '-----------', '-----------', '-----------', '-----------'))

    means = [0.0]*nsubsets
    stdevs = [0.0]*nsubsets
    for i in range(nsubsets):
        mean = sums[i]/nsimreps
        var = (sumsqs[i] - mean*mean*nsimreps)/(nsimreps-1)
        sd = math.sqrt(var)    
        means[i] = mean
        stdevs[i] = sd

    # Output line for rep0 (empirical data)
    output('%12s' % 'empirical', None)
    info = repinfo[0]
    for j,which in enumerate(subsets):
        ipct = info[which]['Ipct']
        output(' %12.5f' % ipct, None)
    output()
        
    output('%12s' % 'mean', None)
    for i in range(nsubsets):
        output(' %12.5f' % means[i], None)
    output()

    output('%12s' % 'sd', None)
    for i in range(nsubsets):
        output(' %12.5f' % stdevs[i], None)
    output()
```
