## Supplementary material for "Estimating Bayesian Phylogenetic Information Content Using Geodesic Distances": _readme.txt

To create fig-bloodroot:

Directory layout showing only directories and files mentioned in the instructions below:

     supplementary-materials
         calcinfo.py
         mds.py
         5-fig-bloodroot
             data
                 rps11-3p.nex
                 rps11-5p.nex
                 rps11.nex
*            results            
*               mds
*                   3a3b-hpd
*                   3a3b5a5b-hpd
*                   5a5b-hpd
*               mean-diffs
*               rps11
*                   3prime1
*                   3prime2
*                   5prime1
*                   5prime2
*                   all1
*                   all2
*               pairwise-dissonance.txt
             _readme.txt
             fig-bloodroot.Rev
             rps11-slurm.sh
             runmds.sh
             summarize.py

* file/directory created as a result of these instructions

1. Edit the rps11-slurm.sh and adapt the SBATCH commands at the top 
    to your cluster. Be sure the path to the RevBayes and OP
    executables are correct.

    If you do not have these programs installed, here is where you can get them:
        OP:         https://github.com/plewis/OP
        RevBayes:   https://revbayes.github.io/download
    
2. Specify "hpc" as an alias to your remote cluster in  ~/.ssh/config and
   create a "work/diss" directory inside your home directory on your cluster
   
   ssh hpc
   mkdir -p work/bloodroot/data
    
3. Upload files to your cluster:

    cd supplementary-materials
    scp calcinfo.py           hpc:work

    cd 5-fig-bloodroot
    scp fig-bloodroot.Rev     hpc:work/bloodroot
    scp rps11-slurm.sh        hpc:work/bloodroot

    cd data
    scp rps11-3p.nex          hpc:work/bloodroot/data
    scp rps11-5p.nex          hpc:work/bloodroot/data
    scp rps11.nex             hpc:work/bloodroot/data
    
4. Run the analyses on the cluster:

    ssh hpc
    cd work/bloodroot
    sbatch rps11-slurm.sh
    
    This will generate an rps11 directory containing subdirectories 3prime1,
    3prime2, 5prime1, 5prime2, all1, and all2.
    
5. Bring results back to your local computer:

    cd supplementary-materials/5-fig-bloodroot
    mkdir results
    cd results
    rsync -azPv hpc:work/bloodroot/rps11 .

6. Summarize results:

    cd supplementary-materials/5-fig-bloodroot
    python3 summarize.py --dir results
    
    This will generate a results/mean-diffs directory and 
    a results/pairwise-dissonance.txt file.
    
7. Execute the runmds.sh bash script to generate multidimensional scaling plots
    of the two 3' posterior samples, the two 5' posterior samples, and all four 
    combined. 
    
    cd supplementary-materials/5-fig-bloodroot
    chmod +x runmds.sh
    ./runmds.sh
    
    This will generate the results/mds directory containing subdirectories
    a3b-hpd, a3b5a5b-hpd, and a5b-hpd. Inside each of these subdirectories is a 
    file mds-plot.png containing the mds plot in PNG format.
    
