## Supplementary material for "Estimating Bayesian Phylogenetic Information Content Using Geodesic Distances": rps11-slurm.sh

```
#!/bin/bash
#SBATCH --job-name=rps11                        # job name
#SBATCH --array=1-6                             # 3prime1, 3prime2, 5prime1, 5prime2, all1, all2 (each analysis replicated once)
#SBATCH --partition=general                     # Name of Partition
#SBATCH --ntasks=1                              # Maximum CPU cores for each job
#SBATCH --nodes=1                               # Ensure all cores are from the same node for each job
#SBATCH --cpus-per-task=1                       # cpu-cores per task (>1 if multi-threaded tasks) for each job
#SBATCH --constraint='epyc128'                  # Target the AMD Epyc 128-core node architecture
#SBATCH --mem=20G                               # Request 20GB of available RAM
#SBATCH --mail-type=END                         # Event(s) that triggers email notification (BEGIN,END,FAIL,ALL)
#SBATCH --mail-user=        # Destination email address

echo "Creating directories and running revbayes"

# Specify paths to programs used
basedir=`pwd`
RB=$HOME/bin/rb131
OP=$HOME/bin/op194

# Set epsilon for Frechet mean stopping criterion
FRECHET_EPSILON="0.00001"

# Set master random number seed
RANDOM=45971

# Load modules needed
module load python/3.12.5
#module load gcc/15.1.0

# Generate seed for each analysis
post3prime1seed=("$RANDOM")
post3prime2seed=("$RANDOM")
post5prime1seed=("$RANDOM")
post5prime2seed=("$RANDOM")
postall1seed=("$RANDOM")
postall2seed=("$RANDOM")

prior3prime1seed=("$RANDOM")
prior3prime2seed=("$RANDOM")
prior5prime1seed=("$RANDOM")
prior5prime2seed=("$RANDOM")
priorall1seed=("$RANDOM")
priorall2seed=("$RANDOM")

data3prime="\"$basedir/data/rps11-3p.nex\""
data5prime="\"$basedir/data/rps11-5p.nex\""
dataall="\"$basedir/data/rps11.nex\""

echo
echo "********************************"
echo "***** Creating directories *****"
echo "********************************"
# Create the main directory
mkdir -p $basedir/rps11

if [[ $SLURM_ARRAY_TASK_ID -eq 1 ]]; then
    # Create the 3prime1 sites posterior directory
    mkdir -p $basedir/rps11/3prime1/posterior
    echo "datafile <- $data3prime; rnseed <- $post3prime1seed; run_under_prior <- FALSE" > $basedir/rps11/3prime1/posterior/adhoc.Rev
    
    # Create the 3prime1 sites prior directory
    mkdir -p $basedir/rps11/3prime1/prior
    echo "datafile <- $data3prime; rnseed <- $prior3prime1seed; run_under_prior <- TRUE" > $basedir/rps11/3prime1/prior/adhoc.Rev
elif [[ $SLURM_ARRAY_TASK_ID -eq 2 ]]; then
    # Create the 3prime2 sites posterior directory
    mkdir -p $basedir/rps11/3prime2/posterior
    echo "datafile <- $data3prime; rnseed <- $post3prime2seed; run_under_prior <- FALSE" > $basedir/rps11/3prime2/posterior/adhoc.Rev
    
    # Create the 3prime2 sites prior directory
    mkdir -p $basedir/rps11/3prime2/prior
    echo "datafile <- $data3prime; rnseed <- $prior3prime2seed; run_under_prior <- TRUE" > $basedir/rps11/3prime2/prior/adhoc.Rev
elif [[ $SLURM_ARRAY_TASK_ID -eq 3 ]]; then
    # Create the 5prime1 sites posterior directory
    mkdir -p $basedir/rps11/5prime1/posterior
    echo "datafile <- $data5prime; rnseed <- $post5prime1seed; run_under_prior <- FALSE" > $basedir/rps11/5prime1/posterior/adhoc.Rev
    
    # Create the 5prime1 sites prior directory
    mkdir -p $basedir/rps11/5prime1/prior
    echo "datafile <- $data5prime; rnseed <- $prior5prime1seed; run_under_prior <- TRUE" > $basedir/rps11/5prime1/prior/adhoc.Rev
elif [[ $SLURM_ARRAY_TASK_ID -eq 4 ]]; then
    # Create the 5prime2 sites posterior directory
    mkdir -p $basedir/rps11/5prime2/posterior
    echo "datafile <- $data5prime; rnseed <- $post5prime2seed; run_under_prior <- FALSE" > $basedir/rps11/5prime2/posterior/adhoc.Rev
    
    # Create the 5prime2 sites prior directory
    mkdir -p $basedir/rps11/5prime2/prior
    echo "datafile <- $data5prime; rnseed <- $prior5prime2seed; run_under_prior <- TRUE" > $basedir/rps11/5prime2/prior/adhoc.Rev
elif [[ $SLURM_ARRAY_TASK_ID -eq 5 ]]; then
    # Create the all1 posterior directory
    mkdir -p $basedir/rps11/all1/posterior
    echo "datafile <- $dataall; rnseed <- $postall1seed; run_under_prior <- FALSE" > $basedir/rps11/all1/posterior/adhoc.Rev
    
    # Create the all1 prior directory
    mkdir -p $basedir/rps11/all1/prior
    echo "datafile <- $dataall; rnseed <- $priorall1seed; run_under_prior <- TRUE" > $basedir/rps11/all1/prior/adhoc.Rev
else
    # Create the all2 posterior directory
    mkdir -p $basedir/rps11/all2/posterior
    echo "datafile <- $dataall; rnseed <- $postall2seed; run_under_prior <- FALSE" > $basedir/rps11/all2/posterior/adhoc.Rev
    
    # Create the all2 prior directory
    mkdir -p $basedir/rps11/all2/prior
    echo "datafile <- $dataall; rnseed <- $priorall2seed; run_under_prior <- TRUE" > $basedir/rps11/all2/prior/adhoc.Rev
fi

if [[ $SLURM_ARRAY_TASK_ID -eq 1 ]]; then
    echo
    echo "************************************************"
    echo "***** Running RevBayes (3prime1 posterior) *****"
    echo "************************************************"
    # Sample the posterior and compute the prior mean tree
    cd $basedir/rps11/3prime1/posterior 
    $RB $basedir/fig-bloodroot.Rev
    
    echo
    echo "********************************************"
    echo "***** Running RevBayes (3prime1 prior) *****"
    echo "********************************************"
    # Sample the prior and compute the prior mean tree
    cd $basedir/rps11/3prime1/prior
    $RB $basedir/fig-bloodroot.Rev

    echo
    echo "***********************************"
    echo "***** Calculating information *****"
    echo "***********************************"
    cd $basedir/rps11/3prime1 
    rm -rf results
    python3 $basedir/../calcinfo.py --oppath $OP --dirname results --prefix info --skip 1 \
        --postfile posterior/output/rps11.trees --priorfile prior/output/rps11.trees 
    python3 $basedir/../calcinfo.py --oppath $OP --hpdrad --radpct 95 --dirname results --prefix info95 --skip 1 \
        --postfile posterior/output/rps11.trees --priorfile prior/output/rps11.trees 
elif [[ $SLURM_ARRAY_TASK_ID -eq 2 ]]; then
    echo
    echo "************************************************"
    echo "***** Running RevBayes (3prime2 posterior) *****"
    echo "************************************************"
    # Sample the posterior and compute the prior mean tree
    cd $basedir/rps11/3prime2/posterior 
    $RB $basedir/fig-bloodroot.Rev
    
    echo
    echo "********************************************"
    echo "***** Running RevBayes (3prime2 prior) *****"
    echo "********************************************"
    # Sample the prior and compute the prior mean tree
    cd $basedir/rps11/3prime2/prior
    $RB $basedir/fig-bloodroot.Rev

    echo
    echo "***********************************"
    echo "***** Calculating information *****"
    echo "***********************************"
    cd $basedir/rps11/3prime2
    rm -rf results
    python3 $basedir/../calcinfo.py --oppath $OP --dirname results --prefix info --skip 1 \
        --postfile posterior/output/rps11.trees --priorfile prior/output/rps11.trees 
    python3 $basedir/../calcinfo.py --oppath $OP --hpdrad --radpct 95 --dirname results --prefix info95 --skip 1 \
        --postfile posterior/output/rps11.trees --priorfile prior/output/rps11.trees 
elif [[ $SLURM_ARRAY_TASK_ID -eq 3 ]]; then
    echo
    echo "************************************************"
    echo "***** Running RevBayes (5prime1 posterior) *****"
    echo "************************************************"
    # Sample the posterior and compute the prior mean tree
    cd $basedir/rps11/5prime1/posterior 
    $RB $basedir/fig-bloodroot.Rev
    
    echo
    echo "********************************************"
    echo "***** Running RevBayes (5prime1 prior) *****"
    echo "********************************************"
    # Sample the prior and compute the prior mean tree
    cd $basedir/rps11/5prime1/prior
    $RB $basedir/fig-bloodroot.Rev

    echo
    echo "***********************************"
    echo "***** Calculating information *****"
    echo "***********************************"
    cd $basedir/rps11/5prime1 
    rm -rf results
    python3 $basedir/../calcinfo.py --oppath $OP --dirname results --prefix info --skip 1 \
        --postfile posterior/output/rps11.trees --priorfile prior/output/rps11.trees 
    python3 $basedir/../calcinfo.py --oppath $OP --hpdrad --radpct 95 --dirname results --prefix info95 --skip 1 \
        --postfile posterior/output/rps11.trees --priorfile prior/output/rps11.trees 
elif [[ $SLURM_ARRAY_TASK_ID -eq 4 ]]; then
    echo
    echo "************************************************"
    echo "***** Running RevBayes (5prime2 posterior) *****"
    echo "************************************************"
    # Sample the posterior and compute the prior mean tree
    cd $basedir/rps11/5prime2/posterior 
    $RB $basedir/fig-bloodroot.Rev
    
    echo
    echo "********************************************"
    echo "***** Running RevBayes (5prime2 prior) *****"
    echo "********************************************"
    # Sample the prior and compute the prior mean tree
    cd $basedir/rps11/5prime2/prior
    $RB $basedir/fig-bloodroot.Rev

    echo
    echo "***********************************"
    echo "***** Calculating information *****"
    echo "***********************************"
    cd $basedir/rps11/5prime2 
    rm -rf results
    python3 $basedir/../calcinfo.py --oppath $OP --dirname results --prefix info --skip 1 \
        --postfile posterior/output/rps11.trees --priorfile prior/output/rps11.trees 
    python3 $basedir/../calcinfo.py --oppath $OP --hpdrad --radpct 95 --dirname results --prefix info95 --skip 1 \
        --postfile posterior/output/rps11.trees --priorfile prior/output/rps11.trees 
elif [[ $SLURM_ARRAY_TASK_ID -eq 5 ]]; then
    echo
    echo "*********************************************"
    echo "***** Running RevBayes (all1 posterior) *****"
    echo "*********************************************"
    # Sample the posterior and compute the prior mean tree
    cd $basedir/rps11/all1/posterior 
    $RB $basedir/fig-bloodroot.Rev
    
    echo
    echo "*****************************************"
    echo "***** Running RevBayes (all1 prior) *****"
    echo "*****************************************"
    # Sample the prior and compute the prior mean tree
    cd $basedir/rps11/all1/prior
    $RB $basedir/fig-bloodroot.Rev

    echo
    echo "***********************************"
    echo "***** Calculating information *****"
    echo "***********************************"
    cd $basedir/rps11/all1 
    rm -rf results
    python3 $basedir/../calcinfo.py --oppath $OP --dirname results --prefix info --skip 1 \
        --postfile posterior/output/rps11.trees --priorfile prior/output/rps11.trees 
    python3 $basedir/../calcinfo.py --oppath $OP --hpdrad --radpct 95 --dirname results --prefix info95 --skip 1 \
        --postfile posterior/output/rps11.trees --priorfile prior/output/rps11.trees 
else
    echo
    echo "*********************************************"
    echo "***** Running RevBayes (all2 posterior) *****"
    echo "*********************************************"
    # Sample the posterior and compute the prior mean tree
    cd $basedir/rps11/all2/posterior
    $RB $basedir/fig-bloodroot.Rev
    
    echo
    echo "*****************************************"
    echo "***** Running RevBayes (all2 prior) *****"
    echo "*****************************************"
    # Sample the prior and compute the prior mean tree
    cd $basedir/rps11/all2/prior
    $RB $basedir/fig-bloodroot.Rev

    echo
    echo "***********************************"
    echo "***** Calculating information *****"
    echo "***********************************"
    cd $basedir/rps11/all2 
    rm -rf results
    python3 $basedir/../calcinfo.py --oppath $OP --dirname results --prefix info --skip 1 \
        --postfile posterior/output/rps11.trees --priorfile prior/output/rps11.trees 
    python3 $basedir/../calcinfo.py --oppath $OP --hpdrad --radpct 95 --dirname results --prefix info95 --skip 1 \
        --postfile posterior/output/rps11.trees --priorfile prior/output/rps11.trees 
fi
```
