## Supplementary material for "Estimating Bayesian Phylogenetic Information Content Using Geodesic Distances": runmds.sh

```
#!/bin/bash

DIR="$( cd "$( dirname "$0" )" && pwd )"
RED_DOT_SIZE="2.0"

FILL_THREE_PRIME_ONE="#00009b"    # navy
STROKE_THREE_PRIME_ONE="#00009b"  # navy

FILL_THREE_PRIME_TWO="#99f8fb"    # cyan
STROKE_THREE_PRIME_TWO="#000000"  # black

FILL_FIVE_PRIME_ONE="#8d3b34"     # maroon
STROKE_FIVE_PRIME_ONE="#8d3b34"   # maroon

FILL_FIVE_PRIME_TWO="#ed8532"     # orange
STROKE_FIVE_PRIME_TWO="#000000"   # black

python3 ../mds.py \
    --hpdrad \
    --radpct 95 \
    --dirname 'results/mds' \
    --prefix '3a3b-hpd' \
    --basedir "$DIR" \
    --fill '#ff0000' \
    --stroke '#000000' \
    --ptsize $RED_DOT_SIZE \
    --ptstroke 0.1 \
    --trees 'results/rps11/3prime1/posterior/output/rps11.trees' \
    --subsample 500 \
    --subseed 12345 \
    --fill $FILL_THREE_PRIME_ONE \
    --stroke $STROKE_THREE_PRIME_ONE \
    --ptsize 1.0 \
    --ptstroke 0.1 \
    --trees 'results/rps11/3prime2/posterior/output/rps11.trees' \
    --subsample 500 \
    --subseed 54321 \
    --fill $FILL_THREE_PRIME_TWO \
    --stroke $STROKE_THREE_PRIME_TWO \
    --ptsize 1.0 \
    --ptstroke 0.1

python3 ../mds.py \
    --hpdrad \
    --radpct 95 \
    --dirname 'results/mds' \
    --prefix '5a5b-hpd' \
    --basedir "$DIR" \
    --fill '#ff0000' \
    --stroke '#000000' \
    --ptsize $RED_DOT_SIZE \
    --ptstroke 0.1 \
    --trees 'results/rps11/5prime1/posterior/output/rps11.trees' \
    --subsample 500 \
    --subseed 12345 \
    --fill $FILL_FIVE_PRIME_ONE \
    --stroke $STROKE_FIVE_PRIME_ONE \
    --ptsize 1.0 \
    --ptstroke 0.1 \
    --trees 'results/rps11/5prime2/posterior/output/rps11.trees' \
    --subsample 500 \
    --subseed 54321 \
    --fill $FILL_FIVE_PRIME_TWO \
    --stroke $STROKE_FIVE_PRIME_TWO \
    --ptsize 1.0 \
    --ptstroke 0.1

python3 ../mds.py \
    --hpdrad \
    --radpct 95 \
    --dirname 'results/mds' \
    --prefix '3a3b5a5b-hpd' \
    --basedir "$DIR" \
    --fill '#ff0000' \
    --stroke '#000000' \
    --ptsize $RED_DOT_SIZE \
    --ptstroke 0.1 \
    --trees 'results/rps11/3prime1/posterior/output/rps11.trees' \
    --subsample 250 \
    --subseed 12345 \
    --fill $FILL_THREE_PRIME_ONE \
    --stroke $STROKE_THREE_PRIME_ONE \
    --ptsize 1.0 \
    --ptstroke 0.1 \
    --trees 'results/rps11/3prime2/posterior/output/rps11.trees' \
    --subsample 250 \
    --subseed 12345 \
    --fill $FILL_THREE_PRIME_TWO \
    --stroke $STROKE_THREE_PRIME_TWO \
    --ptsize 1.0 \
    --ptstroke 0.1 \
    --trees 'results/rps11/5prime1/posterior/output/rps11.trees' \
    --subsample 250 \
    --subseed 54321 \
    --fill $FILL_FIVE_PRIME_ONE \
    --stroke $STROKE_FIVE_PRIME_ONE \
    --ptsize 1.0 \
    --ptstroke 0.1 \
    --trees 'results/rps11/5prime2/posterior/output/rps11.trees' \
    --subsample 250 \
    --subseed 54321 \
    --fill $FILL_FIVE_PRIME_TWO \
    --stroke $STROKE_FIVE_PRIME_TWO \
    --ptsize 1.0 \
    --ptstroke 0.1

# python3 ../mds.py \
#     --hpdrad \
#     --radpct 95 \
#     --dirname 'results/mds' \
#     --prefix '3a5a-hpd' \
#     --basedir "$DIR" \
#     --fill '#ff0000' \
#     --stroke '#000000' \
#     --ptsize $RED_DOT_SIZE \
#     --ptstroke 0.1 \
#     --trees 'results/rps11/3prime1/posterior/output/rps11.trees' \
#     --subsample 500 \
#     --subseed 12345 \
#     --fill $FILL_THREE_PRIME_ONE \
#     --stroke $STROKE_THREE_PRIME_ONE \
#     --ptsize 1.0 \
#     --ptstroke 0.1 \
#     --trees 'results/rps11/5prime1/posterior/output/rps11.trees' \
#     --subsample 500 \
#     --subseed 54321 \
#     --fill $FILL_FIVE_PRIME_ONE \
#     --stroke $STROKE_FIVE_PRIME_ONE \
#     --ptsize 1.0 \
#     --ptstroke 0.1
# 
# python3 ../mds.py \
#     --hpdrad \
#     --radpct 95 \
#     --dirname 'results/mds' \
#     --prefix '3a5b-hpd' \
#     --basedir "$DIR" \
#     --fill '#ff0000' \
#     --stroke '#000000' \
#     --ptsize $RED_DOT_SIZE \
#     --ptstroke 0.1 \
#     --trees 'results/rps11/3prime1/posterior/output/rps11.trees' \
#     --subsample 500 \
#     --subseed 12345 \
#     --fill $FILL_THREE_PRIME_ONE \
#     --stroke $STROKE_THREE_PRIME_ONE \
#     --ptsize 1.0 \
#     --ptstroke 0.1 \
#     --trees 'results/rps11/5prime2/posterior/output/rps11.trees' \
#     --subsample 500 \
#     --subseed 54321 \
#     --fill $FILL_FIVE_PRIME_TWO \
#     --stroke $STROKE_FIVE_PRIME_TWO \
#     --ptsize 1.0 \
#     --ptstroke 0.1
# 
# python3 ../mds.py \
#     --hpdrad \
#     --radpct 95 \
#     --dirname 'results/mds' \
#     --prefix '3b5a-hpd' \
#     --basedir "$DIR" \
#     --fill '#ff0000' \
#     --stroke '#000000' \
#     --ptsize $RED_DOT_SIZE \
#     --ptstroke 0.1 \
#     --trees 'results/rps11/3prime2/posterior/output/rps11.trees' \
#     --subsample 500 \
#     --subseed 12345 \
#     --fill $FILL_THREE_PRIME_TWO \
#     --stroke $STROKE_THREE_PRIME_TWO \
#     --ptsize 1.0 \
#     --ptstroke 0.1 \
#     --trees 'results/rps11/5prime1/posterior/output/rps11.trees' \
#     --subsample 500 \
#     --subseed 54321 \
#     --fill $FILL_FIVE_PRIME_ONE \
#     --stroke $STROKE_FIVE_PRIME_ONE \
#     --ptsize 1.0 \
#     --ptstroke 0.1
# 
# python3 ../mds.py \
#     --hpdrad \
#     --radpct 95 \
#     --dirname 'results/mds' \
#     --prefix '3b5b-hpd' \
#     --basedir "$DIR" \
#     --fill '#ff0000' \
#     --stroke '#000000' \
#     --ptsize $RED_DOT_SIZE \
#     --ptstroke 0.1 \
#     --trees 'results/rps11/3prime2/posterior/output/rps11.trees' \
#     --subsample 500 \
#     --subseed 12345 \
#     --fill $FILL_THREE_PRIME_TWO \
#     --stroke $STROKE_THREE_PRIME_TWO \
#     --ptsize 1.0 \
#     --ptstroke 0.1 \
#     --trees 'results/rps11/5prime2/posterior/output/rps11.trees' \
#     --subsample 500 \
#     --subseed 54321 \
#     --fill $FILL_FIVE_PRIME_TWO \
#     --stroke $STROKE_FIVE_PRIME_TWO \
#     --ptsize 1.0 \
#     --ptstroke 0.1
```
