## Supplementary material for "Estimating Bayesian Phylogenetic Information Content Using Geodesic Distances": summarize.py

```
from scipy.stats import describe
import argparse, os, sys, math, subprocess, glob, re

parser = argparse.ArgumentParser(
    formatter_class=argparse.RawDescriptionHelpFormatter,
    description='''Calculates dissonance for all pairwise comparisons.''',
    epilog='''Be sure OPpath is the correct path to the OP executable. OP may be obtained from https://github.com/plewis/OP.''')
parser.add_argument('--dir', required=True, help='directory containing rps11 directory')
parser.add_argument('--precision', type=int, default=5, help='number of digits precision')
parser.add_argument('--hpd', default=False, action='store_true', help='report results for HPD method rather than the default RAD method')
settings = vars(parser.parse_args())

OPpath = 'op'
dirname   = settings['dir']
precision = settings['precision']
hpdmethod = settings['hpd']

# Get path of current directory
current_working_dir = os.getcwd()

# Create a directory to hold results
mean_diffs_dir = os.path.join(dirname, 'mean-diffs')
if os.path.exists(mean_diffs_dir):
    sys.exit('Please move or rename directory "%s" and try running summarize.py again' % mean_diffs_dir)
os.mkdir(mean_diffs_dir)

outf = None

def output(s = None):
    global outf
    if s is None:
        print()
        outf.write('\n')
    else:
        print(s)
        outf.write('%s\n' % s)

# Open the output file
outfn = os.path.join(dirname, 'pairwise-dissonance.txt')
if os.path.exists(outfn):
    sys.exit('Please move or rename the file "%s" and try running summarize.py again' % outfn)
outf = open(outfn, 'w')

def extractInfo(fn):
    stuff = open(fn, 'r').read()

    m = re.search(r'([0-9]+) = posterior sample size', stuff, re.M | re.S)
    assert m is not None, 'Could not extract posterior sample size from file "%s"' % fn
    post_sample_size = float(m.group(1))

    m = re.search(r'([0-9]+) = prior sample size', stuff, re.M | re.S)
    assert m is not None, 'Could not extract prior sample size from file "%s"' % fn
    prior_sample_size = float(m.group(1))

    m = re.search(r'([.0-9]+) = posterior mean tree length', stuff, re.M | re.S)
    assert m is not None, 'Could not extract posterior mean tree length from file "%s"' % fn
    post_mean_TL = float(m.group(1))

    m = re.search(r'([.0-9]+) = prior mean tree length', stuff, re.M | re.S)
    assert m is not None, 'Could not extract prior mean tree length from file "%s"' % fn
    prior_mean_TL = float(m.group(1))

    m = re.search(r'([.0-9]+) = posterior variance', stuff, re.M | re.S)
    assert m is not None, 'Could not extract posterior variance from file "%s"' % fn
    post_var = float(m.group(1))

    m = re.search(r'([.0-9]+) = prior variance', stuff, re.M | re.S)
    assert m is not None, 'Could not extract prior variance from file "%s"' % fn
    prior_var = float(m.group(1))

    m = re.search(r'([.0-9]+) = posterior scaler', stuff, re.M | re.S)
    assert m is not None, 'Could not extract posterior scaler from file "%s"' % fn
    post_scaler = float(m.group(1))

    m = re.search(r'([.0-9]+) = prior scaler', stuff, re.M | re.S)
    assert m is not None, 'Could not extract prior scaler from file "%s"' % fn
    prior_scaler = float(m.group(1))

    m = re.search(r'([.0-9]+) = posterior 95% radius', stuff, re.M | re.S)
    assert m is not None, 'Could not extract posterior 95% radius from file "%s"' % fn
    post_radius = float(m.group(1))

    m = re.search(r'([.0-9]+) = prior 95% radius', stuff, re.M | re.S)
    assert m is not None, 'Could not extract prior 95% radius from file "%s"' % fn
    prior_radius = float(m.group(1))

    m = re.search(r'([-.0-9]+) = LCRrad', stuff, re.M | re.S)
    assert m is not None, 'Could not extract LCRrad from file "%s"' % fn
    LCRrad = float(m.group(1))

    m = re.search(r'([-.0-9]+) = information as percentage', stuff, re.M | re.S)
    assert m is not None, 'Could not extract information as percentage from file "%s"' % fn
    Irad = float(m.group(1))

    return {
        'post_sample_size': post_sample_size,
        'prior_sample_size': prior_sample_size,
        'post_mean_TL': post_mean_TL,
        'prior_mean_TL': prior_mean_TL,
        'post_var': post_var,
        'prior_var': prior_var,
        'post_scaler': post_scaler,
        'prior_scaler': prior_scaler,
        'post_radius': post_radius,
        'prior_radius': prior_radius,
        'LCRrad': LCRrad,
        'Irad': Irad
    }
        
def calcPairwiseDissonance(i, j, analyses):
    first = analyses[i]
    second = analyses[j]
    
    infofn = hpdmethod and 'info95-output.txt' or 'info-output.txt'
    
    fn1     = os.path.join(dirname, 'rps11', first, 'results', infofn)
    info1   = extractInfo(fn1)
    n1      = info1['post_sample_size']
    lcrrad1 = info1['LCRrad']
    irad1   = info1['Irad']
    var1    = info1['post_var']
    radius1 = info1['post_radius']
    
    fn2     = os.path.join(dirname, 'rps11', second, 'results', infofn)
    info2   = extractInfo(fn2)
    n2      = info2['post_sample_size']
    lcrrad2 = info2['LCRrad']
    irad2   = info2['Irad']
    var2    = info2['post_var']
    radius2 = info2['post_radius']

    # Run OP to calculate difference in mean trees
    infodir = hpdmethod and 'info95' or 'info'
    mean1fn = os.path.join(dirname, 'rps11', first,  'results', infodir, 'op-post.tre')
    mean2fn = os.path.join(dirname, 'rps11', second, 'results', infodir, 'op-post.tre')
    prefix = os.path.join(mean_diffs_dir, 'pair-%d-%d' % (i, j))
    cmds = [OPpath, 
        '--reftree', mean1fn,
        '--refskip', '0',
        '--refscale', '%g' % info1['post_scaler'],
        '--treefile', mean2fn,
        '--skip', '0',
        '--scale', '%g' % info2['post_scaler'],
        '--prefix', prefix
    ]
    completed_process = subprocess.run(cmds, cwd=current_working_dir, capture_output=True)
    try:
        completed_process.check_returncode()
    except sub.CalledProcessError:
        output('\nAn error occurred in calcPairwiseDissonance while processing pair %d (%s) vs %d (%s)' % (i, analyses[i], j, analyses[j]))
        output('Standard error:')
        output(completed_process.stderr)
        output('Standard output:')
        output(completed_process.stdout)
        sys.exit('Aborted.')
        
    # Extract mean difference from treefile saved by OP
    treefn = os.path.join(dirname, 'mean-diffs', 'pair-%d-%d.tre' % (i, j))
    stuff = open(treefn, 'r').read()
    m = re.search(r'bhvdist = ([-.0-9]+)', stuff, re.M | re.S)
    assert m is not None, 'Could not extract bhvdist from treefile "%s"' % treefn
    meandiff = float(m.group(1))
    
    # Calculate average information content
    numer = float(n1 - 1)*irad1 + float(n2 - 1)*irad2
    denom = float(n1 + n2 - 2)
    iavg = numer/denom
        
    diss = None
    if radius1 > 0.0 and radius2 > 0.0:
        numer = float(n1 - 1)*pow(radius1,2) + float(n2 - 1)*pow(radius2,2)
        denom = float(n1 + n2 - 2)
        pooled_radius = math.sqrt(numer/denom)
        diss = meandiff/pooled_radius
    return (n1, radius1, lcrrad1, irad1, n2, radius2, lcrrad2, irad2, iavg, meandiff, pooled_radius, diss)

if __name__ == '__main__':
    # Show the specified settings
    output('Settings:')
    output('  dir       = "%s"' % dirname)
    output('  precision = %d' % precision)
    output('  hpd       = %s' % (hpdmethod and "yes" or "no",))
    output()
    
    analyses = ['3prime1', '3prime2', '5prime1', '5prime2', 'all1', 'all2']
    nanalyses = len(analyses)
    analysis_dict = {}
    output('%12s %12s %12s %12s %12s %12s %12s %12s %12s %12s %12s %12s' % ('first', 'second', 'n_i', 'radius_i', 'info_i', 'n_j', 'radius_j', 'info_j', 'avg_info', 'mean_diff', 'pooled_radius', 'dissonance'))
    for i in range(nanalyses - 1):
        for j in range(i, nanalyses):
            n1, radius1, lcr1, i1, n2, radius2, lcr2, i2, iavg, mean_diff, pooled_radius, diss = calcPairwiseDissonance(i, j, analyses)
            if diss is not None:
                output('%12s %12s %12d %12.5f %12.5f %12d %12.5f %12.5f %12.5f %12.5f %12.5f %12.5f' % (analyses[i], analyses[j], n1, radius1, i1, n2, radius2, i2, iavg, mean_diff, pooled_radius, diss))
            else:
                output('%12s %12s %12d %12.5f %12.5f %12d %12.5f %12.5f %12.5f %12.5f %12s %12s' % (analyses[i], analyses[j], n1, radius1, i1, n2, radius2, i2, iavg, mean_diff, '---', '---'))
    
    outf.close()
```
