## Supplementary material for "Estimating Bayesian Phylogenetic Information Content Using Geodesic Distances": calcinfo.py

```
import argparse,re,sys,os,math,subprocess as sub
from math import log

parser = argparse.ArgumentParser(
    formatter_class=argparse.RawDescriptionHelpFormatter,
    description='''Expects a directory, postfile, priorfile, and a prefix to be provided.
The directory specified by dirname will be created if it does not exist. This directory
will contain the results.''',
    epilog='''No epilog''')
parser.add_argument('--oppath', default='op', help='path to OP executable')
parser.add_argument('--hpdrad', default=False, action='store_true', help='create mean tree and radius from HPD credible set of trees')
parser.add_argument('--radpct', default=95, type=int, help='radius includes this percentage of trees closest to mean')
parser.add_argument('--skip', default=1, type=int, help='number of trees to skip')
parser.add_argument('--epsilon', default=0.00001, type=float, help='stopping criterion for Frechet mean calculation')
parser.add_argument('--dirname', required=True, help='directory in which to store results')
parser.add_argument('--prefix', default='op-results', help='forms beginning of result file paths')
parser.add_argument('--postfile', default=None, help='file name of posterior tree file')
parser.add_argument('--priorfile', default=None, help='file name of prior tree file')
parser.add_argument('--file1', default=None, help='file name of first tree file in dissonance analysis')
parser.add_argument('--mean1', default=None, help='file name of first mean tree file in dissonance analysis')
parser.add_argument('--info1', default=None, help='output file from analysis of information in first dataset')
parser.add_argument('--scaler1', default=1.0, type=float, help='scaler for trees in first tree file')
parser.add_argument('--file2', default=None, help='file name of second tree file in dissonance analysis')
parser.add_argument('--mean2', default=None, help='file name of second mean tree file in dissonance analysis')
parser.add_argument('--info2', default=None, help='output file from analysis of information in first dataset')
parser.add_argument('--scaler2', default=1.0, type=float, help='scaler for trees in second tree file')
settings = vars(parser.parse_args())

path_to_OP = settings['oppath']
hpdrad     = settings['hpdrad']
radpct     = settings['radpct']
skip       = settings['skip']
epsilon    = settings['epsilon']
dirname    = settings['dirname']
prefix     = settings['prefix']
post_file_path  = settings['postfile']
prior_file_path = settings['priorfile']
file1_path = settings['file1']
mean1_path = settings['mean1']
info1_path = settings['info1']
scaler1    = settings['scaler1']
file2_path = settings['file2']
mean2_path = settings['mean2']
info2_path = settings['info2']
scaler2    = settings['scaler2']

radfrac = 0.01*radpct

# Ensure prefix contains no spaces
prefix,nsubs = re.subn(r'\s+', r'_', prefix)
if nsubs > 0:
    print('Warning: whitespace in supplied prefix was converted to underscores')

script_dir = os.path.dirname(os.path.realpath(sys.argv[0]))
current_working_dir = os.getcwd()

def ensureFilePathExists(fpath):
    if not os.path.exists(fpath):
        sys.exit('\nExpecting this file path to exist: "%s"' % fpath)
    #print('OK, path exists: "%s"' % fpath)

def ensureFilePathDoesNotExist(fpath):
    if os.path.exists(fpath):
        sys.exit('\nPlease delete or rename the file "%s" before running this program' % fpath)
    # print('OK, path does not exist: "%s"' % fpath)

# Create the specified results directory if it does not already exist
if not os.path.exists(dirname):
    os.mkdir(dirname)

# Ensure that an analysis with this prefix has not already been run
outfn = os.path.join(dirname, '%s-output.txt' % prefix)
ensureFilePathDoesNotExist(outfn)

outf = None

def output(s = None):
    global outf
    if s is None:
        print()
        outf.write('\n')
    else:
        print(s)
        outf.write('%s\n' % s)

print()        
print('Settings:')
print('  hpdrad           = "%s"' % (hpdrad and 'yes' or 'no',))
print('  radpct           = %d' % radpct)
print('  path_to_OP       = "%s"' % path_to_OP)
print('  script_dir       = "%s"' % script_dir)
print('  cwd              = "%s"' % current_working_dir)
print('  skip             = %d' % skip)
print('  epsilon          = %g' % epsilon)
print('  dirname          = "%s"' % dirname)
print('  prefix           = "%s"' % prefix)
print('Environment:')
print('  script_dir          = "%s"' % script_dir)
print('  current_working_dir = "%s"' % current_working_dir)

calculating_information = False
if not file1_path is None:
    print('  file1_path   = "%s"' % file1_path)
    ensureFilePathExists(file1_path)

    print('  file2_path   = "%s"' % file2_path)
    ensureFilePathExists(file2_path)

    print('  mean1_path   = "%s"' % mean1_path)
    ensureFilePathExists(mean1_path)

    print('  mean2_path   = "%s"' % mean2_path)
    ensureFilePathExists(mean2_path)

    print('  info1_path   = "%s"' % info1_path)
    ensureFilePathExists(info1_path)

    print('  info2_path   = "%s"' % info2_path)
    ensureFilePathExists(info2_path)

    print('  scaler1      = "%s"' % scaler1)
    print('  scaler2      = "%s"' % scaler2)
else:
    calculating_information = True
    ensureFilePathExists(post_file_path)
    ensureFilePathExists(prior_file_path)
    print('  post_file_path   = "%s"' % post_file_path)
    print('  prior_file_path  = "%s"' % prior_file_path)

def extractTLRadius(fn):
    # fn should be the path to the R file containing OP Frechet mean results
    # For example, the first few lines of the file should look like this:
    # # (Sphaeroplea_robusta_UTEX2350:0.001186000,Ankyra_judai_SAG17_84:0.021105091,(((((((Tumidella_tumida_SAG2265:0.063690636,Dictyochloris_fragrans_UTEX127:0.054760970):0.000087384,(Planktosphaeria_gelatinosa_SAG262_1b:0.034971394,Bracteamorpha_trainori_CC1VF5A:0.049583848):0.000042047):0.000004486,(Bracteacoccus_minor_UTEX66:0.018045030,Bracteacoccus_aerius_UTEX1250:0.018885212):0.020902727):0.000002165,((Radiococcus_polycoccus_SAG217_1c:0.040317939,(Planktosphaeria_botryoides_UTEXLB951:0.009633788,Planktosphaeria_texensis_UTEX1241:0.012156727):0.042179667):0.020826212,Schizochlamys_gelatinosa_SAG66_94:0.043422182):0.019274879):0.000006257,((((Pseudomuriella_schumacherensis_SAG2137:0.016491667,Pseudomuriella_engadinensis_UTEX58:0.013051546):0.030039000,(Mychonastes_homosphaera_CAUPH6502:0.011134576,Mychonastes_jurisii_SAG3798:0.011493697):0.083522576):0.000019037,(Rotundella_rotunda_WJT54VFNP11:0.011019758,Rotundella_sp_ZNP1VF31:0.019132515):0.074008455):0.000007615,Chromochloris_zofingiensis_UTEX56:0.027395091):0.000000614):0.000043396,((((Kirchneriella_aperta_SAG2004:0.020551061,Ankistrodesmus_falcatus_UTEX101:0.023059485):0.013917000,Ourococcus_multisporus:0.036963333):0.021234545,Dictyococcus_varians_UTEXLB62:0.105211637):0.027416613,(((Hariotina_reticulata_UTEX1365:0.032551121,Scenedesmus_rotundus_SEV3VF49:0.017471757):0.007405928,Scenedesmus_obliquus_UTEX393:0.017483576):0.006359791,(((Parapediastrum_biradiatum_UTEX37:0.016587061,Pediastrum_duplex_UTEX1364:0.014214394):0.015451394,(Neochloris_aquatica_UTEX138:0.020831242,(Characiopodium_hindakii:0.011136758,Botryococcus_sudeticus:0.008455455):0.007515939):0.031896334):0.001231364,Chlorotetraedron_incus_SAG4381:0.010722970):0.014301394):0.001061503):0.000092667):0.000148220,Schroederia_setigera_UTEXLB2454:0.051171969):0.038260333)
    # # sample size = 10000
    # # variance = 0.006687729
    # # tree length = 1.341149028
    # # iterations = 32
    # # frechet-e = 0.01
    # # frechet-k = 1000000
    # # frechet-n = 10
    # # pseudorandom number seed = 1
    # # q25 = -0.079683907
    # # q75 = 0.079683907
    # # IQR = 0.159367815
    # # KDE sigma = 0.081776490
    # # KDE bandwidth = 0.010154650
    # # 95% HPD lower = 0.058685569
    # # 95% HPD upper = 0.102040726
    # # 95% radius = 0.101098573
    # # smallest distance = 0.051341071
    # # largest distance = 0.144040701
    ensureFilePathExists(fn)
    stuff = open(fn, 'r').read()
    
    # Extract TL
    m = re.search(r'# tree length = ([.0-9]+)', stuff)
    assert m is not None
    TL = float(m.group(1))

    # Extract variance
    m = re.search(r'# variance = ([.0-9]+)', stuff)
    assert m is not None
    variance = float(m.group(1))

    # Extract radius
    m = re.search(r'# [.0-9]+% radius = ([.0-9]+)', stuff)
    assert m is not None
    radius = float(m.group(1))
    
    # Extract samplesize
    m = re.search(r'# sample size = ([0-9]+)', stuff)
    assert m is not None
    samplesize = float(m.group(1))
        
    return {'samplesize': samplesize, 'TL': TL, 'radius':radius, 'variance': variance}
    
def calcInfoForPair(post_file_path, prior_file_path):
    # Create directory to hold results
    results_dir = os.path.join(current_working_dir,dirname,'%s' % prefix)
    try:
        os.mkdir(results_dir)
    except FileExistsError:
        sys.exit('\nPlease delete or rename the directory "%s" before running this program' % results_dir)
    except FileNotFoundError:
        sys.exit('\nCould not find some part of the path "%s"' % results_dir)

    print('Calculating information for posterior (%s)...' % post_file_path)
    
    # Set up command line for OP to calculate info for post_file_path
    cmds = [path_to_OP]
    if hpdrad:
        cmds.append('--hpdrad')
        cmds.append('--radpct')
        cmds.append('%d' % radpct)
    cmds.append('--treefile')
    cmds.append(post_file_path)
    cmds.append('--skip')
    cmds.append('%d' % skip)
    cmds.append('--frechetmean')
    cmds.append('--frechet-e')
    cmds.append('%g' % epsilon)
    cmds.append('--prefix')
    cmds.append(os.path.join(results_dir, 'op-post'))
            
    # Run OP to calculate Frechet mean for posterior sample
    completed_process = sub.run(cmds, cwd=current_working_dir, capture_output=True)
    try:
        completed_process.check_returncode()
    except sub.CalledProcessError:
        output('\nAn error occurred in calcInfoForPair while processing file "%s"' % post_file_path)
        output('Standard error:')
        output(completed_process.stderr)
        output('Standard output:')
        output(completed_process.stdout)
        sys.exit('Aborted.')
            
    tmp = extractTLRadius(os.path.join(current_working_dir, dirname, prefix, 'op-post.R'))
    post_sample_size = tmp['samplesize']
    post_tl          = tmp['TL']
    post_radius      = tmp['radius']
    post_variance    = tmp['variance']
    
    print('Calculating information for prior (%s)...' % prior_file_path)

    # Set up command line for OP to calculate info for prior_file_path
    cmds = [path_to_OP]
    if hpdrad:
        cmds.append('--hpdrad')
        cmds.append('--radpct')
        cmds.append('%d' % radpct)
    cmds.append('--treefile')
    cmds.append(prior_file_path)
    cmds.append('--skip')
    cmds.append('%d' % skip)
    cmds.append('--frechetmean')
    cmds.append('--frechet-e')
    cmds.append('%g' % epsilon)
    cmds.append('--prefix')
    cmds.append(os.path.join(results_dir, 'op-prior'))
    
    # Run OP to calculate Frechet mean for prior sample
    completed_process = sub.run(cmds, cwd=current_working_dir, capture_output=True)
    try:
        completed_process.check_returncode()
    except sub.CalledProcessError:
        output('\nAn error occurred in calcInfoForPair while processing file "%s"' % prior_file_path)
        output('Standard error:')
        output(completed_process.stderr)
        output('Standard output:')
        output(completed_process.stdout)
        sys.exit('Aborted.')
    
    tmp = extractTLRadius(os.path.join(current_working_dir, dirname, prefix, 'op-prior.R'))
    prior_sample_size = tmp['samplesize']
    prior_tl          = tmp['TL']
    prior_radius      = tmp['radius']
    prior_variance    = tmp['variance']

    post_scaler  = 1.0/post_tl
    prior_scaler = 1.0/prior_tl
    if prior_radius > 0.0 and prior_scaler > 0.0 and post_radius > 0.0 and post_scaler > 0.0:
        # All is well
        lcr_rad = math.log(prior_radius) + math.log(prior_scaler) - math.log(post_radius) - math.log(post_scaler)
        info_pct = 100.0*(1 - math.exp(-lcr_rad))
        return {
            'postSampleSize':post_sample_size, 
            'priorSampleSize':prior_sample_size, 
            'postTL':post_tl, 
            'priorTL':prior_tl, 
            'postScaler':post_scaler, 
            'priorScaler':prior_scaler, 
            'postRadius':post_radius, 
            'priorRadius':prior_radius, 
            'postVariance':post_variance, 
            'priorVariance':prior_variance, 
            'LCRrad':lcr_rad, 
            'InfoPct':info_pct
        }
    else:
        # Report problem
        prior_radius_ok = prior_radius > 0.0
        prior_scaler_ok = prior_scaler > 0.0
        post_radius_ok  = post_radius > 0.0
        post_scaler_ok  = post_scaler > 0.0
        output('prior_radius = %g (%s)' % (prior_radius, prior_radius_ok and 'Ok' or 'bad'))
        output('prior_scaler = %g (%s)' % (prior_scaler, prior_scaler_ok and 'Ok' or 'bad'))
        output('post_radius  = %g (%s)' % (post_radius,  post_radius_ok  and 'Ok' or 'bad'))
        output('post_scaler  = %g (%s)' % (post_scaler,  post_scaler_ok  and 'Ok' or 'bad'))
        sys.exit('Aborted.')

def calcOverlap(target, meantree, radius, target_scaler, mean_scaler, samplesize):
    # Create directory to hold results
    output('Calculating distance between mean tree ("%s")' % meantree)
    output('  and each tree in the target tree file ("%s")' % target)
    output('  (but skipping first %d trees in target)' % skip)
    
    # Set up command line for OP to calculate info for post_file_path
    cmds = [path_to_OP]
    cmds.append('--hpdrad')
    cmds.append('--radpct')
    cmds.append('%d' % radpct)
    
    cmds.append('--reftree')
    cmds.append(meantree)
    cmds.append('--refskip')
    cmds.append('0')
    cmds.append('--refscale')
    cmds.append('%g' % mean_scaler)
    
    cmds.append('--treefile')
    cmds.append(target)
    cmds.append('--skip')
    cmds.append('%d' % skip)
    cmds.append('--scale')
    cmds.append('%g' % target_scaler)
    
    cmds.append('--prefix')
    cmds.append(os.path.join(results_dir, 'op-refdist'))
    
    # Run OP to calculate reference distances
    completed_process = sub.run(cmds, cwd=current_working_dir, capture_output=True)
    try:
        completed_process.check_returncode()
    except sub.CalledProcessError:
        output('\nAn error occurred in calcOverlap while processing file "%s"' % target)
        output('\nCurrent working directory:')
        output('%s' % current_working_dir)
        output('\nStandard error:')
        output('%s' % completed_process.stderr.decode('utf-8').strip())
        output('\nStandard output:')
        output('%s' % completed_process.stdout.decode('utf-8').strip())
        output('\nCommand:')
        output('%s' % ' '.join(cmds))
        sys.exit('Aborted.')
    
    # Open the treefile containing sorted trees and their distance from the meantree    
    refdist_treefile = os.path.join(current_working_dir, dirname, prefix, 'op-refdist.tre')
    ensureFilePathExists(refdist_treefile)
    stuff = open(refdist_treefile, 'r').read()
    
    # Count number of trees with distance < mean_scaler*radius
    scaled_radius = mean_scaler*radius
    dists = re.findall(r'tree t\d+ \[bhvdist = ([.0-9]+)\]', stuff, re.M | re.S)
    ndists = len(dists)
    assert samplesize == ndists, 'Expecting %d distances in "%s" but found %d instead' % (samplesize, refdist_treefile, ndists)
    n = 0
    for dstr in dists:
        d = float(dstr)
        if d < scaled_radius:
            n += 1
    frac = float(n)/samplesize
        
    return frac
            
if __name__ == '__main__':
    outf = open(outfn, 'w')
    output('Information results:')
    if hpdrad:
        output('  Information calculation based on %d%% HPD credible set:' % radpct)
    else:
        output('  Information calculation based on all sampled trees:')
    if calculating_information:
        # post_file_path and prior_file_path were specified
        results = calcInfoForPair(post_file_path, prior_file_path)
        output('  %12d = posterior sample size' % results['postSampleSize'])
        output('  %12d = prior sample size' % results['priorSampleSize'])
        output('  %12.9f = posterior mean tree length' % results['postTL'])
        output('  %12.9f = prior mean tree length' % results['priorTL'])
        output('  %12.9f = posterior variance' % results['postVariance'])
        output('  %12.9f = prior variance' % results['priorVariance'])
        output('  %12.9f = posterior scaler' % results['postScaler'])
        output('  %12.9f = prior scaler' % results['priorScaler'])
        output('  %12.9f = posterior %d%% radius' % (results['postRadius'], radpct))
        output('  %12.9f = prior %d%% radius' % (results['priorRadius'], radpct))
        output('  %12.9f = LCRrad' % results['LCRrad'])
        output('  %12.9f = information as percentage' % results['InfoPct'])
    else:
        results_dir = os.path.join(current_working_dir,dirname,'%s' % prefix)
        try:
            os.mkdir(results_dir)
        except FileExistsError:
            sys.exit('Please delete or rename the directory "%s" before running this program' % results_dir)
        except FileNotFoundError:
            sys.exit('Could not find some part of the path "%s"' % results_dir)
            
        # Extract sample size and radius from info1_path
        tmp = extractTLRadius(info1_path)
        samplesize1 = tmp['samplesize']
        treelength1 = tmp['TL']
        radius1     = tmp['radius']
        
        # output('*** debug ***: info1_path  = %s' % info1_path)
        # output('*** debug ***: samplesize1 = %g' % samplesize1)
        # output('*** debug ***: treelength1 = %g' % treelength1)
        # output('*** debug ***: radius1     = %g' % radius1)
        
        # Extract sample size and radius from info2_path
        tmp = extractTLRadius(info2_path)
        samplesize2 = tmp['samplesize']
        treelength2 = tmp['TL']
        radius2     = tmp['radius']

        # output('*** debug ***: info2_path  = %s' % info2_path)
        # output('*** debug ***: samplesize2 = %g' % samplesize2)
        # output('*** debug ***: treelength2 = %g' % treelength2)
        # output('*** debug ***: radius2     = %g' % radius2)
        
        # Calculating fraction of trees in file1_path that are closer than radius2 to mean2_path
        frac_one_inside_two = calcOverlap(file1_path, mean2_path, radius2, scaler1, scaler2, samplesize1)

        # Calculating fraction of trees in file2_path that are closer than radius1 to mean1_path
        frac_two_inside_one = calcOverlap(file2_path, mean1_path, radius1, scaler2, scaler1, samplesize2)
                
        highest_fraction = frac_one_inside_two
        if frac_two_inside_one > highest_fraction:
            highest_fraction = frac_two_inside_one
            
        # Highest fraction ranges from 0.0 (no overlap) to 
        # about 0.95 (overlap expected if two samples are from
        # the same distribution)
        # highest_fraction  dissonance = (0.95 - highest_fraction)/0.95
        #             0.00         1.0 = (0.95 - 0.0)/0.95
        #             0.95         0.0 = (0.95 - 0.95)/0.95
        corrected_dissonance = (radfrac - highest_fraction)/radfrac
        dissonance = 1.0 - highest_fraction
        
        output('  %12d = file1 sample size' % samplesize1)
        output('  %12d = file1 sample size' % samplesize2)
        output('  %12.9f = frac_one_inside_two' % frac_one_inside_two)
        output('  %12.9f = frac_two_inside_one' % frac_two_inside_one)
        output('  %12.9f = information dissonance' % dissonance)
        #output('  %12.9f = information dissonance (corrected)' % corrected_dissonance)

    outf.close()
    
```
