## Supplementary material for "Estimating Bayesian Phylogenetic Information Content Using Geodesic Distances": mds.py

```
import argparse,re,sys,os,subprocess

parser = argparse.ArgumentParser(
    formatter_class=argparse.RawDescriptionHelpFormatter,
    description='''Creates an ordination using multidimensional scaling.''',
    epilog='''Before running, ensure that the variable "path_to-OP" 
at the top of the "calcinfo.py" file contains the 
fully-specified path to the OP executable''')
parser.add_argument('--dirname',   required=True,                              help='directory in which to store results')
parser.add_argument('--basedir',   required=True,                              help='base directory (--trees paths are relative to this')
parser.add_argument('--hpdrad',    dest='hpdrad',        action='store_true',  help='whether to retain only the radpct highest posterior samples')
parser.add_argument('--radpct',    default=95,           type=int,             help='the percentage of highest posterior samples to retain if hpdrad specified')
parser.add_argument('--skip',      default=1,            type=int,             help='number of trees to skip in each tree file')
parser.add_argument('--noscale',   dest='doscale',       action='store_false', help='do not rescale any trees')
parser.add_argument('--noop',      dest='doop',          action='store_false', help='do not rescale any trees')
parser.add_argument('--prefix',    default='op-results',                       help='forms beginning of result file paths')
parser.add_argument('--subsample', default=None,         action='append',      help='number of trees to randomly sample from tree file')
parser.add_argument('--subseed',   default=None,         action='append',      help='pseudorandom number seed (used in subsampling trees')
parser.add_argument('--fill',      default=None,         action='append',      help='fill color for dot representing tree in MDS plot')
parser.add_argument('--stroke',    default=None,         action='append',      help='stroke color for dot representing tree in MDS plot')
parser.add_argument('--ptsize',    default=None,         action='append',      help='point size for dot representing tree in MDS plot')
parser.add_argument('--ptstroke',  default=None,         action='append',      help='point stroke color for dot representing tree in MDS plot')
parser.add_argument('--trees',     default=None,         action='append',      help='file path (relative to --basedir) of tree file (can specify multiple values)')
settings = vars(parser.parse_args())

hpdrad    = settings['hpdrad']
radpct    = settings['radpct']
skip      = settings['skip']
dirname   = settings['dirname']
basedir   = settings['basedir']
prefix    = settings['prefix']
trees     = settings['trees']
doscale   = settings['doscale']
runOP     = settings['doop']

oppath = 'op'
    
ntrees  = len(trees)
assert ntrees > 0, 'Must specify at least one tree file (using --trees option)'
    
if settings['subsample'] is None:
    subsample = [-1]*ntrees
else:
    subsample = [int(s) for s in settings['subsample']]
    nsubsample = len(subsample)
    assert nsubsample == ntrees, 'Expecting %d subsample settings but only %d were supplied' % (ntrees, nsubsample)
    
if settings['subseed'] is None:
    subseed = [1]*ntrees
else:
    subseed = [int(s) for s in settings['subseed']]
    nsubseed = len(subseed)
    assert nsubseed == ntrees, 'Expecting %d subseed settings but only %d were supplied' % (ntrees, nsubseed)
    
if settings['fill'] is None:
    fill = ["#999999"]*(ntrees+1)
else:
    fill = ['"%s"' % s for s in settings['fill']]
    nfill = len(fill)
    assert nfill == ntrees + 1, 'Expecting %d fill settings but only %d were supplied' % (ntrees + 1, nfill)
    
if settings['stroke'] is None:
    stroke = ["#999999"]*(ntrees+1)
else:
    stroke = ['"%s"' % s for s in settings['stroke']]
    nstroke = len(fill)
    assert nstroke == ntrees + 1, 'Expecting %d stroke settings but only %d were supplied' % (ntrees + 1, nstroke)
    
if settings['ptsize'] is None:
    ptsize = [1.9]*(ntrees+1)
else:
    ptsize = [float(s) for s in settings['ptsize']]
    nptsize = len(fill)
    assert nptsize == ntrees + 1, 'Expecting %d ptsize settings but only %d were supplied' % (ntrees + 1, nptsize)
    
if settings['ptstroke'] is None:
    ptstroke = [0.1]*(ntrees+1)
else:
    ptstroke = [float(s) for s in settings['ptstroke']]
    nptstroke = len(fill)
    assert nptstroke == ntrees + 1, 'Expecting %d ptstroke settings but only %d were supplied' % (ntrees + 1, nptstroke)

def ensureFilePathExists(fpath):
    if not os.path.exists(fpath):
        sys.exit('Expecting this file path to exist: "%s"' % fpath)

def ensureFilePathDoesNotExist(fpath):
    if os.path.exists(fpath):
        sys.exit('Please delete or rename the file "%s" before running this program' % fpath)

def extractTLRadius(pct, fn):
    # fn should be the path to the R file containing OP Frechet mean results
    # For example, the first few lines of the file should look like this:
    # # (Sphaeroplea_robusta_UTEX2350:0.001186000,Ankyra_judai_SAG17_84:0.021105091,(((((((Tumidella_tumida_SAG2265:0.063690636,Dictyochloris_fragrans_UTEX127:0.054760970):0.000087384,(Planktosphaeria_gelatinosa_SAG262_1b:0.034971394,Bracteamorpha_trainori_CC1VF5A:0.049583848):0.000042047):0.000004486,(Bracteacoccus_minor_UTEX66:0.018045030,Bracteacoccus_aerius_UTEX1250:0.018885212):0.020902727):0.000002165,((Radiococcus_polycoccus_SAG217_1c:0.040317939,(Planktosphaeria_botryoides_UTEXLB951:0.009633788,Planktosphaeria_texensis_UTEX1241:0.012156727):0.042179667):0.020826212,Schizochlamys_gelatinosa_SAG66_94:0.043422182):0.019274879):0.000006257,((((Pseudomuriella_schumacherensis_SAG2137:0.016491667,Pseudomuriella_engadinensis_UTEX58:0.013051546):0.030039000,(Mychonastes_homosphaera_CAUPH6502:0.011134576,Mychonastes_jurisii_SAG3798:0.011493697):0.083522576):0.000019037,(Rotundella_rotunda_WJT54VFNP11:0.011019758,Rotundella_sp_ZNP1VF31:0.019132515):0.074008455):0.000007615,Chromochloris_zofingiensis_UTEX56:0.027395091):0.000000614):0.000043396,((((Kirchneriella_aperta_SAG2004:0.020551061,Ankistrodesmus_falcatus_UTEX101:0.023059485):0.013917000,Ourococcus_multisporus:0.036963333):0.021234545,Dictyococcus_varians_UTEXLB62:0.105211637):0.027416613,(((Hariotina_reticulata_UTEX1365:0.032551121,Scenedesmus_rotundus_SEV3VF49:0.017471757):0.007405928,Scenedesmus_obliquus_UTEX393:0.017483576):0.006359791,(((Parapediastrum_biradiatum_UTEX37:0.016587061,Pediastrum_duplex_UTEX1364:0.014214394):0.015451394,(Neochloris_aquatica_UTEX138:0.020831242,(Characiopodium_hindakii:0.011136758,Botryococcus_sudeticus:0.008455455):0.007515939):0.031896334):0.001231364,Chlorotetraedron_incus_SAG4381:0.010722970):0.014301394):0.001061503):0.000092667):0.000148220,Schroederia_setigera_UTEXLB2454:0.051171969):0.038260333)
    # # sample size = 10000
    # # variance = 0.006687729
    # # tree length = 1.341149028
    # # iterations = 32
    # # frechet-e = 0.01
    # # frechet-k = 1000000
    # # frechet-n = 10
    # # pseudorandom number seed = 1
    # # q25 = -0.079683907
    # # q75 = 0.079683907
    # # IQR = 0.159367815
    # # KDE sigma = 0.081776490
    # # KDE bandwidth = 0.010154650
    # # 95% HPD lower = 0.058685569
    # # 95% HPD upper = 0.102040726
    # # 95% radius = 0.101098573
    # # smallest distance = 0.051341071
    # # largest distance = 0.144040701
    ensureFilePathExists(fn)
    stuff = open(fn, 'r').read()
    
    # Extract TL
    m = re.search(r'# tree length = ([.0-9]+)', stuff)
    assert m is not None
    TL = float(m.group(1))

    # Extract radius
    m = re.search(r'# %d%% radius = ([.0-9]+)' % pct, stuff)
    assert m is not None
    radius = float(m.group(1))
    
    # Extract samplesize
    m = re.search(r'# sample size = ([0-9]+)', stuff)
    assert m is not None
    samplesize = float(m.group(1))
        
    return {'samplesize': samplesize, 'TL': TL, 'radius':radius}
    
outfn = None
outf = None
def output(s = None):
    global outf
    if s is None:
        print()
        outf.write('\n')
    else:
        print(s)
        outf.write('%s\n' % s)

if __name__ == '__main__':
    # Create the specified results directory if it does not already exist
    #if not os.path.exists(dirname):
    #    os.mkdirs(dirname, exist_ok=False)
    resultsdir = os.path.join(dirname, prefix)
    if not os.path.exists(resultsdir):
        os.makedirs(resultsdir, exist_ok=False)
        
    # Ensure that an analysis with this prefix has not already been run
    outfn = os.path.join(dirname, prefix, 'output.txt')
    ensureFilePathDoesNotExist(outfn)

    # Open the output file
    outf = open(outfn, 'w')
    
    ######################################################
    ## Calculate mean tree for each supplied treefile   ##
    ## saving the subsampled trees for later use in MDS ##
    ######################################################
    TLs = []
    radii = []
    meantrees = []
    for i,treefn in enumerate(trees):
        t = i + 1

        # Creating the op.conf file to compute the mean tree...
        confn = os.path.join(resultsdir, 'op.conf')
        conf = open(confn, 'w')
        conf.write('# Configuration file for op version 1.8\n')
        conf.write('# https://github.com/plewis/op/\n')
        conf.write('hpdrad = %s\n' % (hpdrad and 'yes' or 'no'))
        conf.write('radpct = %d\n' % radpct)
        conf.write('savetrees = yes\n')
        conf.write('prefix = op-mean-%d\n' % t)
        conf.write('frechetmean = yes\n')
        treepath = os.path.join(basedir, treefn)
        conf.write('treefile = %s\n' % treepath)
        conf.write('skip = %d\n' % skip)
        conf.write('subsample = %d\n' % subsample[i])
        conf.write('subseed = %d\n' % subseed[i])
        conf.write('scale = 1.0\n')
        #conf.write('rooted = no\n')
        conf.close()
        
        # Running OP to compute the mean tree...
        if runOP:
            print('Finding mean for treefile %d...' % t)
            completed_process = subprocess.run([oppath], cwd=resultsdir, capture_output=True)
            try:
                completed_process.check_returncode()
            except subprocess.CalledProcessError:
                output('\nAn error occurred while computing mean tree for treefile %d' % t)
                output('\nStandard error:')
                output('%s' % completed_process.stderr.decode('utf-8').strip())
                output('\nStandard output:')
                output('%s' % completed_process.stdout.decode('utf-8').strip())
                sys.exit('\nAborted.')
        
        outf.write('\nStatistics for treefile %d...\n' % t)
        stats = extractTLRadius(radpct, os.path.join(resultsdir, 'op-mean-%d.R' % t))        
        outf.write('  %12d sample size\n' % stats['samplesize'])
        outf.write('  %12.9f tree length\n' % stats['TL'])
        outf.write('  %12.9f radius\n' % stats['radius'])
        TLs.append(stats['TL'])
        radii.append(stats['radius'])
        meantrees.append(os.path.join(resultsdir, 'op-mean-%d.tre' % t))
        
    # Determine scaling factors (scale all treefiles to have mean TL 1.0)
    scalers = []
    for tl in TLs:
        assert tl is not None, 'tl was None'
        if doscale:
            scalers.append(1.0/tl)
        else:
            scalers.append(1.0)

    print('\nScalers:')            
    for s in scalers:
        print('  %g' % s)
        
    #################################################
    ## Calculate mean tree for the combined sample ##
    #################################################
    # Create the op.conf file
    # Note: we do not include --hpdrad option here even if hpdrad = True. This is because
    # if OP sees the --hpdrad option it will try to read the treefiles as RevBayes
    # treefiles (because hpdrad requires log posterior kernel values). This, in turn,
    # is bad because we are reading in treess from nexus files saved previously when
    # the mean was determined.
    confn = os.path.join(resultsdir, 'op.conf')
    conf = open(confn, 'w')
    conf.write('# Configuration file for op version 1.8\n')
    conf.write('# https://github.com/plewis/op/\n')
    conf.write('radpct = 95\n')
    conf.write('savetrees = yes\n')
    conf.write('prefix = op-mean-0\n')
    conf.write('frechetmean = yes\n')
    for i in range(len(trees)):
        t = i + 1
        treepath = 'op-mean-%d-saved.tre' % t
        conf.write('treefile = %s\n' % treepath)
        conf.write('scale = %g\n' % scalers[i])
        #conf.write('rooted = no\n')
    conf.close()

    # Running OP to compute the mean tree for the combined sample...
    if runOP:
        print('\nFinding mean for combined treefile')
        completed_process = subprocess.run([oppath], cwd=resultsdir, capture_output=True)
        try:
            completed_process.check_returncode()
        except subprocess.CalledProcessError:
            output('\nAn error occurred while calculating mean for the combined sample')
            output('\nStandard error:')
            output('%s' % completed_process.stderr.decode('utf-8').strip())
            output('\nStandard output:')
            output('%s' % completed_process.stdout.decode('utf-8').strip())
            sys.exit('\nAborted.')
    
    outf.write('\nStatistics for combined treefile...\n')
    stats = extractTLRadius(95, os.path.join(resultsdir, 'op-mean-0.R'))
    outf.write('  %12d sample size\n' % stats['samplesize'])
    outf.write('  %12.9f tree length\n' % stats['TL'])
    outf.write('  %12.9f radius\n' % stats['radius'])
    overall_mean_tree = 'op-mean-0.tre'
    
    ####################################################
    ## Calculate pairwise distances between all trees ##
    ####################################################
    # Create the op.conf file
    confn = os.path.join(resultsdir, 'op.conf')
    conf = open(confn, 'w')
    conf.write('# Configuration file for op version 1.8\n')
    conf.write('# https://github.com/plewis/op/\n')
    conf.write('prefix = op-pairwise\n')
    conf.write('pairwise = yes\n')
    conf.write('treefile = %s\n' % overall_mean_tree)
    conf.write('scale = 1.0\n')
    #conf.write('rooted = no\n')
    for i in range(len(trees)):
        t = i + 1
        # Use saved treefiles containing only subsampled trees (not the original treefiles)
        treepath = 'op-mean-%d-saved.tre' % t
        conf.write('treefile = %s\n' % treepath)
        conf.write('scale = %g\n' % scalers[i])
        #conf.write('rooted = no\n')
    conf.close()

    # Running OP to compute all pairwise distances...
    if runOP:
        print('Computing pairwise distances in combined treefile')
        completed_process = subprocess.run([oppath], cwd=resultsdir, capture_output=True)
        try:
            completed_process.check_returncode()
        except subprocess.CalledProcessError:
            output('\nAn error occurred while calculating pairwise distances')
            output('\nStandard error:')
            output('%s' % completed_process.stderr.decode('utf-8').strip())
            output('\nStandard output:')
            output('%s' % completed_process.stdout.decode('utf-8').strip())
            sys.exit('\nAborted.')
    
    # Create an R file to carry out the multidimensional scaling
    samplesizes = [ss == -1 and 0 or ss for ss in subsample]
    
    # Renders the red (mean) dot last, ensuring it is visible
    # Note that this also requires the conf2 data frame to have its rows reversed
    fills     = ['rep(%s, %d)' % (f,ss) for (f,ss) in zip(fill[1:],samplesizes)]     + ['%s' % fill[0]]
    strokes   = ['rep(%s, %d)' % (s,ss) for (s,ss) in zip(stroke[1:],samplesizes)]   + ['%s' % stroke[0]]
    ptsizes   = ['rep(%g, %d)' % (s,ss) for (s,ss) in zip(ptsize[1:],samplesizes)]   + ['%g' % ptsize[0]]
    ptstrokes = ['rep(%g, %d)' % (s,ss) for (s,ss) in zip(ptstroke[1:],samplesizes)] + ['%g' % ptstroke[0]]
    
    rfile = open(os.path.join(resultsdir,'mds.R'), 'w')
    
    rfile.write('# Load libraries\n')
    rfile.write('library(smacof)\n')
    rfile.write('library(ggplot2)\n')
    rfile.write('library(plotly)\n')
    rfile.write('library(htmlwidgets)\n')
    rfile.write('source("./op-pairwise.R")\n')
    
    rfile.write('\n# Run 2d MDS\n')
    rfile.write('weights <- 1 / (d^2)               # larger distances downweighted more\n')
    rfile.write('weights[is.infinite(weights)] <- 0 # Set infinite weights to 0\n')
    rfile.write('weights[is.na(weights)] <- 0       # Handle any NA values, e.g. for missing data\n')
    rfile.write('mds2 <- smacofSym(delta = d, weightmat = weights, ndim = 2)\n')
    
    rfile.write('\n# Uncomment the line below to create Shepard diagram\n')
    rfile.write('# plot(mds2, plot.type = "Shepard", main = "Shepard Diagram (Ratio Transformation)")\n')
    
    rfile.write('\n# Reverse the rows in the data frame conf2 so that the dot\n')
    rfile.write('# representing the mean comes last (ensuring it will be visible)\n')
    rfile.write('conf2 <- as.data.frame(mds2$conf)\n')
    rfile.write('conf2.reversed <- conf2[seq(nrow(conf2),1),]\n')

    rfile.write('\n# Specify fills, strokes, point sizes, and point strokes\n')
    rfile.write('# (keeping in mind that the mean dot comes last)\n')
    rfile.write('fills <- c(%s)\n' % ','.join(fills))
    rfile.write('strokes <- c(%s)\n' % ','.join(strokes))
    rfile.write('pointsizes <- c(%s)\n' % ','.join(ptsizes))
    rfile.write('pointstrokes <- c(%s)\n' % ','.join(ptstrokes))

    rfile.write('\n# Plot the points in 2-d space using ggplot2.\n')
    rfile.write('p <- ggplot(conf2.reversed, aes(x = D1, y = D2))\n')
    rfile.write('p <- p + geom_point(shape = 21, size = pointsizes, stroke = pointstrokes, color = strokes, fill = fills) + coord_fixed(ratio = 1)\n')
    rfile.write('#+ ggtitle("rbcL Posterior Sample")\n')
    
    rfile.write('\n# Save the plot to a file for inclusion in the manuscript.\n') 
    rfile.write('ggsave("mds-plot.png", p, width = 4, height = 2, units = "in", dpi = 300)\n')

    rfile.close()    
    
    # Running mds.R to carry out the multidimensional scaling
    subprocess.run(['rscript', 'mds.R'], cwd=resultsdir)

    # Close the output file
    outf.close()
```
