## Supplementary material for "Estimating Bayesian Phylogenetic Information Content Using Geodesic Distances": hpdstd.R

```
library(MASS)     # for mvrnorm
library(mvtnorm)  # for dmvnorm
library(ggplot2)  # for ggplot

# Open a file for the plot
cwd = system('cd "$( dirname "$0" )" && pwd', intern = TRUE)
setwd(cwd)
pdf("fig-hpdstd.pdf")

# circleFun from joran answer at https://stackoverflow.com/questions/6862742/draw-a-circle-with-ggplot2
# generates data for a circle with a specified center and radius
circleFun <- function(center = c(0,0), radius = 1, npoints = 100) {
    tt <- seq(0,2*pi,length.out = npoints)
    xx <- center[1] + radius * cos(tt)
    yy <- center[2] + radius * sin(tt)
    return(data.frame(x = xx, y = yy))
}

# lineFun generates data for a line with specified origin, radius, and angle (theta)
lineFun <- function(origin = c(mux, muy), radius = stdrad, theta = 0.0, npoints=100) {
    tt <- seq(0, radius, length.out = npoints)
    xx <- origin[1] + tt * cos(theta)
    yy <- origin[2] + tt * sin(theta)
    return(data.frame(x = xx, y = yy))
}

# Generate ten thousand points from correlated bivariate normal
true.mu <- c(0,0)
true.sigma <- matrix(c(1, .95, .95, 1),nrow=2)
n <- 10000
data <- mvrnorm(n, true.mu, true.sigma)
x <- data[,1]
y <- data[,2]

# Determine the sample means
mux <- mean(x)
muy <- mean(y)

# Find distances from mean to each point
dist <- sqrt((x - mux)^2 + (y - muy)^2)

# Find the radius enclosing 95% of points
stdrad <- quantile(dist, 0.95)
stdrad

# Compute probability density of each point
dens <- dmvnorm(data, true.mu, true.sigma)

# ggplot2 expects data to be in a data frame
df <- data.frame(x, y, dens, dist)

# Sort df so that highest density points are first
df.sorted <- df[order(df$dens, decreasing = TRUE), ]

# Find cutoff such that 95% of points have density higher than cutoff
cutoff <- df.sorted[9500, "dens"]
cutoff

# create bool array indicating which points are in the HPD credible set
hpd <- df.sorted$dens >= cutoff
count <- sum(hpd == TRUE)
count

# Add hpd as a column in df.sorted
df.sorted$in_hpd <- factor(hpd, levels=c(TRUE, FALSE), labels=c("yes", "no"))

# Subset df.sorted including only points in which hpd is TRUE
hpd.subset <- df.sorted[df.sorted$dens >= cutoff, ]
hpdrad <- max(hpd.subset$dist)
hpdrad

# Generate data for circle of radius stdrad
stdrad.circle <- circleFun(center = c(mux, muy), radius=stdrad, npoints=100)

# Generate data for circle of radius hpdrad
hpdrad.circle <- circleFun(center = c(mux, muy), radius=hpdrad, npoints=100)

# Generate data for line of length stdrad
stdrad.line <- lineFun(origin = c(mux, muy), radius = stdrad, theta = 0.0, npoints=100)

# Generate data for line of length hpdrad
hpdrad.line <- lineFun(origin = c(mux, muy), radius = hpdrad, theta = pi/8, npoints=100)

# Run ggplot2
ggplot(data = df.sorted, aes(x = x, y = y, color = in_hpd)) +
  scale_color_manual(values = c("navy", "gray")) +
  geom_point(size=0.5) +
  geom_path(data=stdrad.circle, colour="darkorchid", linewidth=1) +
  geom_path(data=stdrad.line, color="darkorchid", linewidth=1) +
  geom_path(data=hpdrad.circle, colour="darkorange", linewidth=1) +
  geom_path(data=hpdrad.line, color="darkorange", linewidth=1) +
  labs(title = "", x = "Dimension 1", y = "Dimension 2") + 
  theme_minimal() +
  theme(legend.position = "none", aspect.ratio = 1)

# Close the file
dev.off()
```
