## Supplementary material for "Estimating Bayesian Phylogenetic Information Content Using Geodesic Distances": supplementary-figures.pdf

### Radius based on alternative measures of dispersion

If the distribution of tree distances from the mean tree is not symmetric — that is, has ridges of high density — then the method for determining the 95% radius described in the paper (the RAD method) will omit some points that lie within the highest probability density (HPD) region and will include some points that have relatively low posterior probability density. An admittedly imperfect analogy is provided by a sample of 10,000 points from a correlated bivariate normal distribution (Fig. S1). The circle having the reference radius is shown in purple. The 9,500 points falling within the HPD region are navy-colored, with those outside the HPD region in gray. Clearly the reference radius omits some points in the HPD region. An alternative is to create a circle just large enough to enclose all points within the HPD set (the HPD method). The HPD radius and reference radius would be identical if the covariances were all zero; it is the asymmetry of the distribution that leads to the HPD radius being larger than the reference radius.

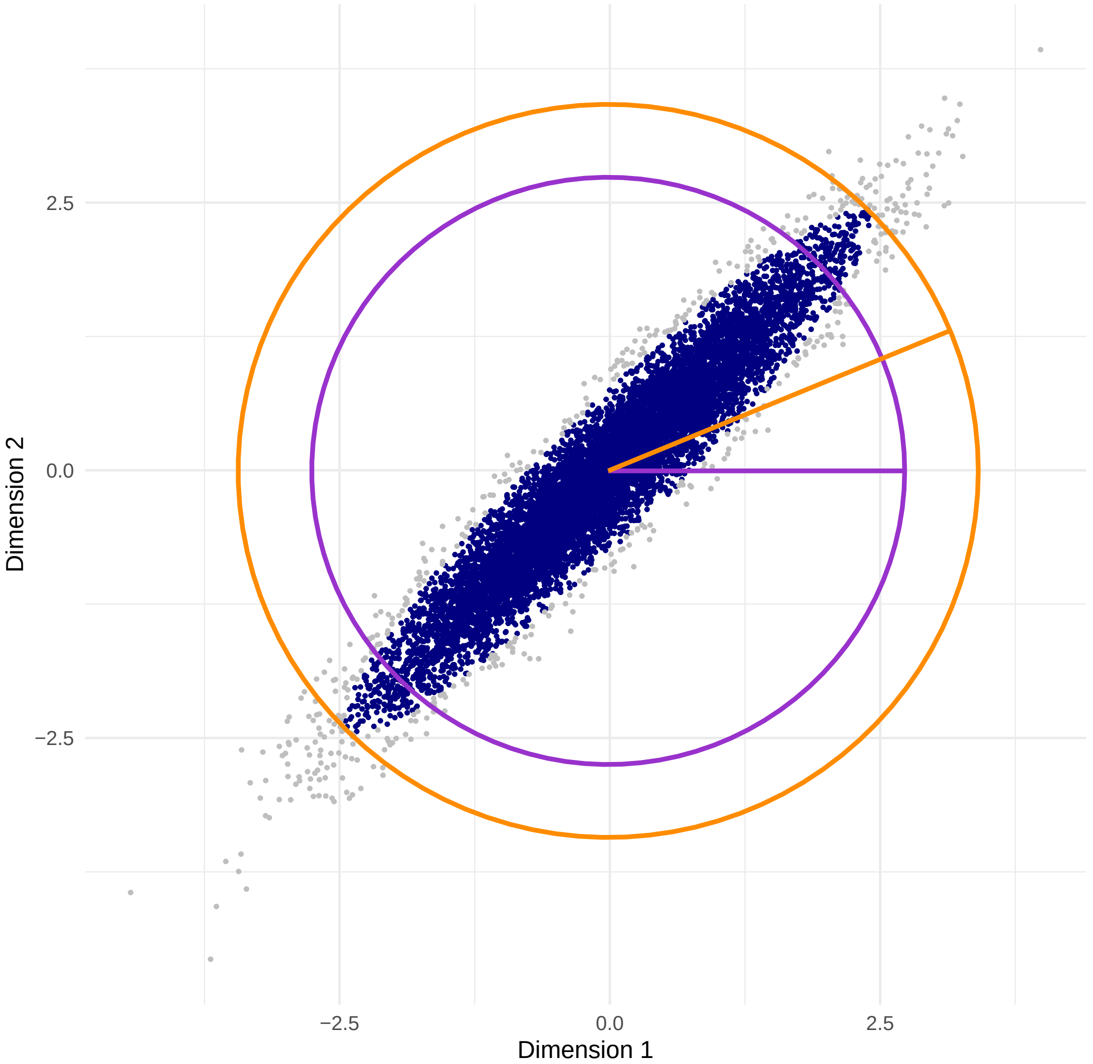


Figure S1: Scatterplot showing points sampled from a bivariate normal distribution with mean (0.0,0.0), variances each 1.0, and covariances each 0.95. Points within the 95% highest probability density (HPD) ellipse are navy; points outside are gray. Purple circle encloses 95% of points closest to the sample mean. Orange circle encloses all points within the 95% HPD set.

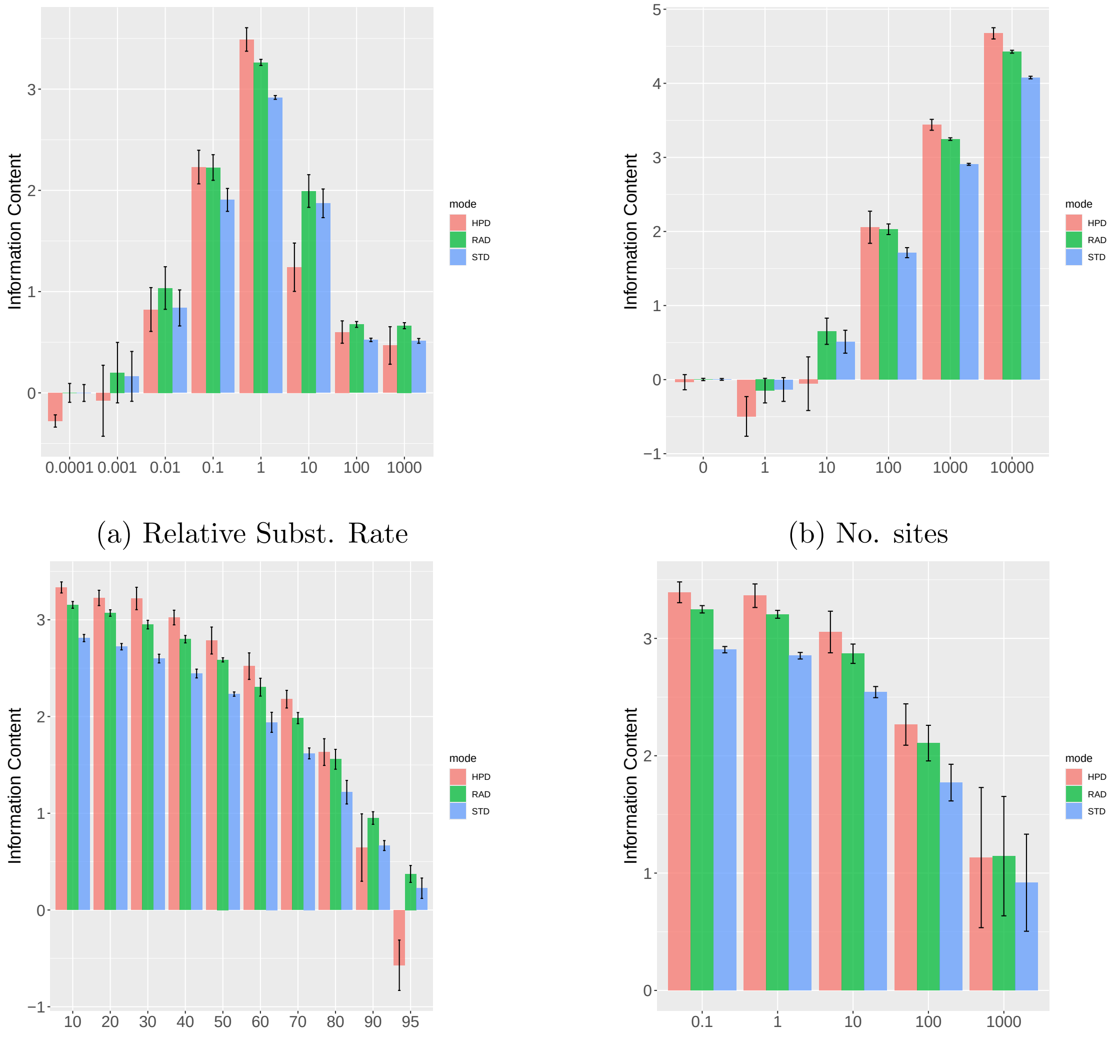


(a) Relative Subst. Rate

(b) No. sites

(c) Missing %

(d) ASRV

Figure S2: Information content ($LCR$) for 5-taxon datasets using the 95% radius described in the paper (RAD) as well as the 95% HPD radius (HPD) and the standard deviation (STD) as the “volume” in the LCR formula. See the caption of the corresponding figure in the main paper for other details.

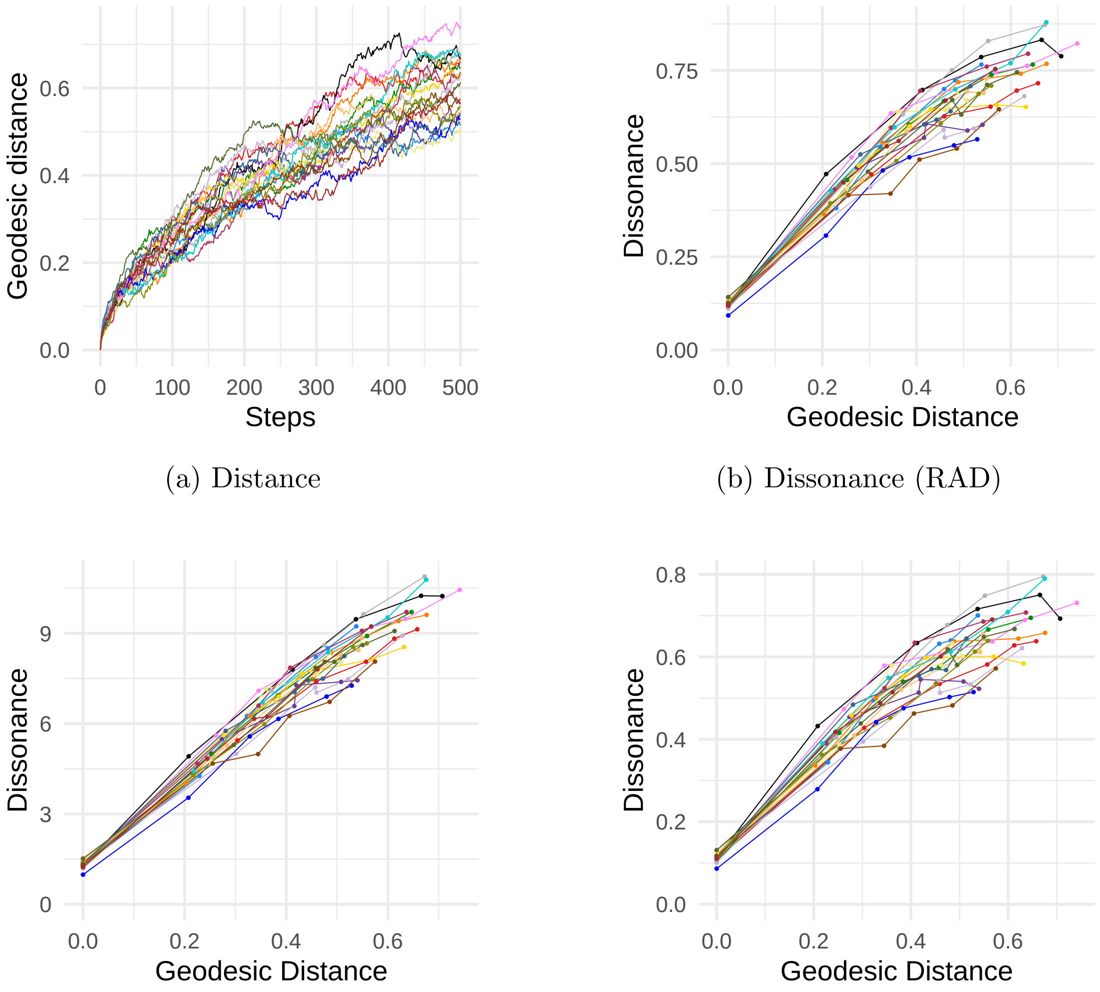


(a) Distance

(b) Dissonance (RAD)

(c) Dissonance (STD)

(d) Dissonance (HPD)

Figure S3: (a) Geodesic distance accumulated during random walk. (b) Dissonance using 95% radius (RAD method). (c) Dissonance using standard deviation (STD method). (d) Dissonance using 95% HPD radius (HPD method)
