## Supplementary figures and images for "Estimating Bayesian Phylogenetic Information Content Using Geodesic Distances"

### fig-hpdstd.pdf

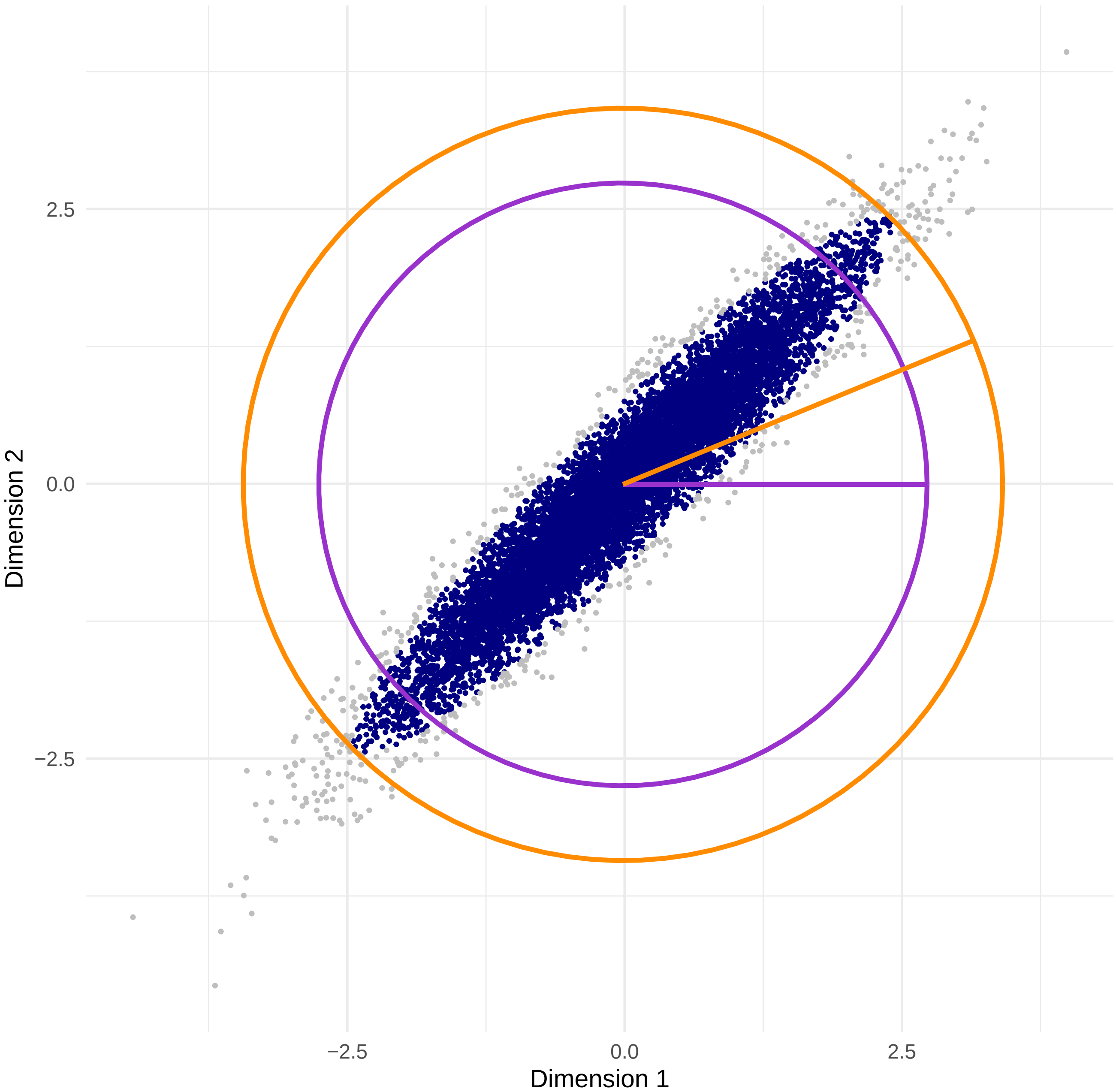
